# Cell type-specific coexpression networks funnel genetic risk into core genes

**DOI:** 10.64898/2026.11.09.764265

**Authors:** Xinpei Wang, Sriram Sankararaman, Xuanyao Liu, Minhui Chen, Andy Dahl

## Abstract

Genetic effects on common disease are mediated through many genes in multiple cell types, yet how networks connect genome-wide genetic risk to core disease genes remains unclear. One key limitation is that disease-related gene networks are likely cell type-specific, but these are poorly captured by bulk and pseudobulk coexpression networks. Here we used population-scale single-cell immune transcriptomes to decompose interindividual gene coexpression into cell type-shared and -specific networks, which have nearly opposite properties. Shared network hubs were enriched in constrained housekeeping genes, whereas cell type-specific hubs were dynamically expressed, rapidly evolving, and enriched in cell type-matched enhancers and transcription factors. Specific hubs enriched for immune disease GWAS heritability, Mendelian genes, and drug targets; these enrichments were absent or weak for non-immune disease and shared hubs. Gene-gene network edges enriched for causal *trans* regulation identified by *trans* eQTLs and perturb-seq, with shared hubs skewed to upstream regulators and specific hubs skewed to downstream disease-proximal genes. Specific hubs increased the relative success of genetically supported immune drug targets from 1.7-fold to 2.7-fold; for example, the lupus drug target *BAFF* is a monocyte hub connected to the Mendelian gene *TREX1* and interferon-responsive genes. Overall, accounting for shared housekeeping networks reveals that cell type-specific networks funnel genetic risk into downstream and dynamic core disease genes.

## Introduction

Common diseases arise from the coordinated activity of many genes in multiple cell types. Genetic effects on gene expression (eQTLs) provide a lens into these processes, but current approaches only explain a small minority of disease-related gene regulation (Mostafavi et al. 2023; Connally et al. 2022; Umans et al. 2021; Yao et al. 2020). One key limitation is that most known gene regulatory effects act in *cis*, yet *trans* effects that propagate through gene networks explain substantially more variance (Liu et al. 2019; Price et al. 2011). Resolving gene coexpression networks across individuals is therefore essential for answering if and how polygenic *trans* regulatory signals converge into core genes that directly affect disease.

Network models of bulk tissue RNA-seq, such as WGCNA (Langfelder and Horvath 2008; Voineagu et al. 2011), can identify modules that capture convergence of genome-wide risk genes (Gordon et al. 2026; Wen et al. 2024) but are often dominated by housekeeping processes (Farahbod and Pavlidis 2020). *trans*-eQTLs can explain how particular risk variants propagate through networks to impact complex traits (Salamone et al. 2026; Võsa et al. 2021; Wang et al. 2024; Warmerdam et al. 2026; Wen et al. 2024), but they often have low power due to selective constraint, weak effects, and high multiple testing burden (Battle et al. 2013; Liu et al. 2019). Perturb-seq has emerged as a powerful approach to identify *trans* effects (Ota et al. 2026; Replogle et al. 2022; Zhu et al. 2026), but it usually remains limited to a few donors, a few (often transformed) cell types, and unrealistically large effect sizes. Overall, we lack an unbiased account of how genome-wide genetic risk converges into core disease genes (García-González and O’Reilly 2026). We hypothesize this gap arises because current coexpression networks primarily capture cell type-shared processes, but disease-related gene regulation is primarily cell type-specific (Minhui Chen et al. 2026; Krockenberger et al. 2026; Mitchel, Nazeen, et al. 2025; Gilad and Battle 2026).

Population-scale single-cell RNA-seq (scRNA-seq) (Nathan et al. 2022; Perez et al. 2022; Yazar et al. 2022) enables decomposing interindividual coexpression across cell types. However, prior networks model variation across cells rather than individuals (Aibar et al. 2017; Morabito et al. 2023; Su et al. 2023) and do not account for cell type-shared processes. Here, we developed a method called GECO to separate cell type-shared and -specific interindividual coexpression networks, which revealed that they are qualitatively distinct: shared networks dominate standard methods and capture essential housekeeping processes, while specific networks capture dynamic immune processes funneling *trans* regulatory effects into core immune disease genes.

## Results

### GECO identifies cell type-specific gene networks

Our two-step approach, GECO, is illustrated in **Fig 1**. First, GECO uses a linear mixed model to unbiasedly partition interindividual covariance between genes into components that are cell type-shared, *V*, or specific to cell type *c*, *Ω^c^* **(Methods)**. Second, GECO constructs cell type-shared and -specific gene coexpression networks using a new approach based on graphical lasso (glasso, Friedman, Hastie, and Tibshirani 2008), outputting sparse gene-gene networks **(Methods)**. We derived analytic solutions for GECO covariance estimates and standard errors, making it computationally efficient for large scRNA-seq datasets (**Note S1**).

**Figure 1:**
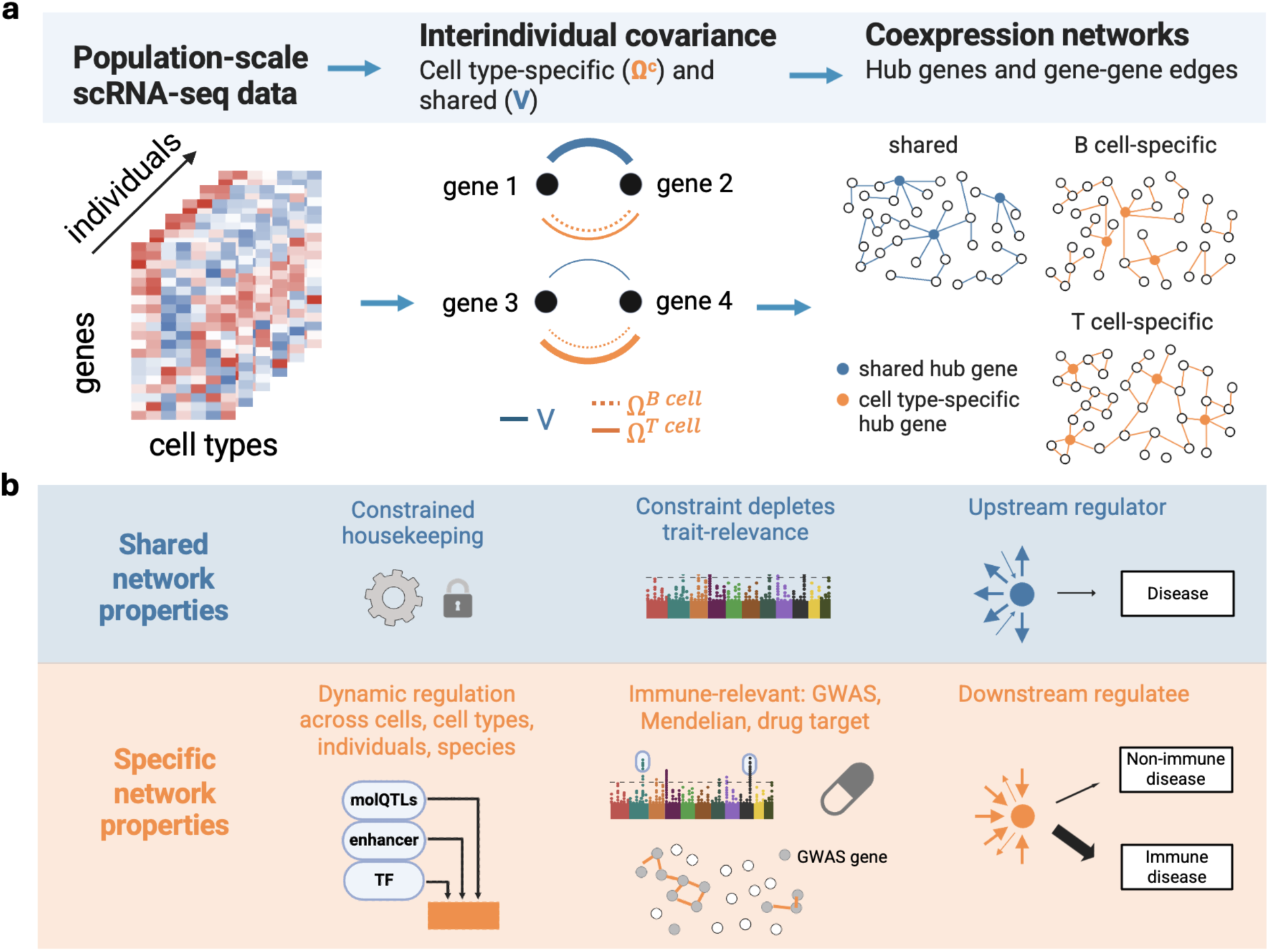
Overview of GECO and the distinct properties of shared and cell type-specific coexpression networks. **(a)** GECO unbiasedly partitions shared and specific gene coexpression matrices across individuals and then summarizes them into sparse partial correlation networks with a model based on graphical lasso. In the example, genes 1 and 2 have large shared covariance, genes 3 and 4 have large T cell-specific covariance, and both pairs have small B cell-specific covariance. **(b**) Shared (blue) and specific (orange) gene coexpression networks have qualitatively distinct biological properties and relationships to complex human traits.

We performed simulations based on OneK1K to compare GECO to an established pseudobulk method, WGCNA (Langfelder and Horvath 2008), and to standard glasso (**Methods**). GECO’s first step unbiasedly partitioned interindividual covariance into shared and specific components (**Fig 2a**). Conversely, bulk and pseudobulk covariance estimates mix together both components, as cell type-shared covariance remains present even within a single cell type. GECO’s second step accurately distinguished shared and specific networks in terms of gene-gene edges and hub genes (**Fig 2b**). Conversely, WGCNA primarily recovered shared hubs, even in pseudobulk (**Fig 2b, Ext Data Fig 1 a**), and often failed to identify any modules in low-abundance cell types (**Ext Data Fig 1 b**). We also applied standard glasso on bulk and pseudobulk, which had intermediate performance between GECO and WGCNA in recovering cell type-specific hubs and edges in pseudobulk (**Fig 2b**) but detected fewer edges overall (**Ext Data Fig 1 b**, **Fig S1**). These results are robust to different thresholds for hub genes (**Ext Data Fig 1 d**).

**Figure 2:**
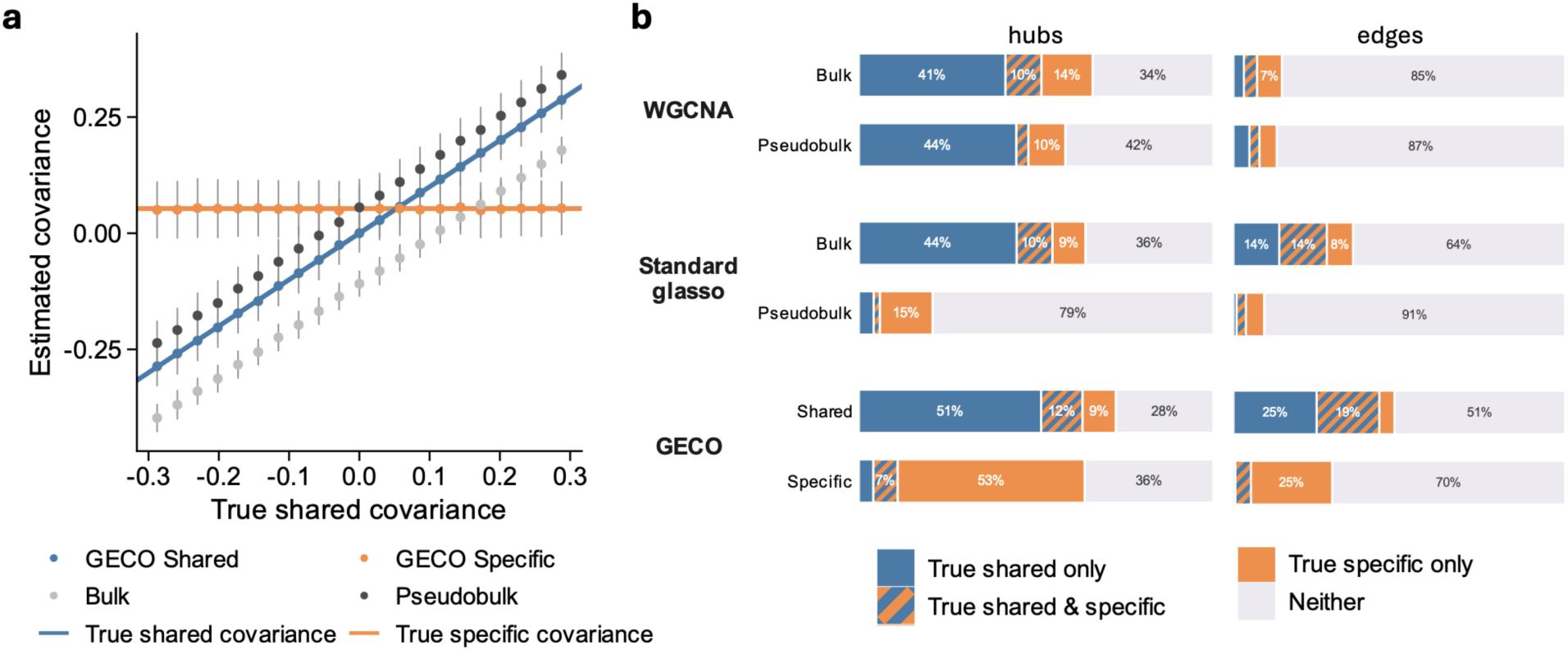
GECO partitions interindividual gene coexpression to fit shared and cell type-specific networks in simulations based on OneK1K data. **(a)** GECO unbiasedly fits shared and cell type-specific interindividual covariance, whereas bulk and pseudobulk fit a single covariance that reflects a mix of shared and specific. Points represent the mean estimates across 1,000 replicates, and error bars represent ±1 standard deviation. Bulk and pseudobulk estimates may be biased up or down depending on simulation parameters. **(b)** Composition of the top hubs and edges estimated by GECO and two comparators, WGCNA and standard glasso. We used the top 100 edges and chose the number of top hubs to match the true number of simulated hubs (when WGCNA and glasso networks were too sparse, we used all connected genes as hubs, **Ext Data** Fig 1b). The slashed orange/blue represent true hub genes in both shared and cell type-specific networks. In “Bulk” and “Shared”, “True specific” is the union of true cell type-specific hubs across all cell types; in “Pseudobulk” and “Specific”, “True specific” is the true cell type-specific hubs per cell type and averaged across cell types (**Ext Data** Fig 1a**,c** shows all cell types).

We applied GECO to 11,526 genes, N=928 individuals and the 7 most abundant cell types in OneK1K (**Methods**) to estimate a shared network and 7 cell type-specific networks. We defined hub genes as the 5% most-connected genes per network and used them as representative proxies since they spanned most connected genes (62-92%, **Fig S2a**). GECO shared hubs had low overlap with cell type-specific hubs (1.2-3.7× fold enriched; **Ext Data Fig 2 c**), whereas pseudobulk WGCNA hubs had substantial overlap with bulk WGCNA hubs (4.6-20.1×; **Ext Data Fig 2 c**) and GECO shared hubs (6.4-19.6×; **Ext Data Fig 2 d**). Standard glasso on pseudobulk fit very sparse networks leaving most genes isolated (**Ext Data Fig 2 a**). Further, WGCNA hubs were enriched in highly expressed genes (**Ext Data Fig 2 e)**, while standard glasso hubs were enriched in genes with zero expression in some individuals (**Ext Data Fig 2 f**). We conclude that GECO accurately recovers cell type-specific networks, unlike bulk and pseudobulk networks.

### Shared and specific hubs have distinct biology and evolutionary history

To get an overview of cell type-shared and -specific hubs, we evaluated simple benchmark gene sets (**Fig 3a**) and GO terms (**Fig 3b**). Shared hubs were enriched for housekeeping genes and broad RNA-related GO terms including mRNA processing and RNA splicing, whereas specific hubs were enriched for immune cell marker genes and immune-related GO terms including adaptive immune response, regulation of lymphocyte activation, and B cell activation. Conversely, WGCNA and standard glasso hubs were enriched in both housekeeping and immune marker genes (**Fig S6**) and had generic top GO terms, such as chromosome segregation (CD4ET hubs) and actin filament organization (BMem hubs, **Ext Data Fig 2 g,h**). We also found that shared hubs were stably expressed across cells within cell types, while specific hubs showed excess variability across cells within cell types as defined by the homeostasis Z-index (Mengjie Chen 2025) and dispersion (Jamison et al. 2026) (**Fig 3a**).

**Figure 3:**
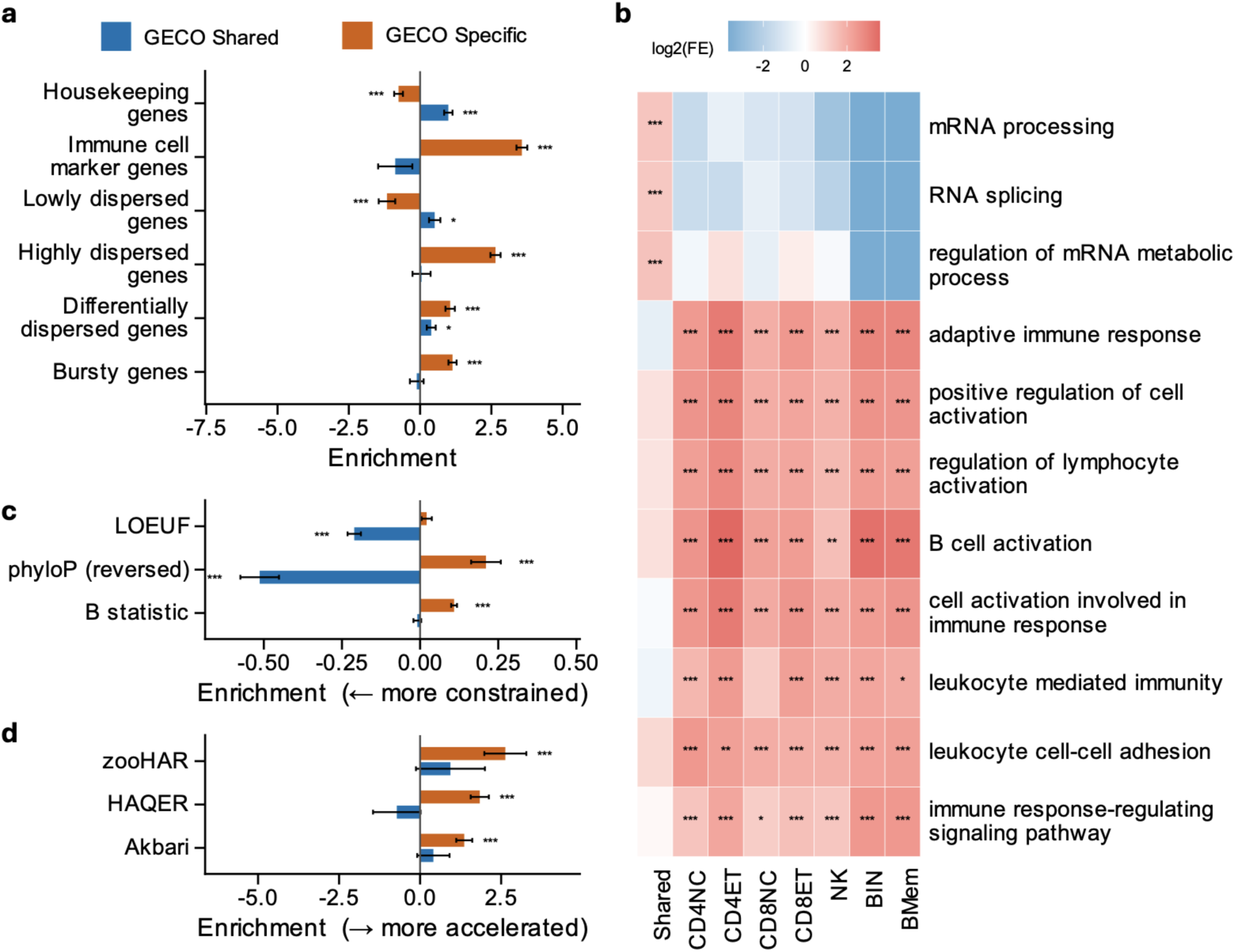
Shared network hubs are constrained housekeeping genes, while cell type-specific hubs are dynamic immune genes. **(a)** Benchmark gene sets show shared hubs are stable housekeeping genes, while specific hubs are dynamic within and between cell types (**Methods**). **(b)** The top 3 GO terms for shared hubs and each cell type’s specific hubs; shared GO terms implicate broad mRNA processes while specific GO terms implicate immune responses (further GO terms are in **Table S1**). **(c)** Shared hubs have greater evolutionary constraint based on LOEUF and phyloP (lower LOEUF and higher phyloP), while specific hubs have lower constraint based on phyloP and background selection (higher B statistic). **(d)** rE2G-linked regulatory regions for specific hubs enrich for annotations indicating directional selection and/or relaxed constraint. Sensitivity analyses are in **Fig S4**; analyses with WGCNA and standard glasso are in **Fig S6**. In **a**, **c**, and **d**, ***: p < 0.001, **: p < 0.01, *: p < 0.05; **b** uses FDR due to testing hundreds of GO terms: ***: FDR < 0.001, **: FDR < 0.01, *: FDR < 0.05. “Enrichment” x-axes represent log2 odds ratios in **a** and **d**, and linear regression coefficients in **c**. Error bars indicate +/- 1 standard error.

Shared and specific hub genes had distinct evolutionary features. Shared hubs had more constraint than average genes: depleted of loss-of-function variants (low LOEUF, S. Chen et al. 2024) and more conserved across species (high phyloP, Christmas et al. 2023, **Fig 3c**). Conversely, specific hubs were less constrained: they had low phyloP, lie in regions with lower background selection (higher B statistic, McVicker et al. 2009), and their enhancers (linked by rE2G (Gschwind et al. 2026)) were enriched in loci that evolved rapidly in the human lineage (zooHAR (Keough et al. 2023), HAQER (Mangan et al. 2022)) and are under recent directional selection (Akbari et al. 2026, **Fig 3d**). WGCNA and glasso hubs were generally constrained, like shared hubs (**Fig S6**).

These results were largely robust to adjustment for total expression, differential expression across cell types, gene length, and GC content (**Fig S4**), and to different hub thresholds (**Fig S5**). Together, they show that shared hubs are constrained housekeeping genes, while specific hubs are rapidly-evolving and dynamic immune genes.

### Shared and specific hubs capture distinct modes of gene regulation

Because specific hubs are dynamically regulated in immune cells, we asked how they relate to transcription factors (TFs) and functional genomic marks. Cell type-specific hubs were enriched for lineage-matched TFs from Human Protein Atlas (HPA, all p<1.4e-07, **Fig S7a**), while shared hubs were not. Further, GECO network degree predicted cell type lineage for HPA TFs, and shared networks predicted “low specificity” TFs (**Methods**, **Fig 4a**). WGCNA and glasso hubs showed much weaker TF enrichment, and glasso degree did not predict TF lineage (**Fig S8**).

**Figure 4:**
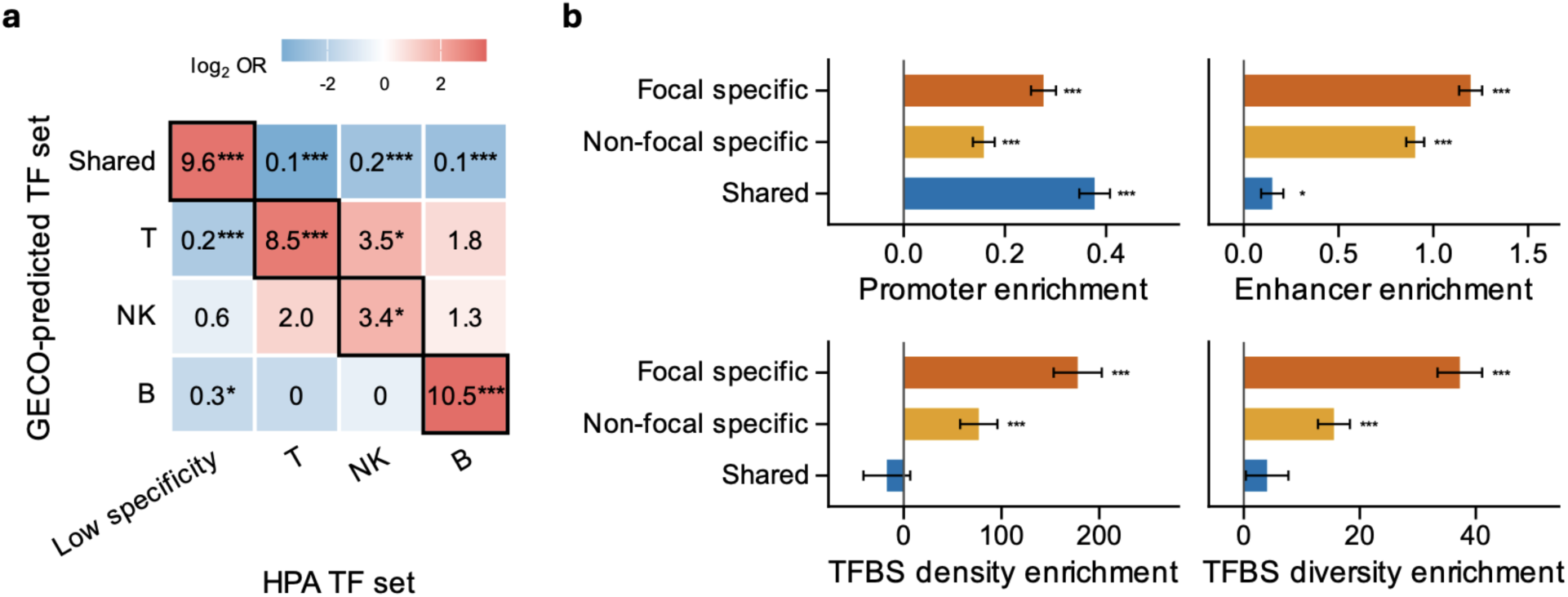
Cell type-specific networks capture cell type-specific gene regulation. **(a)** HPA TFs are most-connected in cell type-specific networks in appropriate lineages (T, NK, B) and, for ‘low specificity’ TFs, in the shared network. Sensitivity analyses are in **Fig S7b**. **(b)** GECO hub enrichment in functional genomic features in immune biosamples (FANTOM5 linked-promoter count, ENCODE-rE2G linked-enhancer count, and TF-TRACE TFBS density and diversity). Specific enrichments were calculated separately for lineage-matched (focal) and non-lineage-matched (non-focal) immune biosamples. Sensitivity analyses shown in **Fig S9**. Error bars indicate +/- 1 standard error from gene-level bootstrap. In **b**, x-axes represent regression coefficients. ***: p < 0.001; **: p < 0.01; *: p < 0.05.

We next assessed functional genomic features of hubs. Promoters were enriched in shared and specific hubs, with promoters from cell type-matched biosamples increasing enrichment in specific hubs (**Fig 4b**). In contrast, rE2G-linked enhancers, TFBS density, and TFBS diversity were strongly enriched in cell type-specific hubs, especially in matched biosamples, but not in shared hubs (**Fig 4b**). These results are consistent with expectations that promoters mostly have shared effects while enhancers are mostly context-specific (Kundaje et al. 2015).

### Specific hubs mediate gene regulation in immune disease

We hypothesized that genetic variation in cell type-specific hubs would explain more phenotypic variance than shared hubs due to their rapid evolution and dynamic regulatory features. We first examined *cis* genetic effects on molecular phenotypes (molQTLs) and found that shared hubs were broadly depleted, whereas cell type-specific hubs were broadly enriched: this held for *cis*-eGenes, *cis*-pGenes, *cis*-meGenes and *cis*-caGenes in diverse bulk and pseudobulk datasets (**Figs 5a**, **S10**). We next evaluated context-specific eQTLs and found that specific hubs were enriched in cell type- and tissue-specific eQTLs (from CLUES and GTEx, respectively) and depleted in tissue-shared eQTLs (**Fig 5a**). Generally, WGCNA and glasso hubs were depleted in *cis*-molGenes, similar to shared hubs (**Fig S11**). We found similar results based on heritability explained by *cis* eQTLs (*cis* h2), which eliminates bias from eQTL detection power: specific hubs had slightly higher bulk tissue *cis* h2 (Ouwens et al. 2020; Price et al. 2011; The GTEx

**Figure 5:**
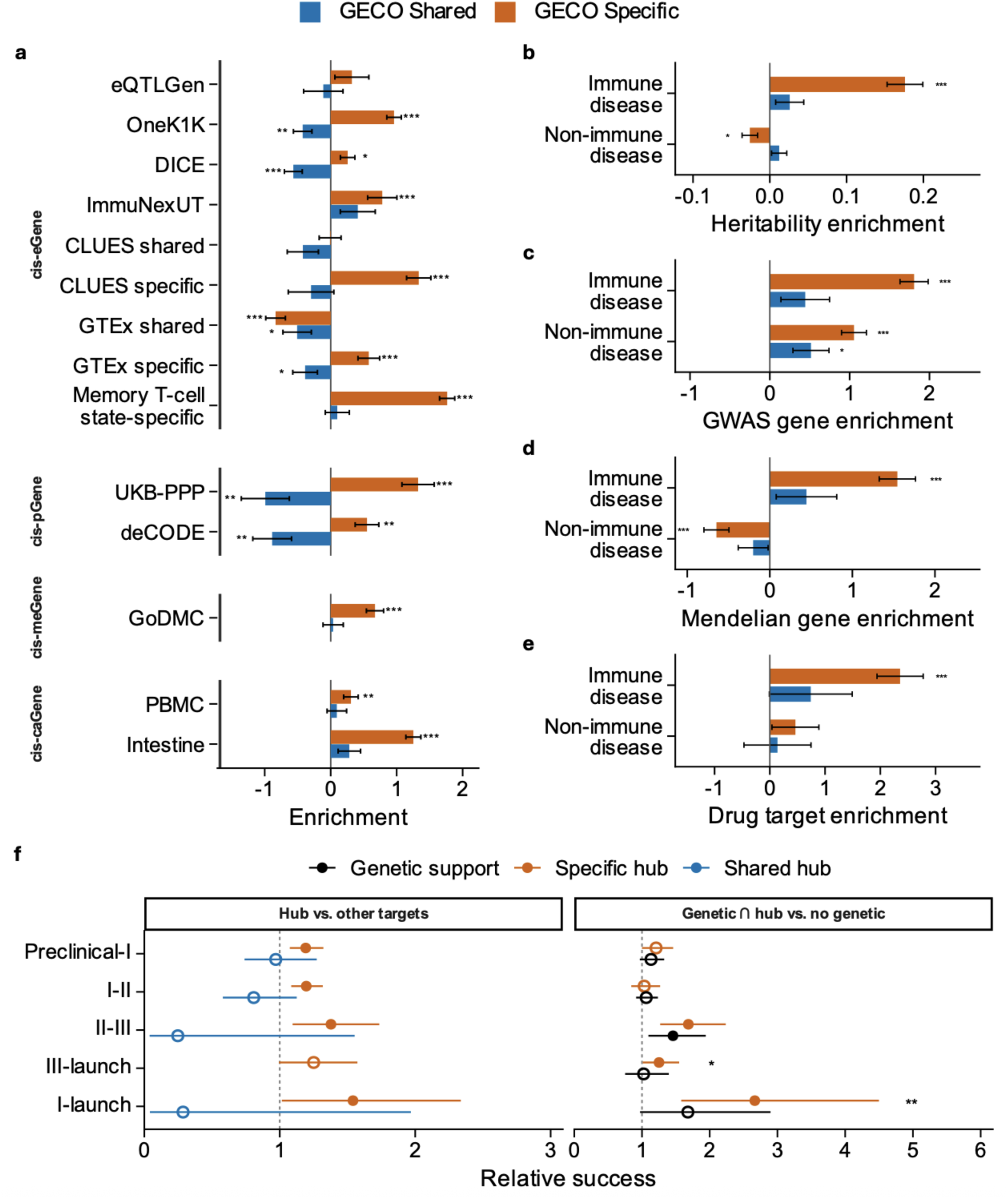
Cell type-specific hubs, but not shared hubs, are enriched in genetic effects on *cis* molGenes and complex traits. **(a)** Cell type-specific hubs were enriched in *cis*-molecular QTL genes, including single-cell eGenes (Memory T cell state-dependent, CLUES, OneK1K), bulk eGenes in sorted immune cells (DICE, ImmuNexUT), cell type-specific eGenes (CLUES), tissue-specific eGenes (GTEx), plasma pGenes based on Olink (UKB-PPP) and SomaScan (deCODE), meGenes from bulk blood (GoDMC), and single-cell caGenes (from PBMC and intestine). eQTLGen has low power since 94.8% of GECO genes are eGenes. **(b-e)** Cell type-specific hubs showed strong enrichment in immune diseases based on S-LDSC heritability (τ*, **b**), L2G-prioritized GWAS genes (**c**), Mendelian genes (**d**), and drug targets (**e**). Sensitivity analyses are shown in **Figs S12-13.** In **a, c, d,** and **e**, x-axes show log2 odds ratios. Error bars indicate +/- 1 standard error. **(f)** Relative success of target-indication pairs in immune indications (MeSH C20) from Citeline Pharmaprojects. Rows show progression of drug targets through phases. Panels show relative success for shared and specific hub genes (left) and for the intersection of specific hubs and indication-matched genetically supported genes as in Minikel et al. (OMIM or GWAS, right). These analyses restrict to genes included in GECO. Bars show 95% CI; filled points significantly differ from 1; and stars compare the orange and black estimates. ***: p < 0.001; **: p < 0.01; *: p < 0.05.

Consortium 2020; Wheeler et al. 2016; Wright et al. 2014, **Ext Data Fig 3 a**) and cell type-specific *cis* h2 (Minhui Chen et al. 2026, **Ext Data Fig 3 b**), yet had lower cell type-shared *cis* h2; shared hubs generally had lower *cis* h2.

We next assessed genetic effects on immune and non-immune diseases. First, we tested heritability enrichment in 117 FinnGen GWAS with stratified LD score regression (LDSC, **Methods**, Finucane et al. 2015). Cell type-specific hubs were enriched for immune disease heritability and slightly depleted for non-immune disease heritability, while shared hubs had no enrichments (**Fig 5b**). These results were robust to baseline annotations, unlike bulk network hubs (Kim et al. 2019), to modifying the gene window size, the number of hub genes, the h2 z-score threshold, and MHC inclusion (**Fig S12**). Second, we compared L2G-prioritized GWAS genes for the same diseases (**Methods,** Mountjoy et al. 2021) and found specific hubs strongly enriched for immune diseases (OR=3.50). Specific and shared hubs modestly enriched for non-immune GWAS genes (OR=2.08, 1.43, **Figs 5c**, **S13**), similar to bulk network hubs (Mostafavi et al. 2023); these results are likely due to confounding by baseline annotations (Kim et al. 2019). Third, using Genomics England PanelApp (Martin et al. 2019) we found that specific hubs were enriched in immune and depleted in non-immune Mendelian genes, while shared hubs had no enrichments (**Fig 5d**).

We compared hubs to drug targets. First, using OpenTargets, we found that specific hubs enriched for approved immune drug targets, but not non-immune drug targets, while shared hubs showed no enrichment (**Fig 5e**). Second, using target-indication pairs from Citeline Pharmaprojects (Minikel et al. 2024), we found that specific hubs had 1.54-fold relative success for phase 1 immune drug targets, similar to the 1.68-fold relative success using indication-matched genetic support. Importantly, combining genetic evidence with specific hubs substantially increased relative success (2.67-fold, Fisher’s p=0.008). These enrichments were null for shared hubs (**Fig 5f**), consistent with prior findings that bulk RNA-seq coexpression networks (Sadler et al. 2023) and physically interacting genes from IntAct (Tan et al. 2025) add minimal utility for drug targets.

Generally, bulk glasso hubs had similar but weaker enrichments as cell type-specific hubs, while hubs from pseudobulk glasso and bulk and pseudobulk WGCNA gave weak or no enrichment (**Figs S14-15**). Together, genetic variation in specific hubs, but not shared hubs, primarily shapes gene regulatory effects on immune disease.

### Specific networks capture *trans* regulation underlying immune disease

A major goal in human genetics is understanding how effects propagate through *trans* networks to ultimately impact complex traits. We evaluated genes with known *trans* molQTL and found that specific hubs, but not shared hubs, were broadly enriched (**Figs 6a**, **S16**). We found the same pattern for *trans* heritability of gene expression (Price et al. 2011; Sun et al. 2023; Wheeler et al. 2016; Wright et al. 2014; **Ext Data Fig 3 c,d**). WGCNA and glasso hubs showed weaker or null enrichments in *trans*-molGenes (**Fig S17**).

**Figure 6.**
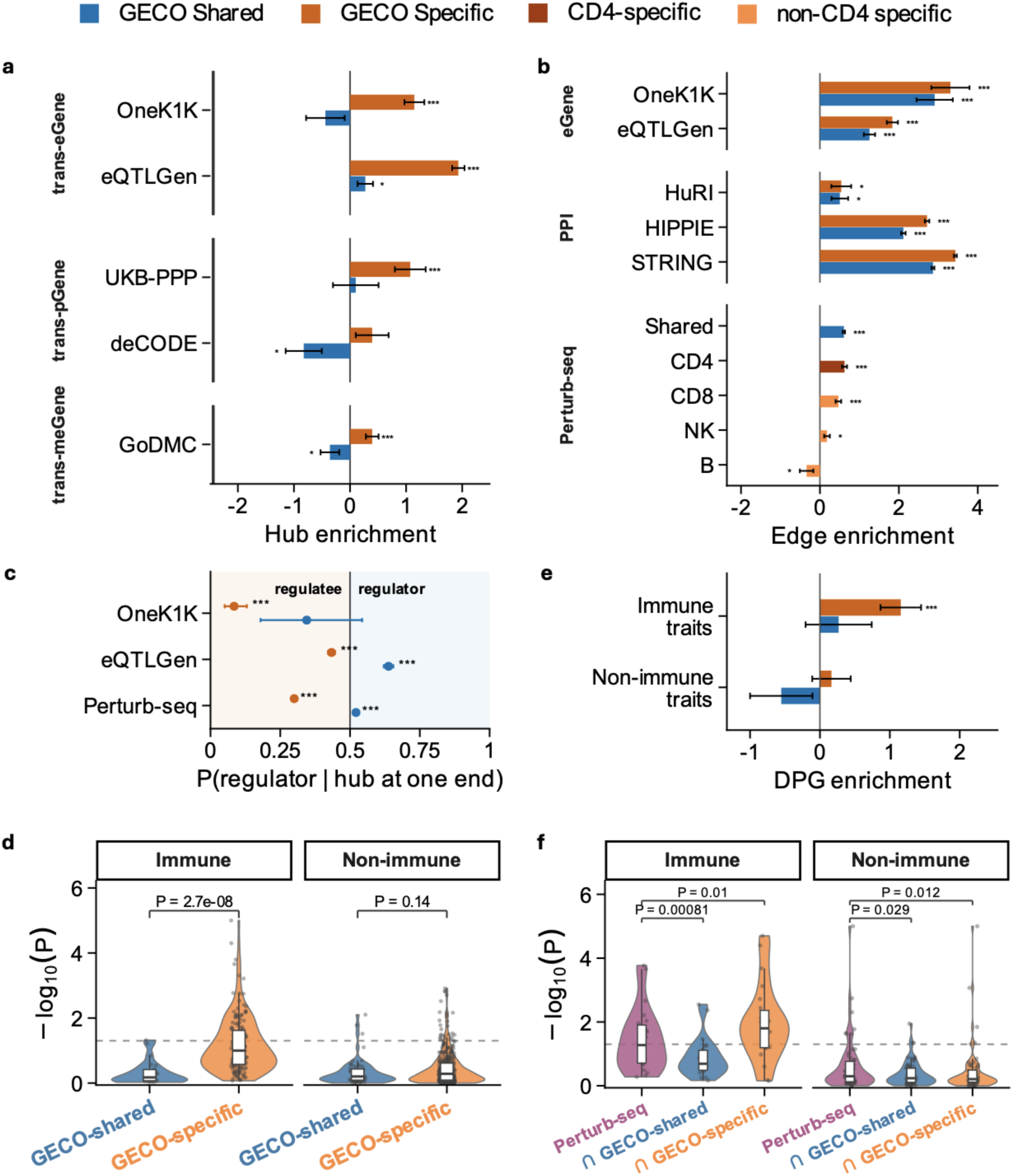
Cell type-specific networks capture *trans*-regulation effects on complex traits. **(a)** Cell type-specific hubs were enriched for *trans*-molecular QTL genes: *trans*-eGenes from OneK1K and bulk blood tissue (eQTLGen); *trans*-pGenes from plasma (UKB-PPP); and *trans*-meGenes from blood (GoDMC). deCODE has low power since 87.8% of considered genes are *trans*-pGenes. Shared hubs were enriched only in eQTLGen, the largest dataset. (**b**) GECO network edges were enriched for: colocalized *cis*-*trans* eGene pairs from OneK1K and eQTLGen; PPIs from binary interaction screens (HuRI), affinity-purification mass spectrometry (BioPlex HCT116 and 293T, **Fig S18**), and integrated databases (HIPPIE and STRING); and perturbation-response pairs from perturb-seq in primary human CD4 T cells. **(c)** Specific hubs were enriched in regulatees (*trans* genes in *cis-trans* eGene pairs or the response genes in perturb-response pairs), while shared hubs were enriched in regulators. Bars show 95% CI from a two-sided binomial test. (**d**) NetworkMAGMA results for shared and specific edges **(d)** and perturb-seq regulator-regulatee pairs before and after intersecting with GECO edges **(f)**. P-values show two-sided Wilcoxon tests, with a dashed line at p = 0.05. **(e)** Specific hubs enriched for disease-proximal genes in immune traits identified by DANDELION. In **a**, **b**, and **e**, x-axes represent log2 odds ratios and error bars indicate +/- 1 standard error.

While GECO only models steady-state expression, we asked if its network edges capture causal *trans* effects. First, using colocalized eQTLs from eQTLGen Phase 2 (Warmerdam et al. 2026) and OneK1K (Yazar et al. 2022) that affect both a *cis* regulator gene and a *trans* regulatee gene, we found that *cis*-*trans* eGene pairs were enriched in shared and specific edges (**Figs 6b, S18**). Second, shared and specific edges also enriched for protein-protein interactions across four databases (HuRI, BioPlex, HIPPIE, and STRING, **Figs 6b, S18**). Third, we used perturb-seq from primary CD4 T cells to create pairs of *cis* regulators (perturbed gene) and *trans* regulatees (differentially expressed after perturbation). Shared and CD4-specific GECO edges enriched for these *cis-trans* pairs from CD4 perturb-seq, with weaker enrichment in CD8 and NK edges and depletion in B edges (**Figs 6b, S18**). Both colocalized eGene pairs and perturb-seq pairs showed cell type-specific hubs skewed toward *trans* regulatees while shared hubs skewed toward *cis* regulators (**Fig 6c**); this was robust to defining regulator and regulatee by effect size (**Fig S19**). WGCNA and glasso hubs had no consistent directionality, consistent with mixing upstream shared hubs and downstream specific hubs (**Fig S20**).

We asked if GECO edges capture *trans* regulatory effects on disease using a novel edge-level enrichment test that combines MAGMA gene-level GWAS scores (Leeuw et al. 2015) and the G statistic (Getis and Ord 1992, **Methods**). We validated our test, NetworkMAGMA, in simulations (**Ext Data Fig 4 a, Fig S21**). We found that NetworkMAGMA gave consistent but stronger results than LDSC hub enrichment, as expected since it uses the entire network (**Ext Data Fig 4 b, Fig S23**). Cell type-specific edges showed stronger enrichment than shared edges in immune disease, but not non-immune disease (**Fig 6d**). NetworkMAGMA linked cell types to biologically plausible diseases, such as CD4ET to psoriasis (p=2.2e-04), CD4NC to autoimmune hypothyroidism (p=1.0e-05), and CD4 and B cell to rheumatoid arthritis (p=8.1e-03 to 4.9e-04, **Fig S22**). Standard glasso networks from bulk and pseudobulk had weaker NetworkMAGMA enrichments in immune disease than cell type-specific networks (p=1.7e-3 and 1.4e-6, respectively, **Ext Data Fig 6**). We found similar results for disease-proximal genes identified by DANDELION (Salamone et al. 2026), which leverages *trans* eQTL effects on complex traits (**Figs 6e, S24**): enrichment for specific hubs in immune traits, but not for non-immune traits or shared hubs.

Perturb-seq captures a mix of disease-related and -unrelated gene regulation (Ota et al. 2026). We hypothesized that shared vs specific networks would partly distinguish these signals. Indeed, we found that intersecting perturb-seq gene pairs with cell type-specific edges increased immune and decreased non-immune disease enrichment in NetworkMAGMA; further, intersecting with shared edges decreased both enrichments (**Fig 6f**). These gains were lineage-specific: immune GWAS enrichment increased most when intersecting with CD4 GECO edges, yet decreased when using B cell edges (**Ext Data Fig 5 a**). The intersected networks illustrate how shared GECO edges identify trait-irrelevant housekeeping signals in perturb-seq, while GECO CD4 edges identify disease-related CD4 signals (**Ext Data Fig 5 b, Figs S25-26**).

Overall, GECO networks capture causal *trans* regulation, with shared hubs skewed toward upstream regulators and specific hubs skewed toward downstream, disease-proximal regulatees.

### Replication in CLUES/ImmVar reveals BAFF biology in lupus

We replicated our main findings in CLUES/ImmVar, an independent cohort measuring PBMCs in N=147 individuals with systemic lupus erythematosus (SLE) and N=96 healthy controls. We again found that shared hubs were more evolutionarily constrained and enriched in housekeeping processes, while specific hubs were rapidly evolving and enriched in dynamic regulation and immune GO terms (**Figs S27-28**). Specific hubs were enriched for immune disease heritability (**Fig 7a**), GWAS genes (**Fig 7b**), Mendelian genes (**Fig 7c**), drug target genes (**Fig 7d**), and drug target success rate (**Fig S32)**; further, specific network edges showed NetworkMAGMA enrichment in immune disease GWAS (**Figs 7e**, **S34**). These enrichments were weaker or absent in non-immune diseases and shared networks (**Fig 7**). These results replicate the distinct biology of cell type-shared and specific networks.

**Figure 7.**
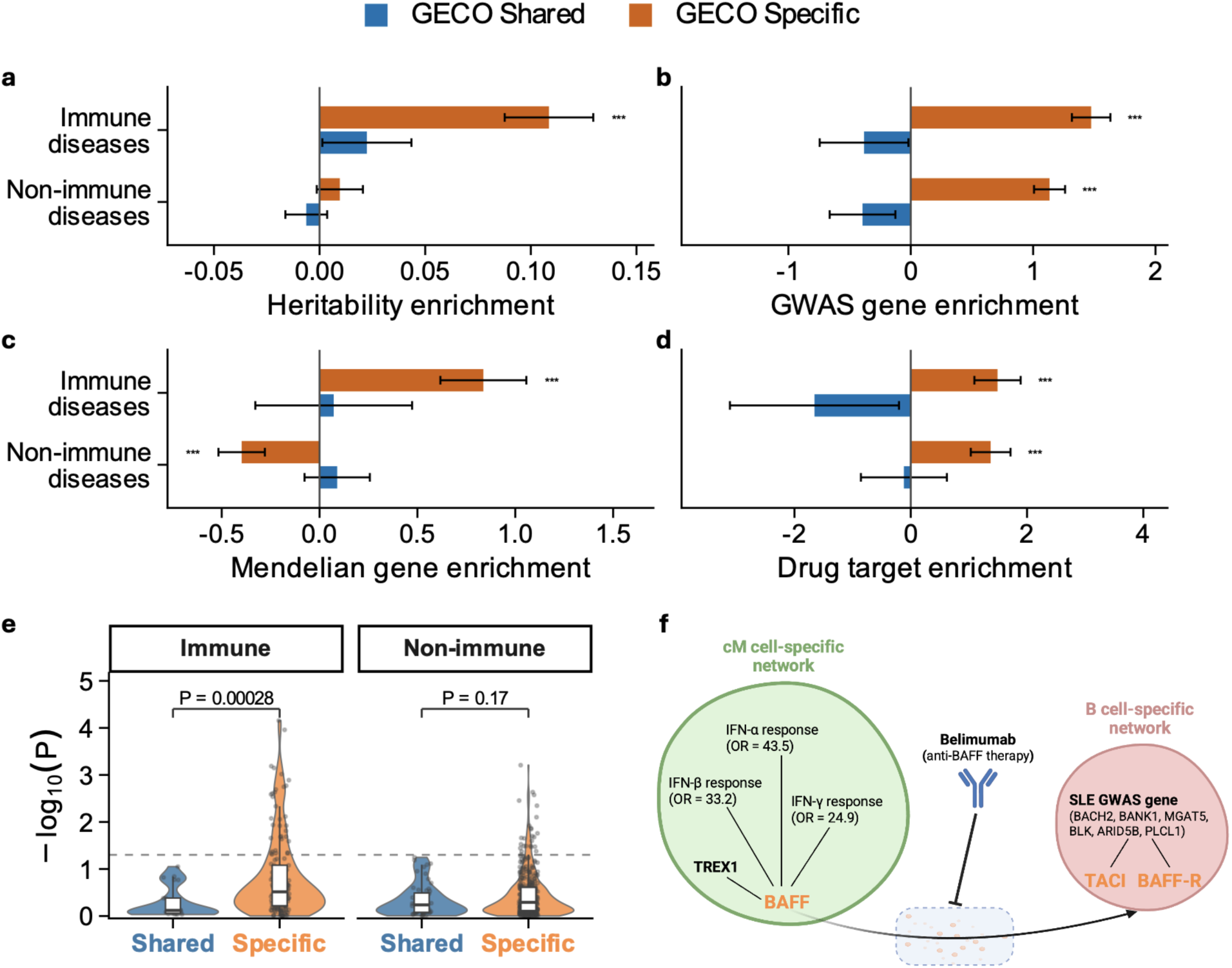
Shared and specific network properties replicate in CLUES/ImmVar and connect lupus genetic risk to a drug target mechanism. GECO hub gene enrichment in GWAS heritability using LDSC **(a)**, GWAS genes using L2G **(b)**, Mendelian genes **(c)**, and drug target genes **(d)**. One outlier, metformin, is excluded from **d** (**Fig S31**). **(e)** GECO gene-gene edge enrichment from NetworkMAGMA; P-values use two-sided Wilcoxon tests. The dashed line denotes p = 0.05. **(f)** Belimumab is an antibody directed against BAFF, which is secreted by monocytes and binds to TACI and BAFF-R on B cells (Scott et al. 2012). BAFF, TACI, and BAFF-R are cell type-specific hubs in the appropriate cell types; edges show network connections to a Mendelian gene (*TREX1*), six GWAS genes, and three interferon pathways enriched among connected genes (**Ext Data** Fig 7). In **b**, **c**, and **d**, x-axes show log2 odds ratios. Error bars indicate +/- 1 standard error.

We used GECO networks to explore B cell activating factor (BAFF) signaling, an interferon-related driver of B cell expansion targeted by belimumab, the first therapy specifically developed and approved for SLE (Navarra et al. 2011). We found *BAFF* is a monocyte (cM) hub (**Fig 7f**), consistent with cM being the dominant PBMC source of *BAFF* (Moore et al. 1999); it was connected to *TREX1*, a Mendelian SLE gene (Lee-Kirsch et al. 2007), and genes enriched in interferon response (p<3e-10, **Fig 7f, Ext Data Fig 7**). We also found that *BAFF* receptors (BAFF-R and TACI) were B cell hubs connected to six SLE-associated GWAS genes (**Fig 7f**). Interestingly, *BAFF* harbors a SLE-associated variant with signatures of positive selection (Steri et al. 2017), consistent with cell type-specific hubs enriching for rapidly evolving genes. Together, these results show how cell type-specific networks can inform mechanisms underlying genetic associations and connect them to drug targets.

## Discussion

We found that cell type-shared and cell type-specific interindividual coexpression networks capture qualitatively distinct biology. Shared networks capture housekeeping processes and dominate standard bulk and pseudobulk approaches, whereas specific networks capture dynamically regulated core disease genes.

Core genes should be highly connected to peripheral genes (Liu et al. 2019; Salamone et al. 2026), which is often true for Mendelian genes (Gordon et al. 2026; Vuckovic et al. 2020), yet bulk coexpression hubs are depleted in known eQTLs (Mostafavi et al. 2023) and not enriched in GWAS heritability above baseline annotations (Kim et al. 2019). Our results explain this apparent paradox: networks do funnel genome-wide genetic risk into core genes, but this occurs in cell type-specific networks that are obscured by housekeeping processes in standard methods. This extends our prior finding that shared eQTLs obscure the cell type-specific eQTLs that explain missing *cis* gene regulation in immune disease (Minhui Chen et al. 2026).

Cell type-specific hubs enriched for drug target success, comparable and complementary to genetic evidence. These enrichments were absent for shared hubs. In the lupus cohort, specific networks informed how genetic and therapeutic effects on lupus funnel into *BAFF* signaling from monocytes to B cells. More generally, cell type-specific networks connected Mendelian genes to drug targets (**Fig S33**), primarily in likely-causal cell types (**Table S2**). This suggests that genes connected to drug targets in appropriate cell types could predict drug response.

Shared and specific networks have distinct evolutionary histories. Shared hubs enriched for constrained housekeeping genes, consistent with protein network hubs in yeast (Fraser et al. 2002; Jeong et al. 2001). Specific hubs were less constrained than shared hubs, consistent with models in which pleiotropic effects experience stronger constraint (Simons et al. 2018; Spence et al. 2026; Koch et al. 2024). But our results go a step further: specific hubs are depleted in constraint and may even experience directional selection, which is inconsistent with stabilizing selection but consistent with positive selection on immune genes (Barreiro and Quintana-Murci 2010).

Our results extend bulk tissue models of shared and specific networks (Pierson et al. 2015; Gao et al. 2016; Xie et al. 2016; Saha et al. 2017). Consistent with our results, tissue-shared hubs are highly constrained and depleted of Mendelian and drug target genes (Ravichandran et al. 2025), and tissue-specific hubs enrich for tissue-specific TFs (Pierson et al. 2015) and pathways (Saha et al. 2017). However, tissue-specific hubs have not been linked to dynamic regulation or core disease genes, suggesting that cell type-level specificity is important for identifying disease-related *trans* regulation. In turn, this suggests that specific networks in finer cell subtypes or states will be even more disease-related (Nathan et al. 2022), at the cost of higher noise.

GECO coexpression edges were enriched in *trans* effects identified by perturb-seq in CD4 cells, with greatest enrichment in shared and CD4 edges and depletion in B cell edges. This shows both that GECO edges partly capture causal *trans* regulation and that perturb-seq identifies a mix of shared and specific signals. The latter partly explains why perturb-seq signals have mixed relevance to complex traits (Ota et al. 2026) and suggests paths to improve the utility of perturb-seq; for example, disease relevance increased after intersecting perturb-seq with cell type-matched networks, but decreased when using shared or cell type-mismatched networks.

Several challenges remain. First, our results are limited to PBMCs; while our approach is general, rapidly evolving cell type-specific networks may be most relevant in brain and immune traits (Christmas et al. 2023; Mangan et al. 2022). Second, GECO networks are trained on steady-state data and need not learn causal regulatory effects (Kernfeld et al. 2024), unlike gene regulatory network models using TF binding motifs and scATAC-seq (Bravo González-Blas et al. 2023; Kamimoto et al. 2023) or perturb-seq (Brown et al. 2025; Ota et al. 2026); nonetheless, GECO edges are enriched in causal *trans* regulation effects from perturb-seq and *cis*-*trans* eQTL pairs. Third, GECO accounts for cell-to-cell variation within cell types but does not model networks within cell types. Fourth, pre-processing pseudobulk data could improve GECO network inference (Lu and Keleş 2023). Finally, coexpression networks output edges and hubs rather than latent factors as in non-negative matrix factorization (Ota et al. 2026) or tensor decomposition (Hore et al. 2016; Mitchel, Gordon, et al. 2025), but our results suggest that stratifying factors into shared and specific axes should be beneficial.

## Methods

### GECO overview

GECO estimates cell type-specific interindividual gene coexpression networks from population-scale scRNA-seq data and pre-defined cell types. GECO has two steps: first, it unbiasedly estimates cell type-shared and -specific interindividual covariance between all gene pairs using method of moments; second, it fits network edges and hub genes using a novel weighted graphical lasso (Friedman et al. 2008).

In the first step, GECO fits a linear mixed model to pseudobulk data:

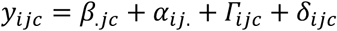

where *y_i_*_j*c*_ is the pseudobulk expression level of gene j in individual i and cell type c, *β*_.j*c*_ is the mean expression level across individuals of gene j in cell type c, *α_i_*_j._ is the individual effect that is shared across cell types, *Γ_i_*_j*c*_ is the individual effect that is specific to cell type c, and *δ_i_*_j*c*_ is the cell-to-cell variation for gene j in individual i and cell type c. We model *β*_.j*c*_ as fixed effects, *α_i_*_j._as a random effect with *α_i_*∼*^iid^N*(0, *V*), and *Γ_i_*_j*c*_ as a random effect with *Γ_i_*_.*c*_∼*^iid^N*(0, *Ω^c^*). By the CLT, we assume 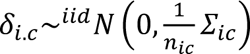, where Z*_ic_* is the cell-to-cell gene covariance matrix in individual i and cell type c which is estimated from cell-level data prior to constructing pseudobulk (Minhui Chen and Dahl 2024, **Note S1**) and *n_ic_* is the number of cells for individual i in cell type c. Thus, GECO partitions interindividual covariance across genes into three components: cell type-shared (*V*), cell type *c*-specific (*Ω^c^*), and cell-to-cell variation 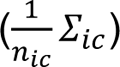

GECO fits this model unbiasedly using the method of moments with population-scale data (**Note S1**). Using multilinear algebra (**Note S1**), we derived simple expressions for these estimators:

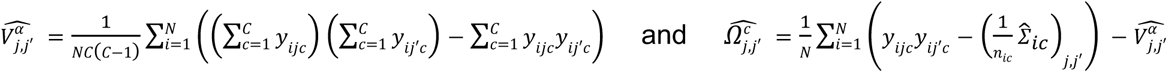

where *V^α^_j,j_*, and *Ω^c^_j,j_* are the cell type-shared and specific covariance of gene *j* and gene *j*′, N and C are the number of individuals and cell types. There are three benefits to this simplification. First, it provides an interpretable description of GECO estimates as decomposing variance. Second, it enables closed-form standard errors, eliminating computationally-intensive jackknifing (**Note S1**). Third, this gives a more efficient estimator for our prior CTMM model as a special case (when *j*=*j*’, Minhui Chen and Dahl 2024). Overall, this simplification enables GECO to scale to genome-wide application to population-scale single-cell datasets.

### Weighted graphical lasso

We developed a novel approach to glasso to fit transcriptome-wide networks from GECO covariance estimates. This step converts marginal covariance patterns into partial covariance relationships, retaining gene-gene connections that remain after conditioning on all other genes. A novel approach is necessary due to the noise in single-cell data, yielding transcriptome-wide covariance matrix estimates that have many negative eigenvalues and heterogeneous noise levels across entries. We handle the former with standard truncated SVD, i.e., zeroing out negative eigenvalues. We handle the edge-level noise using GECO’s uncertainty estimates to weight each gene-gene edge. Specifically, we calculated BH FDR, converted into z-scores, and then penalized each edge proportional to 1/(|z|+0.1). This preferentially penalizes noisy edges. We selected the scalar tuning parameter in GLasso using eBIC (Foygel and Drton 2010).

### Network MAGMA

We developed a novel approach to network-level testing of gene-level effects on complex traits. We used General G statistic (Getis and Ord 1992) to evaluate the relevance of each network to gene-level effects on complex traits based on MAGMA gene-level GWAS scores (Leeuw et al. 2015). This topology-aware analysis goes beyond asking whether trait-associated genes are present in networks hubs and instead tests enrichment in the networks edges themselves.

Specifically, we used

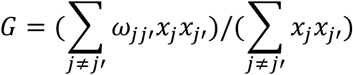

where *x*_j_ is the gene-level complex trait-relevance score and *ω*_jj′_ is the partial correlation of gene *j* and gene *j*′. Excluding *j* = *j*′ from the sum ensures our statistic measures network topology itself rather than solely overlap between GWAS signals and hub genes. In this paper we used the MAGMA -*log*_10_(*p*_j_) for *x*_j_ and the absolute value of the GECO network partial correlations for *ω*_jj′_ (absolute value because the MAGMA -log10(p-value) does not depend on the sign of the effect), but other gene-level scores could be used such as rare variant burden scores. To assess significance, we fixed each network (*ω*_jj′_) and randomly permuted *p*_j_ across j, which tests if trait-relevant gene pairs overlap GECO network edges more than expected if the same MAGMA signals were randomly assigned to genes. Thus, significant results indicate that trait relevance is aligned with GECO network connectivity, rather than reflecting the network density or the marginal distribution of MAGMA scores (**Ext Data Fig 4 a, Fig S21**). We conservatively removed gene pairs within the MHC due to long-range LD. We used a binary variant (*ω*_jj′_ = 0 or 1) when comparing GECO networks to edges from perturb-seq and standard glasso networks.

### Simulations

#### Covariance decomposition

Simulation parameters were derived from the OneK1K data. We simulated gene pairs with prespecified shared, cell type-specific, and measurement-noise covariance components. For each component, variances were set to either the median or the 75th percentile of the corresponding empirical estimates. Correlations were varied from −1 to 1 in increments of 0.1 and converted to covariances using the specified variances. Cell type-specific mean expression levels were similarly set to the median or the 75th percentile. Bulk expression was generated as a cell type-proportion-weighted sum of simulated pseudobulk expression. Each setting was repeated 1,000 times.

#### Hub and edge recovery

We performed genome-wide simulations using all 11,526 OneK1K genes. Shared and cell type-specific covariance matrices were derived from GECO-estimated precision matrices, and pseudobulk expression was simulated using empirical mean expression levels and individual- and cell type-specific measurement noise covariance matrices. GECO, WGCNA, and standard glasso were then applied to each simulated dataset to evaluate hub and edge recovery. Each setting was repeated 100 times.

#### Network MAGMA

On each of the GECO networks derived from OneK1K, we ran two parallel simulation settings (binary (0/1 adjacency) or continuous (|partial correlation|)) that differ only in how edge weights are represented. In every setting a causal set of k = G/10 genes was assembled by first drawing m = 5 seed genes uniformly at random and then adding the remaining genes one at a time with a selection probability that tunes how tightly the causal set clusters. In the binary setting the probability was proportional to q for candidates already adjacent to the selected set and 1 otherwise; in the continuous setting it was proportional to1 + (*q* − 1)(*a*_j_/*w*L), where *a*_j_ = ∑*_s_*_∈_*_S_ w*_j*s*_ is the cumulative partial-correlation weight of candidate j to the already-selected set S and *w*L is the mean edge weight. Each gene received a continuous score *x*_j_ = *F*^N–1^(*φ*(*z*_j_)) via a Gaussian copula, where the latent *z*_j_∼*N*(0,1) for non-causal genes and *z*_j_∼*N*(1,1) for causal genes, and *F*^N^ is the empirical CDF of pooled MAGMA −*log*_10_(*p*). For each q we simulated 1,000 independent replicates and computed the networkMAGMA p-value under both a global label-permutation null and a degree-stratified permutation null, the latter evaluated across a range of degree-bin resolutions (10, 20, 50, 100, and 200 quantile bins).

#### OneK1K data

Our data analyses closely followed our prior study (Minhui Chen and Dahl 2024). In brief, we applied GECO to 928 individuals and 11,526 genes (66,418,575 gene pairs) in the 7 most common cell types (CD4+ Naive and Central Memory T cell (CD4NC), CD4+ Effector Memory T cell (CD4ET), CD8+ Naive and Central Memory T cell (CD8NC), CD8+ Effector Memory T cell (CD8ET), Natural Killer cell (NK), Immature and Naive B cell (BIN), Memory B cell (BMem)). We adjusted for the same fixed effects as in our prior work (Minhui Chen et al. 2026) except for genotype PCs.

#### CLUES data

We used the same quality control procedures as that in our prior work (Minhui Chen et al. 2026), except that we removed the filters required for eQTL mapping (here, we included individuals without genotype data and controls of Asian ancestry). We applied GECO to 243 individuals (147 SLE cases and 96 healthy controls) and 11,087 genes (61,455,241 gene pairs) in the seven most common cell types (CD4 T cells (CD4), CD8 T cells (CD8), natural killer cells (NK), B cells (B), CD14 classical monocytes (cM), CD16 nonclassical monocytes (ncM), conventional dendritic cells (cDC)). We adjusted for the same fixed effects as in our prior work (Minhui Chen et al. 2026) except for genotype PCs.

#### WGCNA and standard graphical lasso

We applied WGCNA (Langfelder and Horvath 2008) and standard glasso (Friedman et al. 2008) to the pseudobulk and overall bulk expression data, adjusting for the same covariates used in GECO. For WGCNA, we excluded genes in the grey module, which were not assigned to any detected coexpression module (Langfelder and Horvath 2008), and selected the top genes by kME from each remaining module in proportion to module size; the union of these genes was defined as the hub gene set. For standard glasso, we defined hubs by degree, as in GECO. In simulations, we selected the same number of hub genes as in the ground truth; in real data, we defined the top 5% of connected genes as hubs.

#### Gene set enrichment analyses

We tested enrichment of each gene set in GECO Shared and GECO Specific hubs using logistic or linear regression. In our baseline test, no covariates are used. In secondary analyses, we adjust for the covariates of overall expression (mean overall expression across individuals), expression variability (variance of mean expression across cell types), gene length, and GC content, subsetting to genes where all these covariates are measured. Severely skewed variables were log-transformed when all values were positive and log1p-transformed when zero values were present (**Table S3**). For gene sets defined only within a subset of genes (e.g., protein coding genes), we restricted enrichment analyses to GECO genes in this subset. TSS coordinates and gene lengths were obtained from the Ensembl GRCh38 gene annotation (Rainer et al. 2019). Gene-body GC content was calculated as the fraction of G and C bases across each annotated gene span.

#### Candidate gene sets

**Housekeeping genes** were taken from the HRT Atlas v1.0 HumanHK list (Hounkpe et al. 2021), which are defined as genes with stable cross-tissue expression. **Immune-cell marker genes** were taken from the LM22 signature matrix (Newman et al. 2015), defined as the CIBERSORT 22-cell-type reference for bulk-PBMC deconvolution. **Gene burstiness** was quantified for CD34+ cells using the homeostasis Z-index (Mengjie Chen 2025), which is high for genes that are highly expressed in a small set of cells; we binarized bursty genes by taking those with FDR < 0.05. **Low/High-dispersed genes** and **cell type-differentially dispersed genes** were taken from the cardiac culture single-cell data from Jamison et al. (Jamison et al. 2026); we binarized by taking genes in the bottom/top 20% of cell-to-cell dispersion in all 6 cell types, separately, and took cell type-differentially dispersed genes as those with FDR < 0.05 in at least one of the 15 cell type pairs. Gene identifiers were mapped to Ensembl IDs from genenames.org (Seal et al. 2026) and were excluded if they did not map one-to-one. **GO term enrichments** were tested using “WebGestalt” (Elizarraras et al. 2024).

#### Evolutionary measures

##### LOEUF

Gene-level loss-of-function intolerance was quantified using LOEUF from gnomAD v4.1 (S. Chen et al. 2024), where lower values indicate stronger depletion in loss-of-function variants in human populations. We retained transcripts annotated as canonical or MANE Select transcripts. For genes with multiple transcript records, we took the minimum LOEUF. Genes with missing or non-finite LOEUF values were excluded.

##### phyloP

We used the mean score across protein-coding sequences for each gene, which was derived from comparative genomics across hundreds of placental mammals (Christmas et al. 2023). Higher phyloP indicates stronger cross-species conservation. For genes with multiple protein-coding records, we took the maximum value across records.

##### Background selection

Background selection was quantified using the McVicker et al. B statistic (McVicker et al. 2009), where lower B indicates stronger linked purifying selection. The hg18 B map was decoded from its run-length-encoded format into genomic segments. Ensembl gene bodies and TSS from GRCh38/hg38 were lifted to hg18 using the UCSC hg38ToHg18 liftOver chain file. Each gene was assigned the base-pair-weighted mean B value across its lifted gene body, excluding regions that failed to map across assemblies.

##### zooHAR

We used the 312 high-confidence Zoonomia human accelerated regions (zooHARs) reported by Keough et al. as evolutionary regulatory annotations (Keough et al. 2023). We further defined a positive-selection subset by retaining zooHARs annotated as positive, yielding 257 regions.

##### HAQER

We used 1,581 human ancestor quickly evolved regions (HAQERs) from Mangan et al. (Mangan et al. 2022). HAQERs are human genomic regions highly divergent from the human-chimpanzee ancestor and are characterized by elevated mutation rates, positive selection, and enrichment for disease-linked variation.

##### Akbari et al 2026

We obtained 479 independent loci under directional selection identified from ancient DNA time-series data (Akbari et al. 2026). For loci with reported gene annotations in the original study, we used the annotated genes directly. The remaining loci were mapped to genes using the below approaches. Genomic coordinates were converted to GRCh38/hg38 using the UCSC hg19ToHg38 liftOver chain file, retaining only loci with unique mappings.

##### Region-to-gene

We used ENCODE-rE2G thresholded predictions (Gschwind et al. 2026) from the released blood primary immune cell biosamples matched to the cell types used in this paper. For each cell type, the annotation was defined as the union of all genes linked by rE2G in that cell type. For shared hubs, enrichment was tested on the union of genes mapped across the seven cell types. For cell type-specific hubs, enrichment was assessed separately for each of the seven cell type-specific hub gene lists and combined across cell types by inverse-variance meta-analysis.

### Transcription factors

Transcription factor sets were obtained from the Human Protein Atlas (Uhlen et al. 2019), comprising “low-specificity” TFs and “lineage-elevated” TFs for T cells, NK cells, and B cells. To test whether GECO can predict TF lineage, every annotated TF in the GECO universe (n = 726) was assigned to one or more “predicted” sets based on the network in which it attained its maximum degree. TFs reaching their maximum in any of the four T-cell networks, in the NK network, or in either B-cell network were assigned to the “predicted T cell”, “predicted NK cell”, and “predicted B cell” sets, respectively.

### Protein-protein interactions (PPI)

We used four protein-protein interaction resources: HuRI, a systematic proteome-wide human reference interactome generated using binary interaction screens (Luck et al. 2020); BioPlex 3.0, generated in two cell lines (HCT116 and 293T) using affinity-purification mass spectrometry (Huttlin et al. 2021); HIPPIE v2.4, which integrates multiple types of experimental evidence (Alanis-Lobato et al. 2017, v. 20); and STRING v12.0, including both the full network and the physical interaction network, retaining interactions with a confidence score ≥700 (high confidence according to STRING) (Szklarczyk et al. 2023).

### Gene regulatory features

#### Promoter activity

We obtained per-sample TPM (tags per million) values and peak coordinates from the FANTOM5 hg38 v9 reprocessed CAGE atlas (Forrest et al. 2014). CAGE peaks were assigned to genes using FANTOM5 annotations. To match the cell types in OneK1K, standard donor primary-cell samples were grouped into four merged lineages: CD4 T cells, CD8 T cells, NK cells, and B cells. A peak was considered active in a cell type if its TPM was >0 in any matched donor, and the per-gene promoter score was defined as the number of distinct active peaks annotated to that gene.

#### Enhancer activity

Thresholded enhancer-gene predictions were obtained from ENCODE-rE2G (Gschwind et al. 2026) for biosamples matching each OneK1K cell type. Within each focal cell type, each gene was assigned the number of unique enhancer regions linked to it as the union across donors.

#### TF binding

We obtained TRACE-derived TF footprint annotations from ENCODE for blood biosamples matched to seven OneK1K immune cell-type labels (Ouyang and Boyle 2020). For annotations listing multiple co-targeted TFs, we credited each footprint to every listed TF for the diversity metric, whereas density counted each footprint once. For each gene, we defined a standardized ±100 kb window around the strand-specific TSS and considered footprints whose midpoint fell within this window. We computed two per-gene metrics within each cell type: TF diversity, defined as the number of distinct TFs with at least one footprint in the window, and TF density, defined as the total number of footprints in the window.

#### molQTL data

##### Significant molGenes

OneK1K *cis*-/*trans*-eGenes and *cis*-*trans* eGene pairs were obtained from the OneK1K project (Yazar et al. 2022). eQTLGen *cis*-/*trans*-eGenes were derived from bulk whole-blood eQTL mapping from eQTLGen Phase 1 (Võsa et al. 2021). The *cis*-*trans* eGene pairs were obtained from the colocalized *cis*-eGene–*trans*-eGene pairs identified in Phase 2 (Warmerdam et al. 2026). DICE eGenes were obtained from the DICE project (Schmiedel et al. 2018), comprising *cis*-eGenes across 15 sorted immune cell types and activation states. ImmuNexUT eGenes were obtained from the ImmuNexUT study (Ota et al. 2021), which profiled bulk RNA-seq of 28 FACS-sorted immune cell populations from East Asian donors. Memory T cell state-dependent eGenes were obtained from Nathan et al. 2022 (Nathan et al. 2022). We selected cell-state-dependent eGenes with significant genotype × cell-state (top 7 independent canonical correlation analysis dimensions) interaction effects. CLUES shared/specific *cis*-eGenes were obtained from Perez et al. (Perez et al. 2022). GTEx shared/specific eGenes were from fastGxC (Krockenberger et al. 2026). UKB *cis*-/*trans*-pGenes were derived from the discovery cohort of the UK Biobank Pharma Proteomics Project (UKB-PPP) (Sun et al. 2023). deCODE *cis*-/*trans*-pGenes were derived from the deCODE plasma proteomics resource (Ferkingstad et al. 2021). Blood *cis*-/*trans*-meGenes were derived from GoDMC (Min et al. 2021). We mapped CpG sites to genes using the Illumina HumanMethylation450 manifest. PBMC single-cell *cis*-caGenes were derived from the meta-analyzed caQTL summary statistics of peripheral immune cells (Mu et al. 2026). We mapped peaks to genes with the nearest TSS. Intestine single-cell *cis*-caGenes were derived from the project conducted in CD cases and controls (Zhao et al. 2024). We mapped peaks to genes by co-activity results provided by the original paper.

##### Heritability estimates

We assembled per-gene expression heritability estimates from eight published datasets of whole blood or peripheral immune cells with genome-wide estimates: GCTA-GREML (Yang et al. 2011) unconstrained *cis* h2 for GTEx v8 whole blood (The GTEx Consortium 2020); GCTA-GREML constrained *cis* h2 and unconstrained *trans* h2 for DGN (Wheeler et al. 2016); GCTA-GREML constrained *cis* h2 for the NTR RNA-seq data (Ouwens et al. 2020); GCTA-GREML constrained *cis* h2 and unconstrained twin-based additive h2 for an earlier array-based NTR dataset (Wright et al. 2014; we defined *trans* h2 as the difference between twin and *cis* h2); unconstrained IBD-based *cis* and *trans* h2 from Icelandic families (Price et al. 2011); CIGMA unconstrained shared and cell type-specific *cis* h2 in OneK1K and CLUES/ImmVar (Minhui Chen et al. 2026); and, at the protein level, SuSiE-fine-mapped *cis*/*trans* pQTL variance explained and genome-wide LDSC h2 (excluding pQTL) for UKB-PPP (Sun et al. 2023). For each hub set, we fit median regression of per-gene heritability on hub membership and bootstrapped genes 10,000 times to calculate confidence intervals. We used medians rather than means because they are more robust to outliers (Minhui Chen et al. 2026) and because several studies have biased mean estimates due to constraining heritability estimates to be nonnegative (Steinsaltz et al. 2020).

### Complex trait data

#### Disease panel

We chose a consistent set of disease phenotypes spanning FinnGen and OpenTargets; we used the former to estimate heritability and the latter to identify GWAS genes and drug target genes.

We began from the 128 release-3 FinnGen codes in Sakaue et al. GWAS atlas (Sakaue et al. 2021). We then mapped these to release-12 endpoints (Kurki et al. 2023; 89 unchanged, 38 replaced by the closest available equivalent, one dropped, and two phenotype pairs merging into single endpoints), and we added four additional diseases (Crohn’s disease, coeliac disease, multiple sclerosis, and ankylosing spondylitis). This yielded 129 FinnGen R12 GWAS. We then mapped these to 126 EFO disease terms in OpenTargets v26.03 (Buniello et al. 2025). We labelled diseases “immune” if annotated to an immune therapeutic area (immune_system_disease, infectious_disease, or hematologic_disease) and manually labelled three as “immune” (asthma, pediatric asthma, and juvenile rheumatoid arthritis); otherwise, we labelled the disease “non-immune”. This left 117 FinnGen R12 GWAS (29 immune, 88 non-immune) spanning 114 EFO disease terms in OpenTargets (28 immune, 86 non-immune) (**Table S4**).

#### GWAS genes

We used Open Targets Locus-to-Gene (L2G) scores to link GWAS loci to genes (Mountjoy et al. 2021), defining binary gene annotations at an L2G threshold of 0.8. For each disease, we took the union of genes exceeding this threshold across all included GWAS, and then pooled disease gene sets into immune and non-immune groups.

#### Heritability enrichment

We used stratified LD score regression (Finucane et al. 2015) to estimate heritability enrichment in GECO Shared and Specific hub genes. Our primary analyses filtered to the 78 diseases with h2 z-score above 4 (22 immune, 56 non-immune) and condition on the baseline LDSC model (v1.2) in addition to all genes used in GECO. Secondary analyses used the baseline-LD model v2.2, included the MHC, extended *cis* window sizes around hubs from ±100 kb to ±500 kb, changed the h2 z-score filter to 0 or 7, or reduced to the top 210 or 420 hub genes.

#### Drug target genes

We took 459 approved drug targets from OpenTargets v25.03 assigned to at least one immune/non-immune disease, 175 of which are genes in our OneK1K analysis and 184 in our CLUES/ImmVar analysis. We defined the appropriate cell types for drug target genes as either the secreting cell type or the cell type in which the drug acts (for cell-surface and intracellular targets, **Table S2**).

#### Mendelian genes

We obtained the Mendelian genes from Genomics England PanelApp (Martin et al. 2019). Of the 433 panels available, we retained the 296 panels with a version signed off for use in the NHS Genomic Medicine Service. We further excluded 13 Super Panels and three broad phenotype panels (DDG2P, Fetal anomalies, and Intellectual disability). From the remaining 280 disease panels, we retained only green genes (PanelApp confidence level 3; diagnostic-grade evidence). Immune Mendelian panels were defined as the 13 panels in the Immunology specialty plus three immune-mediated panels filed under other specialties on manual review: “IPEX – Immunodysregulation Polyendocrinopathy and Enteropathy, X-Linked”, “Rare genetic inflammatory skin disorders”, and “Intestinal failure or congenital diarrhoea”. All remaining panels were classified as non-immune. This produced 419 immune and 3,556 non-immune Mendelian genes (**Table S5**).

#### Relative success (RS)

We used the approach and data of Minikel et al. (Minikel et al. 2024). RS was defined as the proportion of target-indication pairs that successfully moved between phases, with 95% confidence intervals given by the Katz log method. We restricted the analysis to immune indications, defined as the “immune” therapy area of Minikel et al. (MeSH C20), and genes included in GECO.

#### Disease-proximal genes

Disease-proximal genes (DPG) were the genes defined as the central disease-driving genes and identified by DANDELION for 27 blood traits (Salamone et al. 2026). We grouped these DPGs into immune and non-immune traits (**Table S6**).

#### Perturb-seq

Differential-expression statistics for 33,983 perturbation × 10,282 measured-gene contrasts across Rest, Stim8hr and Stim48hr in primary human CD4+ T cells were obtained from Zhu et al. (Zhu et al. 2026). After retaining on-target knock-downs (KD) without off-target flags and with non-missing, negative self-gene log-fold changes, 18,760 primary contrasts (7,193 unique perturbed genes) remained (6,063 / 6,288 / 6,409 per condition). For every (perturbation P, measured gene m) pair we defined the KD-normalized perturbation effect as |log FC(P,m)| / |on-target log FC(P)|, a dimensionless quantity that removes between-perturbation differences in knock-down efficiency before downstream comparisons.

We selected perturbation-measurement pairs whose both perturbed gene and measured gene were in GECO universe (6,603 unique genes). Then we counted the number of regulators, responders, and neighbors (union of regulators and responders) across all three conditions (adj_p < 0.10) for each gene, and took the mean normalized effect across three conditions as the mean effect size of the perturbed gene on the measured gene in each perturbation-measurement pair. For the network MAGMA analysis, we represented the perturb-seq data as a 6,603 × 6,603 adjacency matrix (edge if either direction significant in any condition), and grouped GECO networks as GECO-Shared adjacency network and GECO-Specific adjacency network (union of 7 GECO cell type-specific networks), as well as GECO CD4-specific network (union of CD4NC- and CD4ET-specific networks), CD8-specific network (union of CD8NC- and CD8ET-specific networks), NK-specific network, and B-specific network (union of BIN- and BMem-specific networks).

## Data availability

OneK1K single-cell gene expression data is publicly available on the Gene Expression Omnibus (GEO) under accession number GSE196830. CLUES single-cell gene expression data is available via GEO at GSE174188.

## Code availability

The GECO R package and all code/data used for analyses are available on GitHub https://github.com/xinpeiw/GECO.

## Acknowledgements

We thank Noah Zaitlen for comments. We thank the Center for Research Informatics (CRI) at the University of Chicago for maintaining the Randi High Performance Computing cluster on which analyses were performed. The Center for Research Informatics is funded by the Biological Sciences Division and the Institute for Translational Medicine/CTSA (NIH UL1TR002389) at the University of Chicago. This work was funded by the National Institute of General Medical Sciences of the National Institutes of Health (R35GM150822 to A.D.).

## Author contributions

X.W. and A.D. developed the statistical methodology, conducted the analyses, and wrote the manuscript. S.S, X.L., and M.C. provided critical suggestions and feedback to the manuscript.

## Extended data figs

**Ext Data Fig 1.**
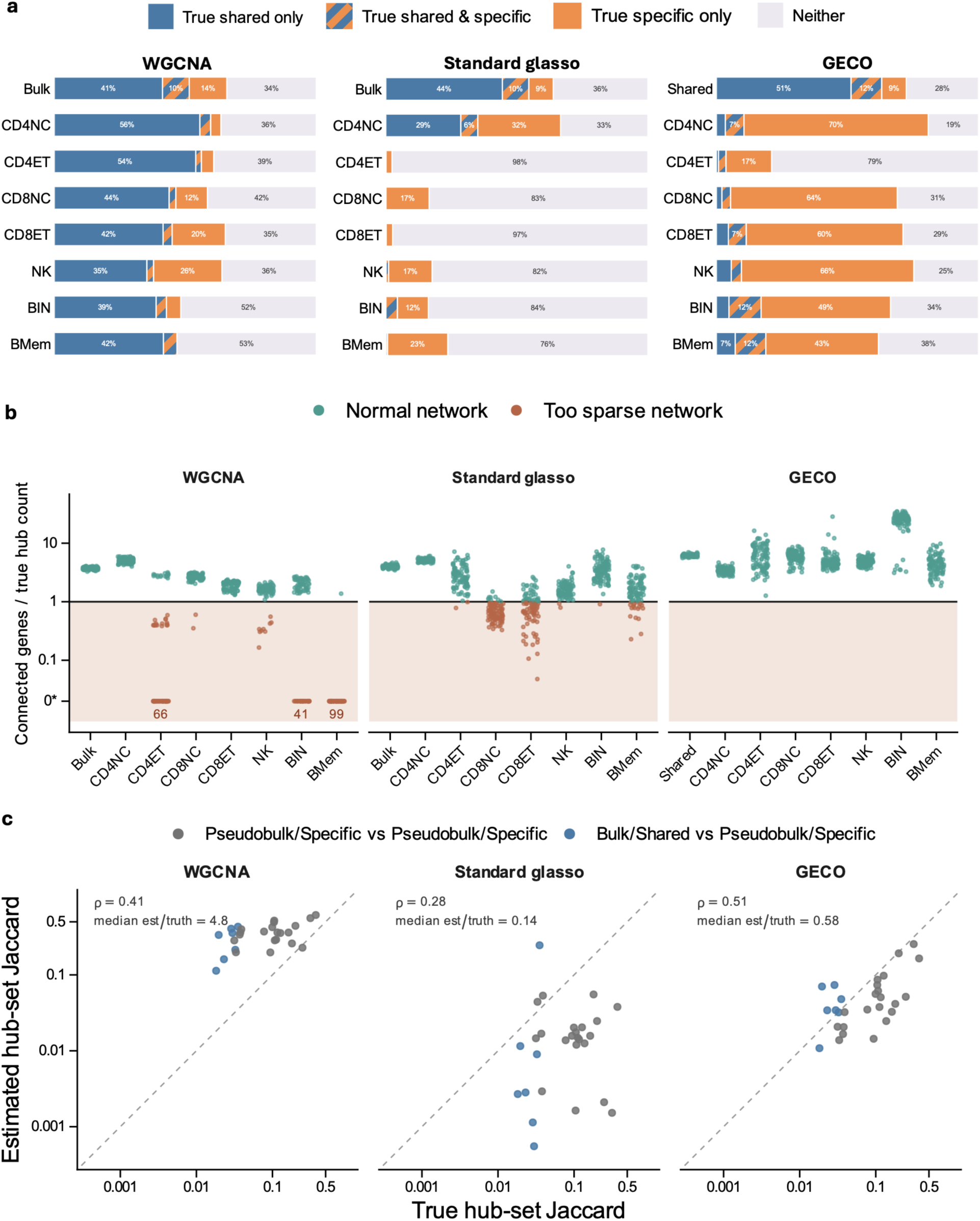
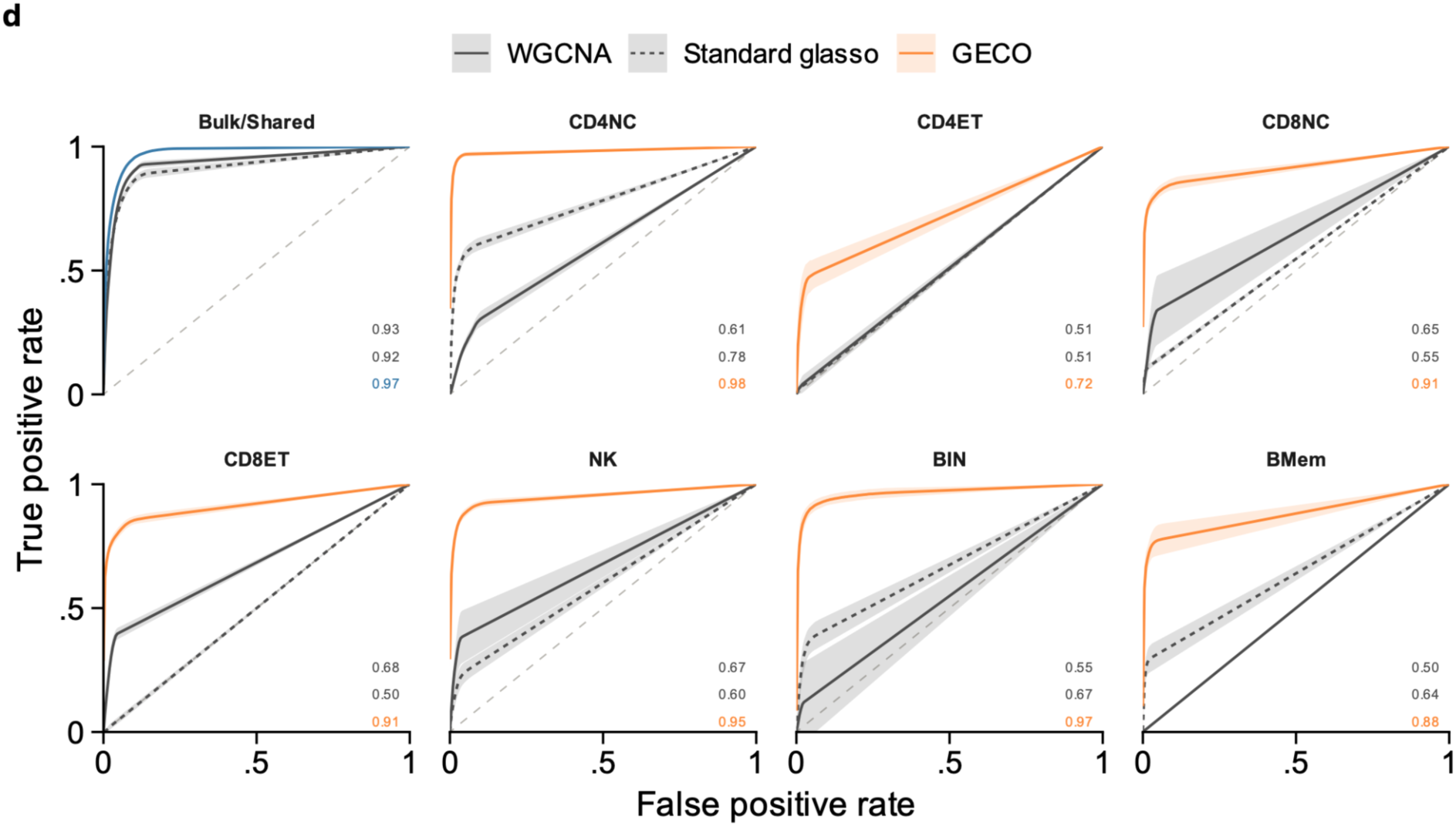
Further comparing GECO to standard coexpression networks in simulations. **(a)** Composition of hub sets predicted by WGCNA, standard glasso, and GECO. For each method and replicate, we selected the same number of genes as in the corresponding true hub set; when fewer connected genes were available, all connected genes were selected. Predicted hubs were classified as true shared hubs, true cell type-specific hubs, or “True shared & specific” if they belonged to both sets. For the “Shared”/“Bulk” networks, “True specific” denotes the union of true cell type-specific hubs across all cell types; for individual cell-type networks, it denotes the true specific hubs of that cell type. **(b)** Number of connected genes in the network inferred by each method in each replicate. 0* indicates no connected genes. WGCNA and standard glasso occasionally produced networks with fewer connected genes than the number of true hubs (234/800 and 175/800 network-replicates, respectively). **(c)** Cross-network hub-set overlap. WGCNA hubs showed high overlap across cell types, whereas standard glasso hubs showed little overlap. GECO recovered cross-cell-type hub overlap consistent with the ground truth. **(d)** Receiver operating characteristic (ROC) curves for ranking true hubs above non-hubs in each simulated network. Genes were ranked by kME for WGCNA and by degree for standard glasso and GECO. Genes that the method leaves unconnected are tied at the bottom of the ranking, so the curve is drawn straight across each such block, which is what averaging over all their possible orderings gives. Lines show mean ROC curves across 100 replicates, with shaded bands indicating ±1 s.d.; the dashed diagonal indicates chance performance. Values show mean per-replicate AUROC. GECO achieved higher AUROC in every network, indicating that its improved hub recovery was robust to the hub-selection threshold.

**Ext Data Fig 2.**
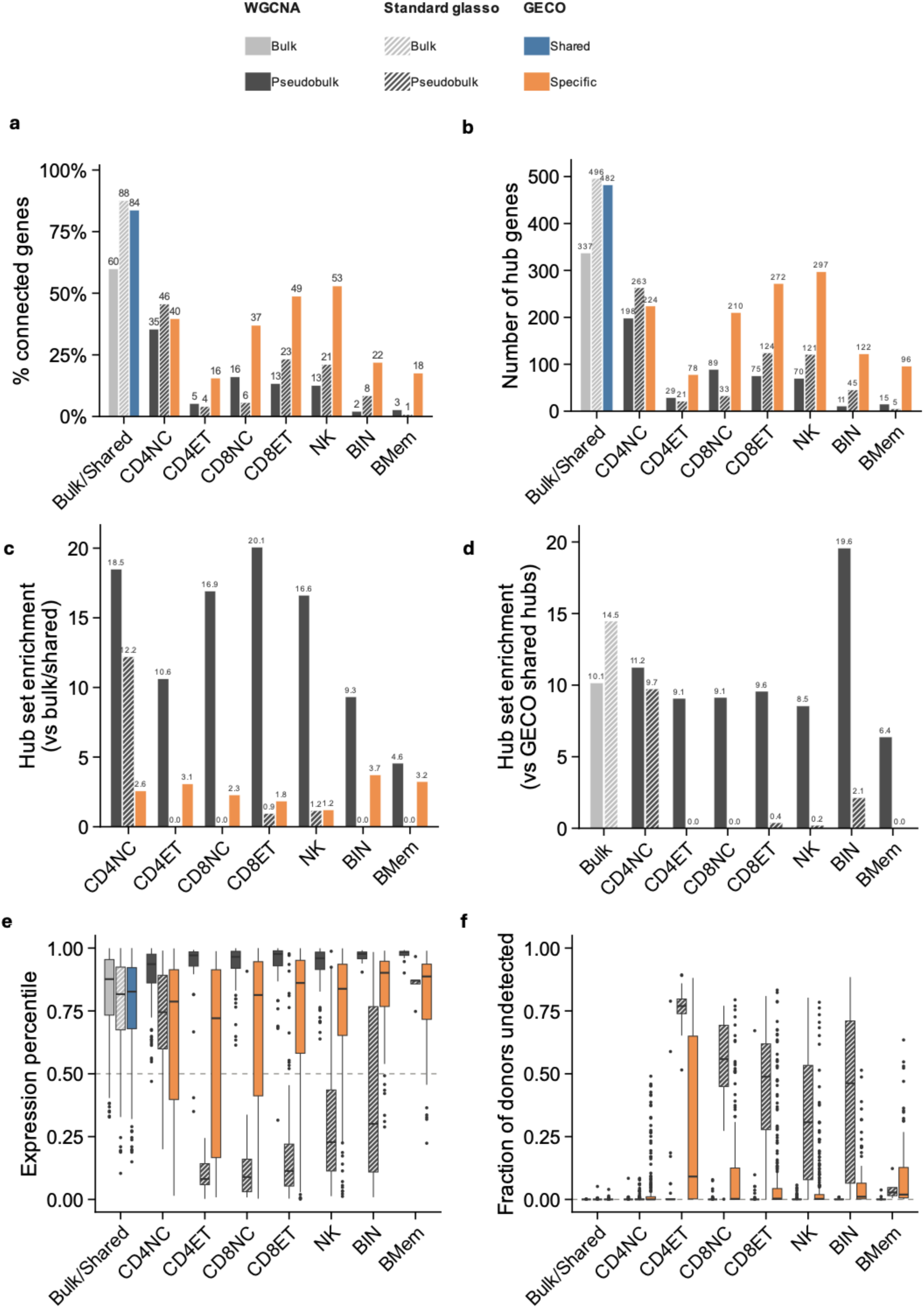
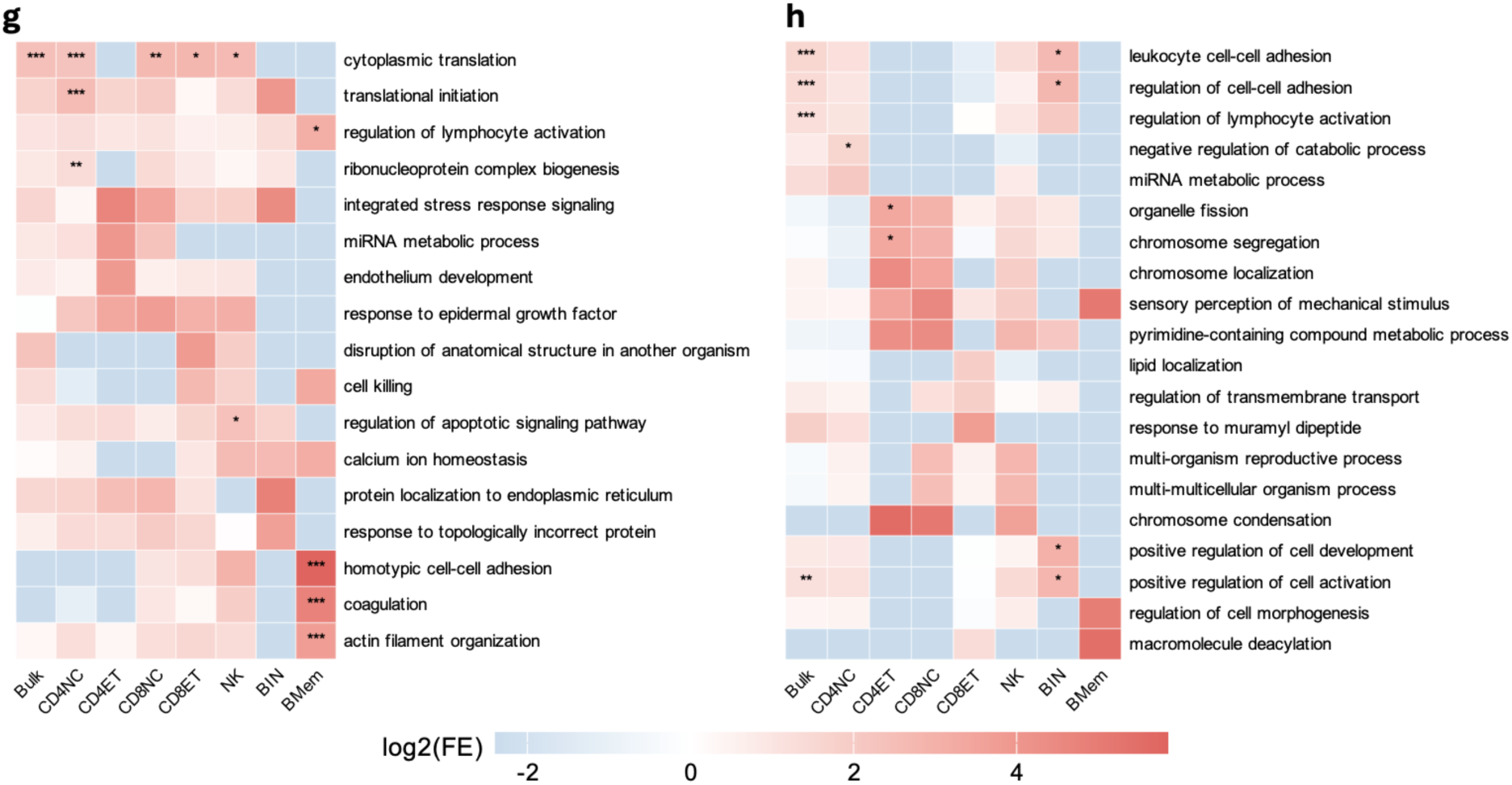
Further comparing GECO to standard coexpression networks in real data. **(a)** Proportion of connected genes in networks generated by WGCNA, standard glasso, and GECO. WGCNA and standard glasso produced particularly sparse networks in low-abundance cell types. **(b)** Number of hub genes identified by WGCNA, standard glasso, and GECO. **(c)** Fold enrichment of pseudobulk (specific) hubs among bulk (shared) hubs. WGCNA pseudobulk hubs were strongly enriched for bulk hubs. **(d)** Fold enrichment of WGCNA/standard glasso pseudobulk hubs among GECO shared hubs. WGCNA hubs, standard glasso bulk and CD4NC pseudobulk hubs were strongly enriched for GECO shared hubs. **(e)** Expression quantiles of hub genes within each cell type, with one point per hub. WGCNA hubs tended to be highly expressed, standard glasso hubs lowly expressed, and GECO hubs intermediate. **(f)** Fraction of donors in which each hub gene was undetected, with one point per hub. In five of seven cell types, standard glasso identified hubs undetected in 31-77% of donors, whereas WGCNA and GECO hubs were rarely undetected. **(g)** WGCNA and **(h)** standard glasso showed diffuse and inconsistent GO enrichment, dominated by broad metabolic processes or heterogeneous immune and cell-cycle terms, with limited coherent cell type specificity. We show the union of the top three GO terms from each hub set. ***: FDR < 0.001; **: FDR < 0.01; *: FDR < 0.05.

**Ext Data Fig 3.**
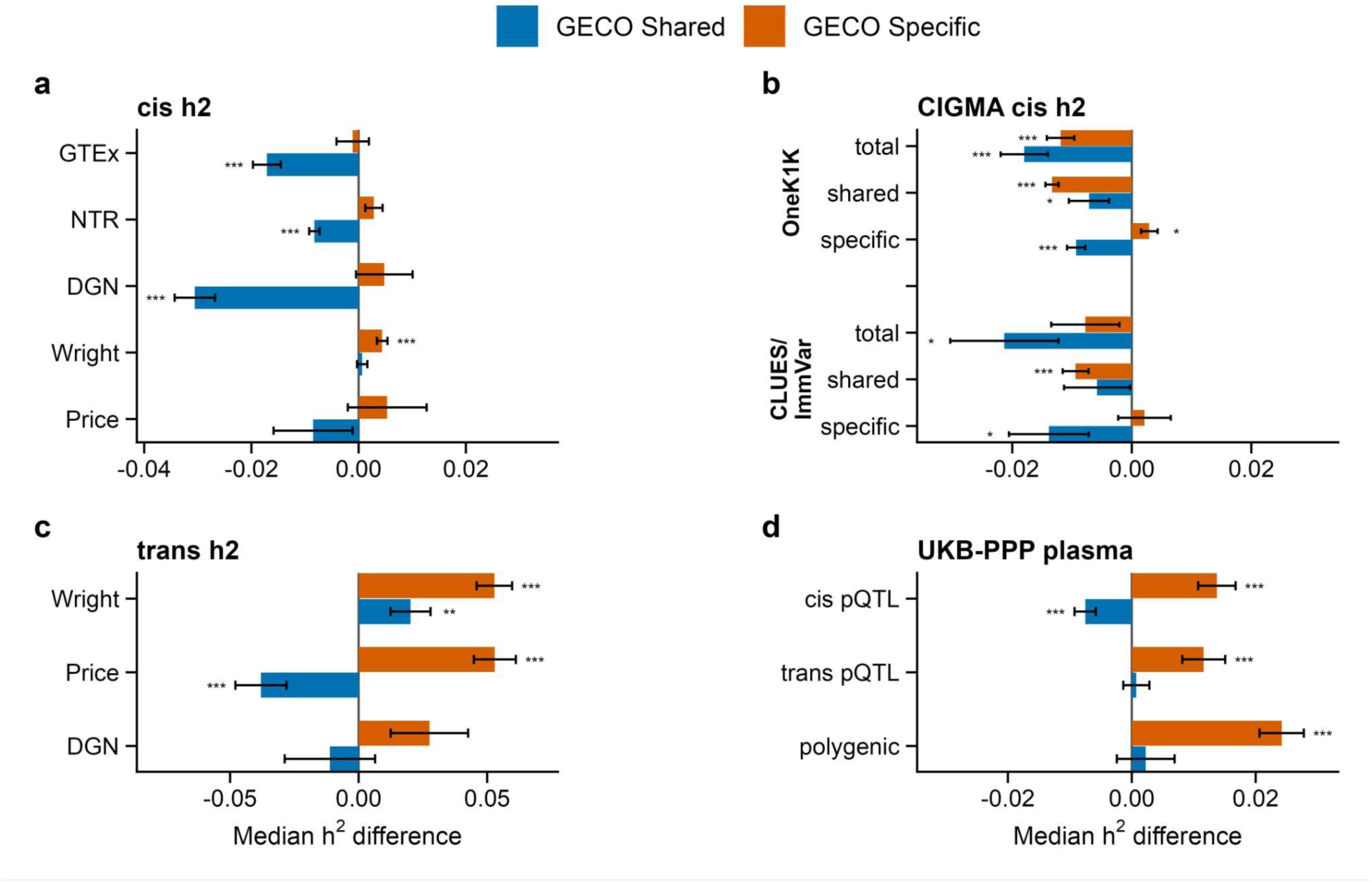
Estimates of molGene heritability in GECO hub genes. (a,c) *cis* and *trans* heritability estimates from whole blood gene expression from NTR (array from Wright et al. 2014, RNA-seq from Ouwens et al. 2020), Icelandic families (Price et al. 2011), DGN (Wheeler et al. 2016), and GTEx (The GTEx Consortium 2020). **(b)** Total, shared, and cell type-specific *cis* heritability estimates from CIGMA (Minhui Chen et al. 2026) in PBMCs from OneK1K (Yazar et al. 2022) and CLUES/ImmVar (Perez et al. 2022). **(d)** Variance explained by significant *cis* and *trans* pQTLs in UKB-PPP and remaining polygenic heritability explained by LDSC (Sun et al. 2023). Error bars indicate +/- 1 standard error. ***: p < 0.001; **: p < 0.01; *: p < 0.05.

**Ext Data Fig 4.**
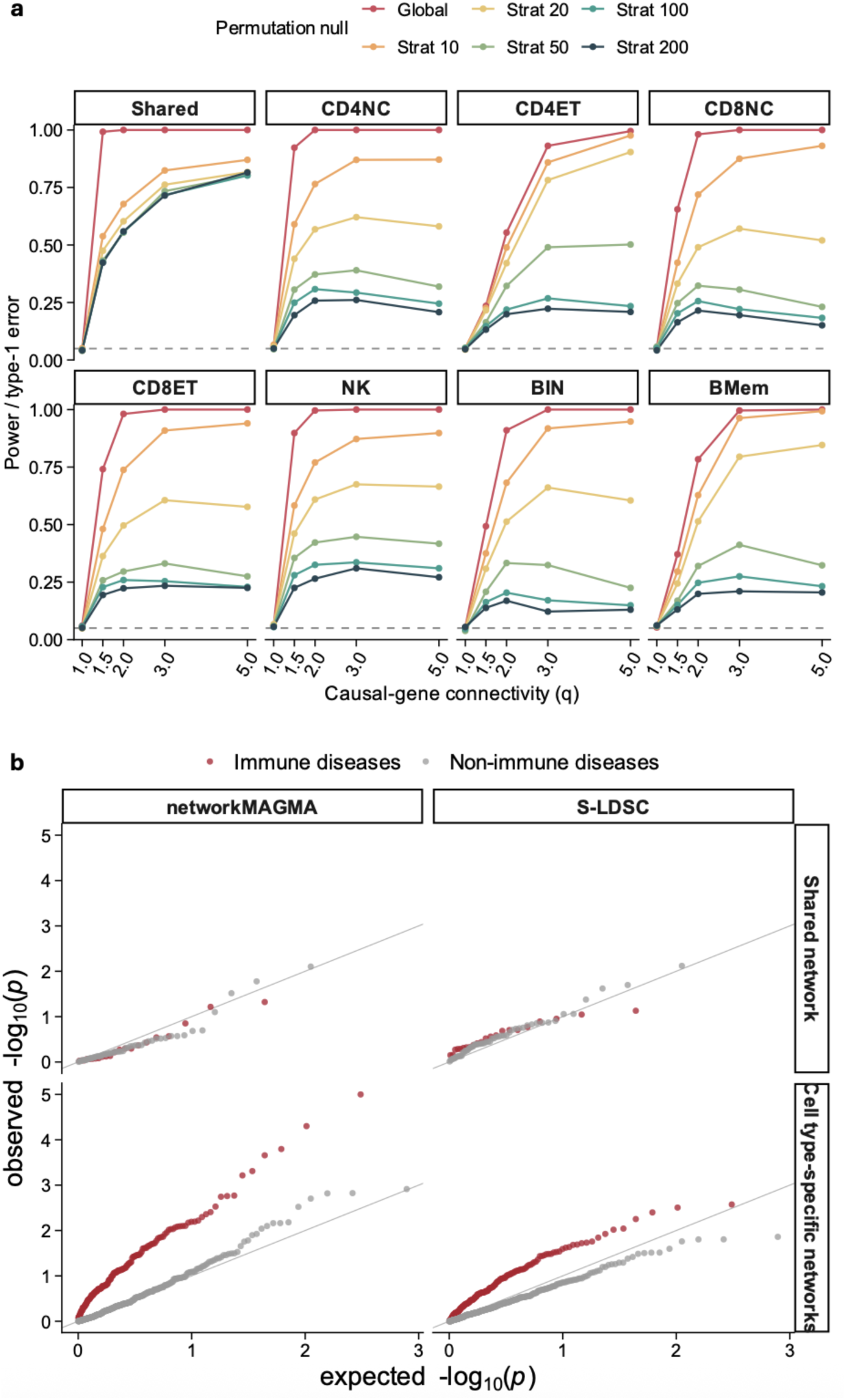
NetworkMAGMA in simulations and OneK1K. **(a)** Type I error and power of networkMAGMA in simulations. Using |partial correlation| weights, networkMAGMA maintained nominal type I error (q = 1) in each permutation null setting across all eight GECO networks. As causal genes formed increasingly connected modules, power under the global null approached 1.0 by q = 2 in all networks, whereas degree-stratified nulls had lower power that further decreased with finer binning. We therefore used the global null in real-data analyses. **(b)** Observed versus expected −log10 P values for enrichment of 78 FinnGen diseases in GECO networks, shown separately by method and network class. Points are colored by disease category. The grey line denotes the identity line (y = x). In cell type-specific networks, immune diseases showed marked deviation from the null for both methods, whereas non-immune diseases remained well calibrated, supporting disease-class-specific enrichment rather than global test-statistic inflation.

**Ext Data Fig 5.**
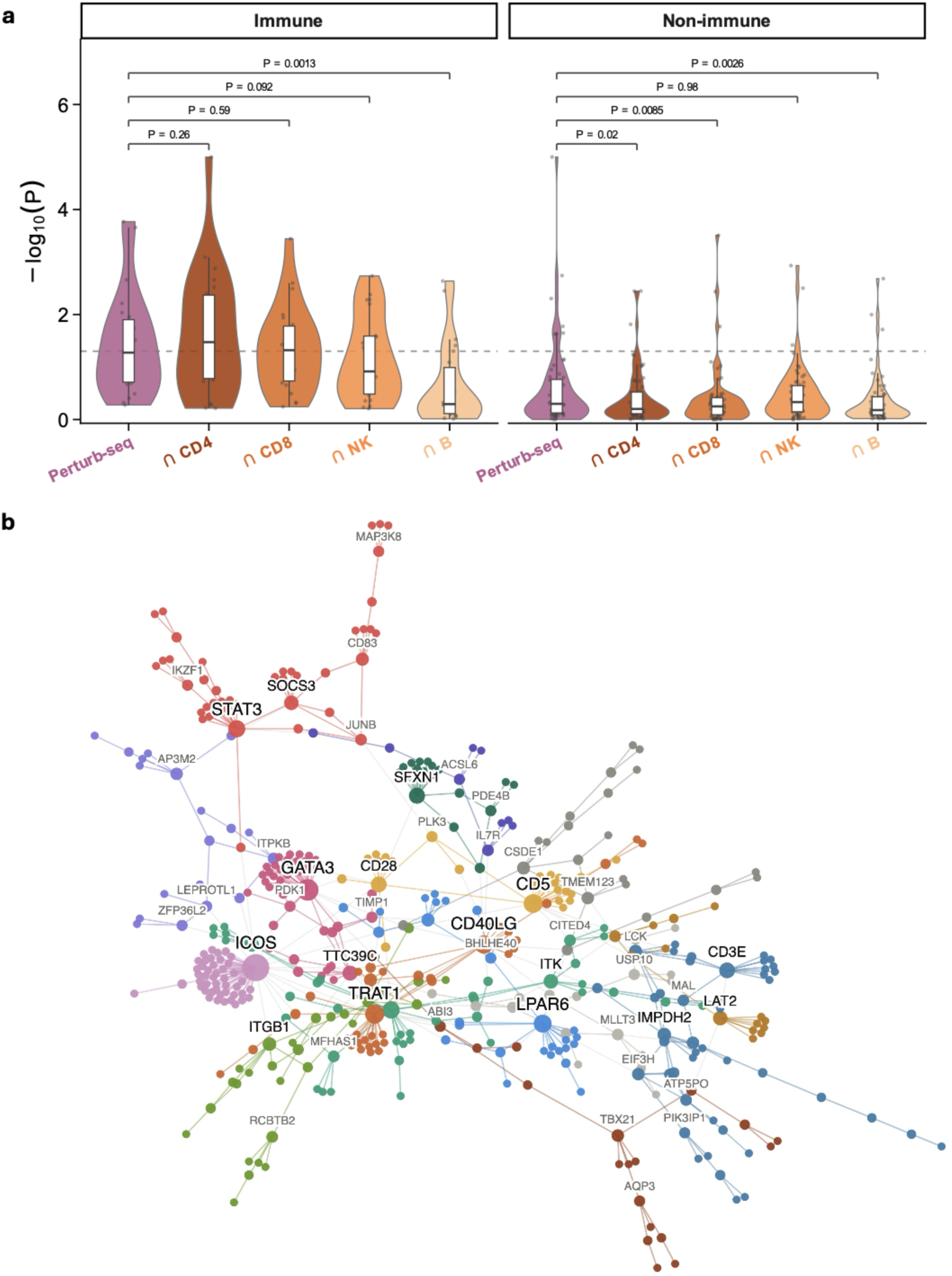
Intersecting GECO networks with perturb-seq network. **(a)** NetworkMAGMA analysis after intersecting the CD4 T cell perturb-seq network with GECO CD4-, CD8-, NK-, and B-specific networks. P values are from two-sided Wilcoxon tests. Overlap with the GECO CD4-specific network increased immune-disease GWAS gene-pair enrichment and reduced non-immune enrichment, whereas overlap with the GECO B-specific network reduced both. **(b)** Largest connected component of the overlap between the GECO CD4-specific and CD4 T cell perturb-seq networks (450 genes, 526 edges). Nodes are colored by Louvain community; node and label size scale with degree. The overlap forms a highly modular network with lineage-relevant hubs and communities.

**Ext Data Fig 6.**
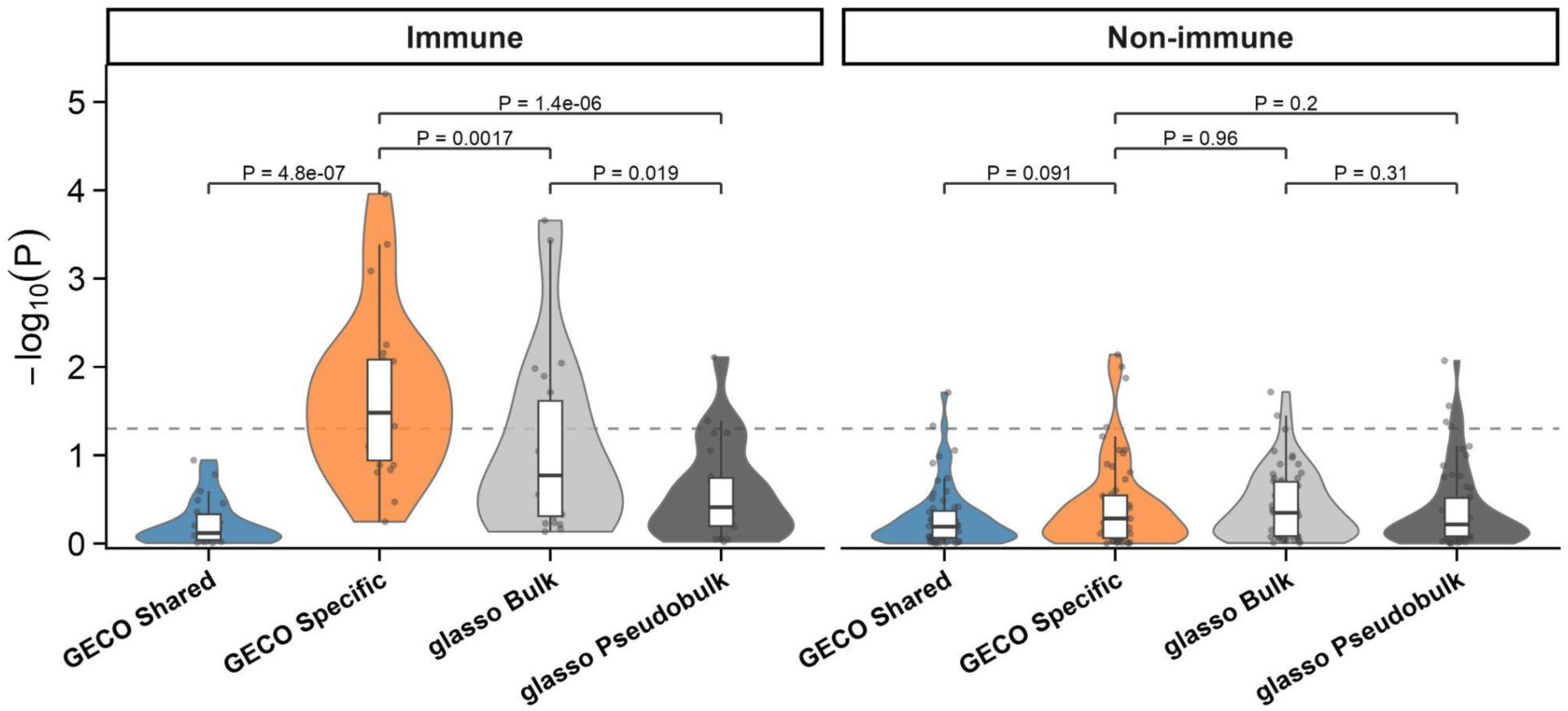
NetworkMAGMA for GECO shared/specific network, standard glasso bulk/pseudobulk network. The GECO specific network showed stronger enrichment for immune disease GWAS gene pairs than both the bulk and pseudobulk glasso networks. To make these networks comparable, we took the union of seven GECO specific networks as the GECO Specific network and the union of seven standard glasso pseudobulk networks as the glasso Pseudobulk network.

**Ext Data Fig 7.**
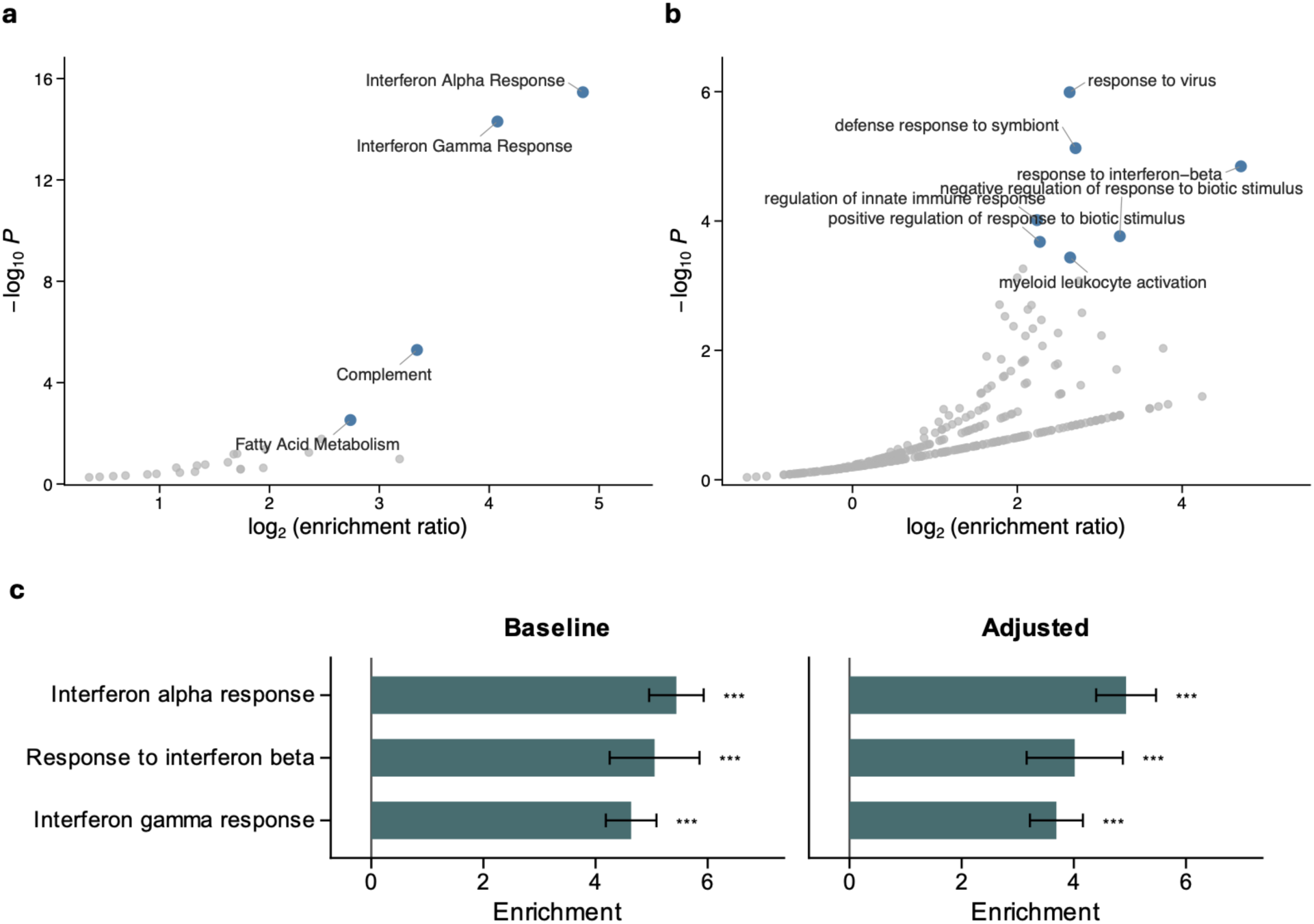
Enrichment of BAFF neighbors in the cM-specific network. **(a, b)** Enrichment of BAFF network neighbors in (**a**) Hallmark gene sets and (**b**) Gene Ontology (GO) terms. P values were calculated using hypergeometric tests. Blue dots indicate significant enrichment (FDR < 0.05). **(c)** Enrichment of interferon-related gene sets assessed using logistic regression, consistent with other enrichment analyses. Baseline represents the primary analysis, whereas Adjusted represents the sensitivity analysis controlling for mean expression, expression variability, gene length, and GC content. Error bars indicate ±1 standard error. The x-axis shows log2 odds ratios (Methods). ***: p < 0.001; **: p < 0.01; *: p < 0.05.

