## supplementary figures for "Cell type-specific coexpression networks funnel genetic risk into core genes"

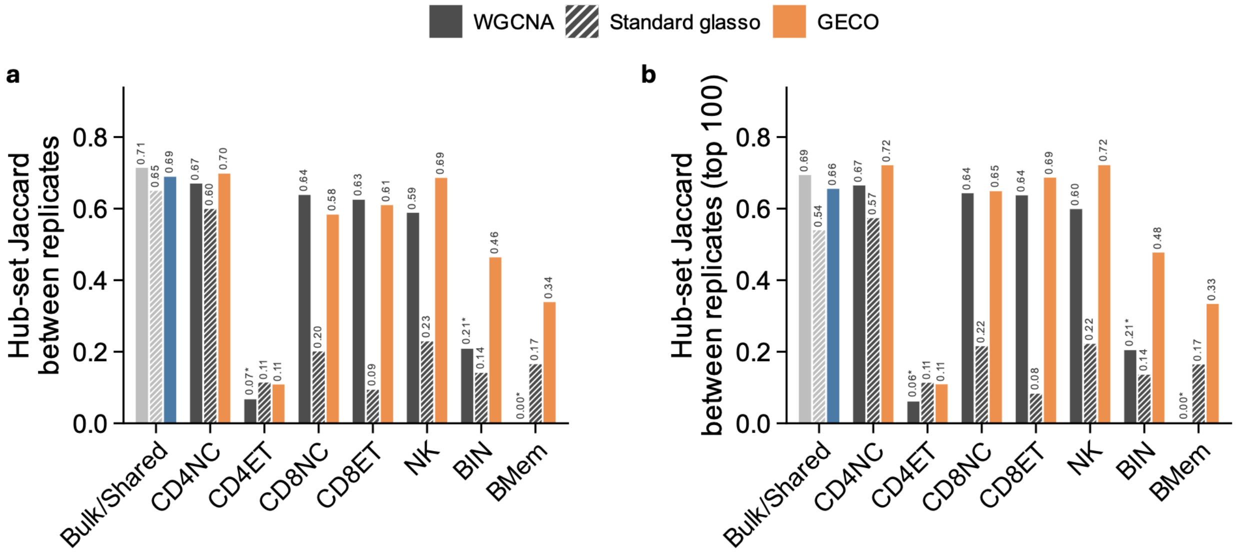


**Supplementary Figure 1. Reproducibility of predicted hub sets in simulated data. (a)** Mean Jaccard index between hub sets predicted across independent replicates of the same network. Replicates yielding no hubs were assigned a Jaccard index of 0; bars marked * include such replicates. Standard glasso showed high reproducibility in the bulk network (0.65) but low reproducibility in six of seven cell types (0.10-0.23). **(b)** Same analysis with the number of predicted hubs fixed at 100 for every network and method.

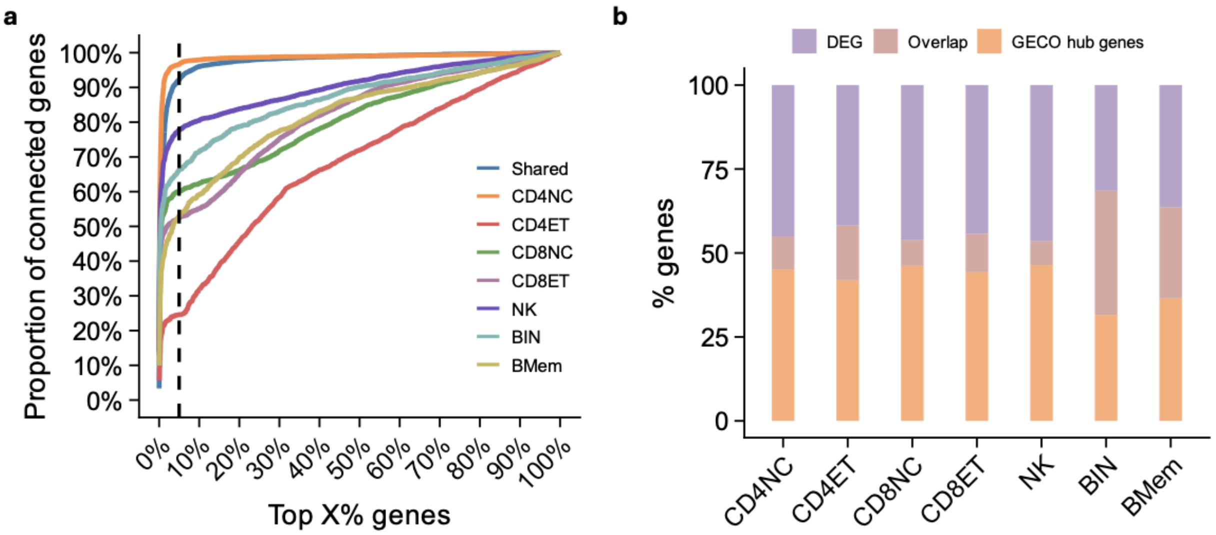


**Supplementary Figure 2. GECO network summary in OneK1K. (a)** Proportion of non-isolated genes connected to the top-ranked genes by degree in GECO shared and cell type-specific networks. A small fraction of high-degree genes connected to a large proportion of the network. Hubs (top 5% by degree) were connected to 61.6% (mean across seven cell type-specific networks) to 92.3% (shared network) of non-isolated genes. **(b)** GECO cell type-specific hubs partially overlapped with the most differentially expressed genes (DEGs; ranked by the variance of mean expression across cell types), with overlaps ranging from 13.5% to 54.1%. For each network, we selected the same number of DEGs as hub genes.

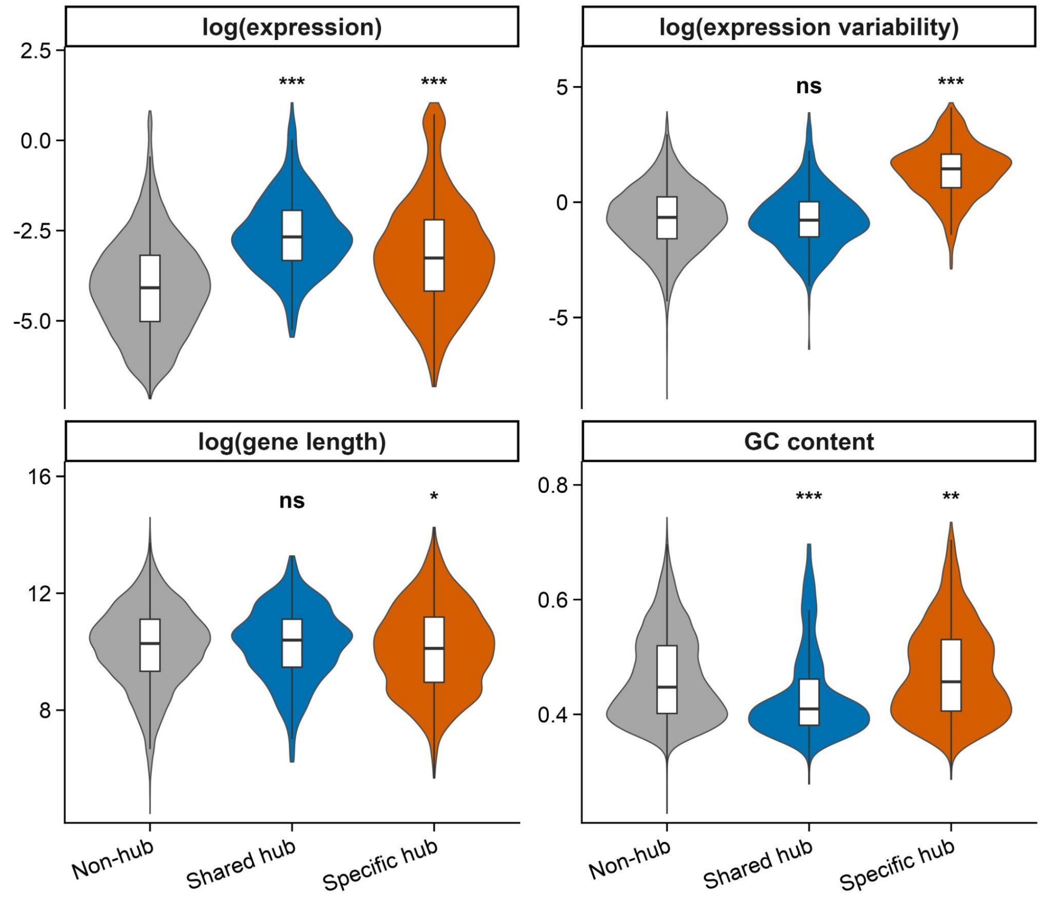


**Supplementary Figure 3. Covariates adjusted for in the sensitivity analysis.** Compared with non-hubs, shared hubs had higher overall expression levels and lower GC content, whereas cell type-specific hubs had higher overall expression levels, greater expression variability, shorter gene length, and higher GC content. ***: p < 0.001; **: p < 0.01; *: p < 0.05.

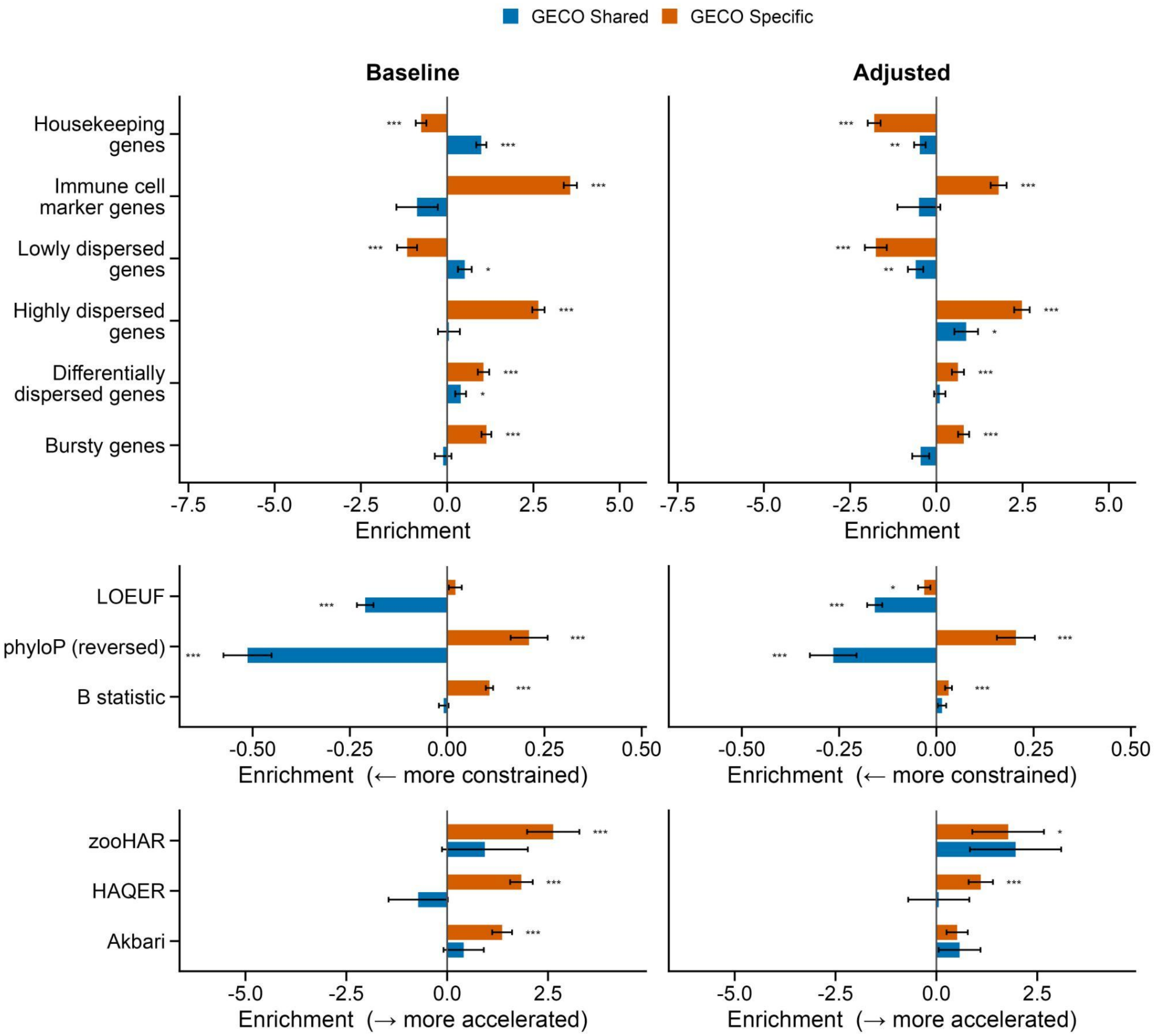


**Supplementary Figure 4. Primary and sensitivity analyses of benchmark gene sets and evolutionary features in OneK1K.** Baseline: primary analysis; Adjusted: sensitivity analysis controlling for mean expression, expression variability, gene length, and GC content. For the six evolutionary features (LOEUF, phyloP, B statistic, zooHAR, HAQER, and Akbari), sensitivity analyses additionally adjusted for recombination rate and number of exons. ***: $p < 0.001$; **: $p < 0.01$; *: $p < 0.05$. Error bars indicate ±1 standard error. For binary annotations, the x-axis shows log2 odds ratios; for continuous annotations, it shows regression coefficients (Methods).

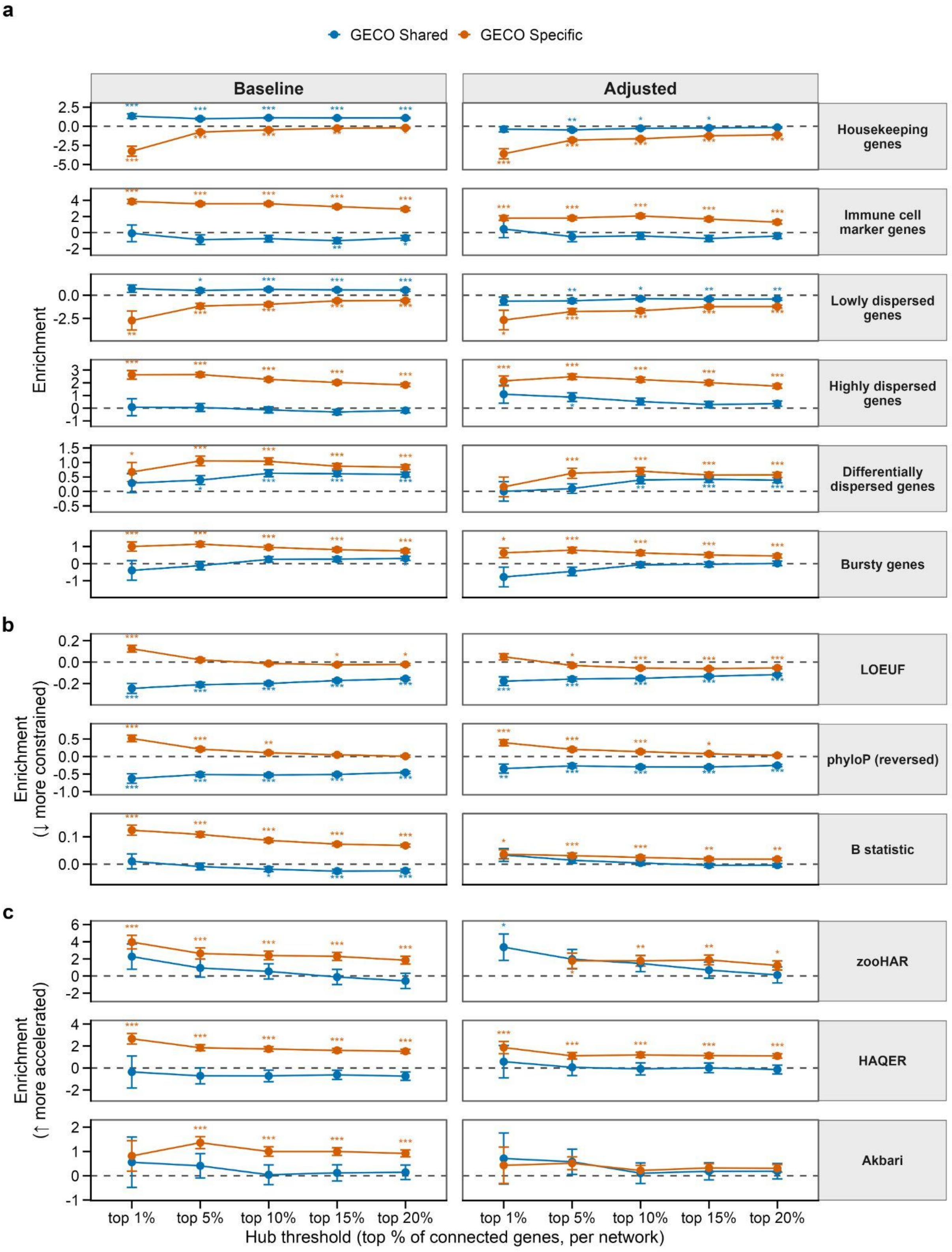


**Supplementary Figure 5. Baseline and sensitivity analyses of benchmark gene sets (a) and evolutionary features (b,c) in OneK1K with different hub thresholds.** Baseline: primary analysis;

Adjusted: sensitivity analysis controlling for mean expression, expression variability, gene length, and GC content. For the six evolutionary features (LOEUF, phyloP, B statistic, zooHAR, HAQER, and Akbari), sensitivity analyses additionally adjusted for recombination rate and number of exons. ***: $p < 0.001$; **: $p < 0.01$; *: $p < 0.05$. Error bars indicate ±1 standard error. For binary annotations, the x-axis shows log2 odds ratios; for continuous annotations, it shows regression coefficients (Methods).

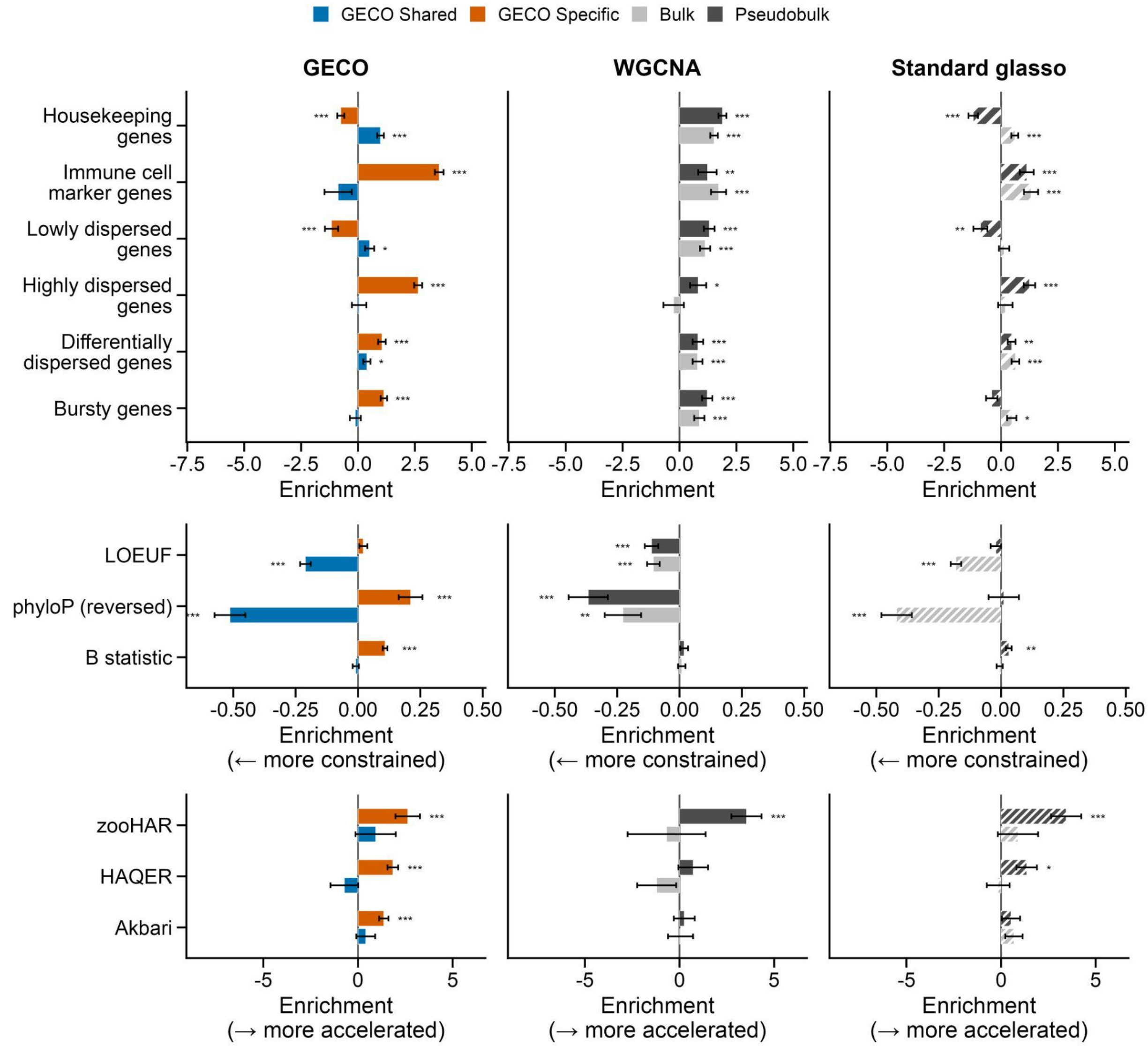


**Supplementary Figure 6. Enrichment of hubs identified by GECO, WGCNA, and standard glasso for benchmark gene sets and evolutionary features.** ***: $p < 0.001$; **: $p < 0.01$; *: $p < 0.05$. Error bars indicate ±1 standard error. For binary annotations, the x-axis shows log2 odds ratios; for continuous annotations, it shows regression coefficients (Methods).

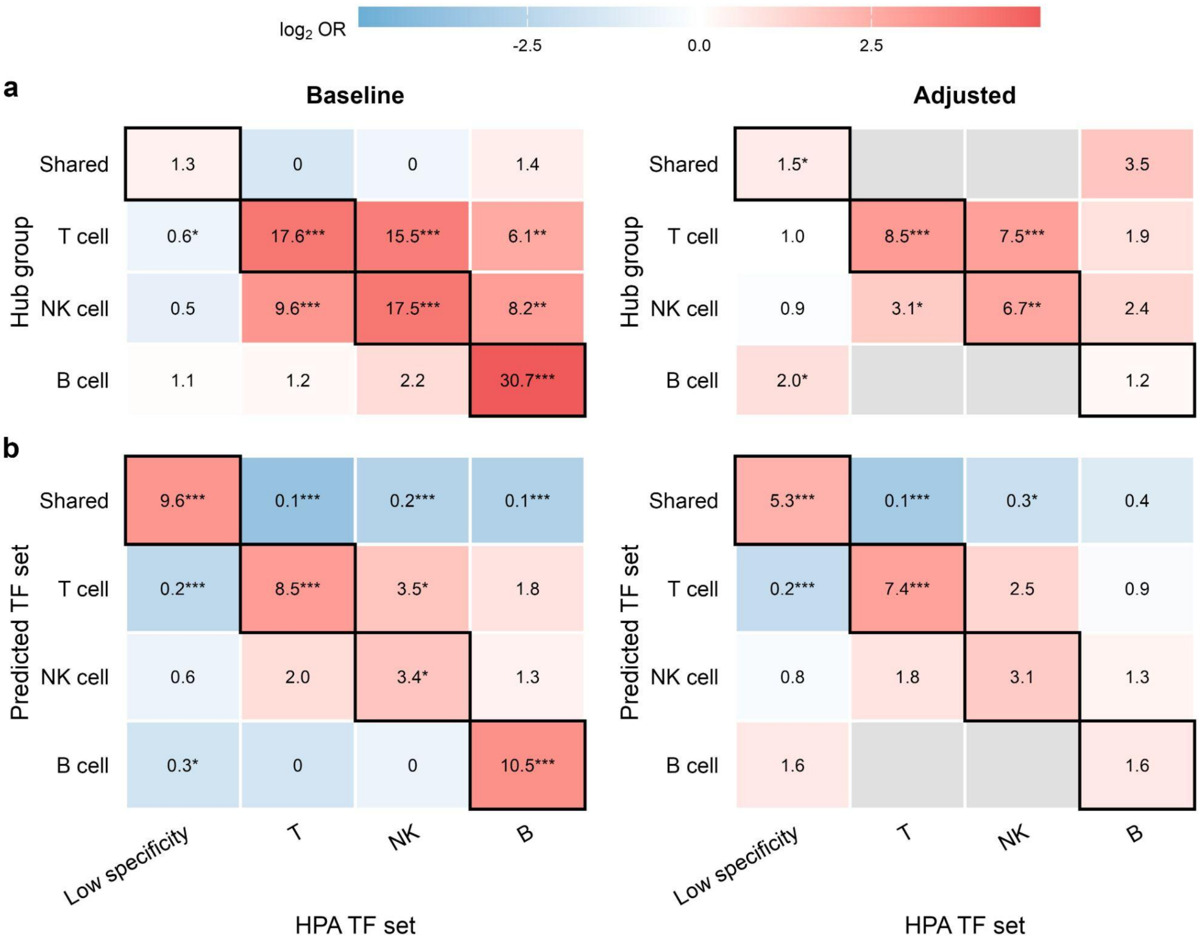


**Supplementary Figure 7. Primary and sensitivity analyses of TF enrichment (a) and TF lineage prediction (b) in OneK1K.** Baseline: primary analysis; Adjusted: sensitivity analysis controlling for mean expression, expression variability, gene length, and GC content. Grey cells indicate non-estimable comparisons: the hub group and TF set share no genes, resulting in complete separation in the covariate-adjusted model and an unbounded odds ratio. ***: $p < 0.001$; **: $p < 0.01$; *: $p < 0.05$.

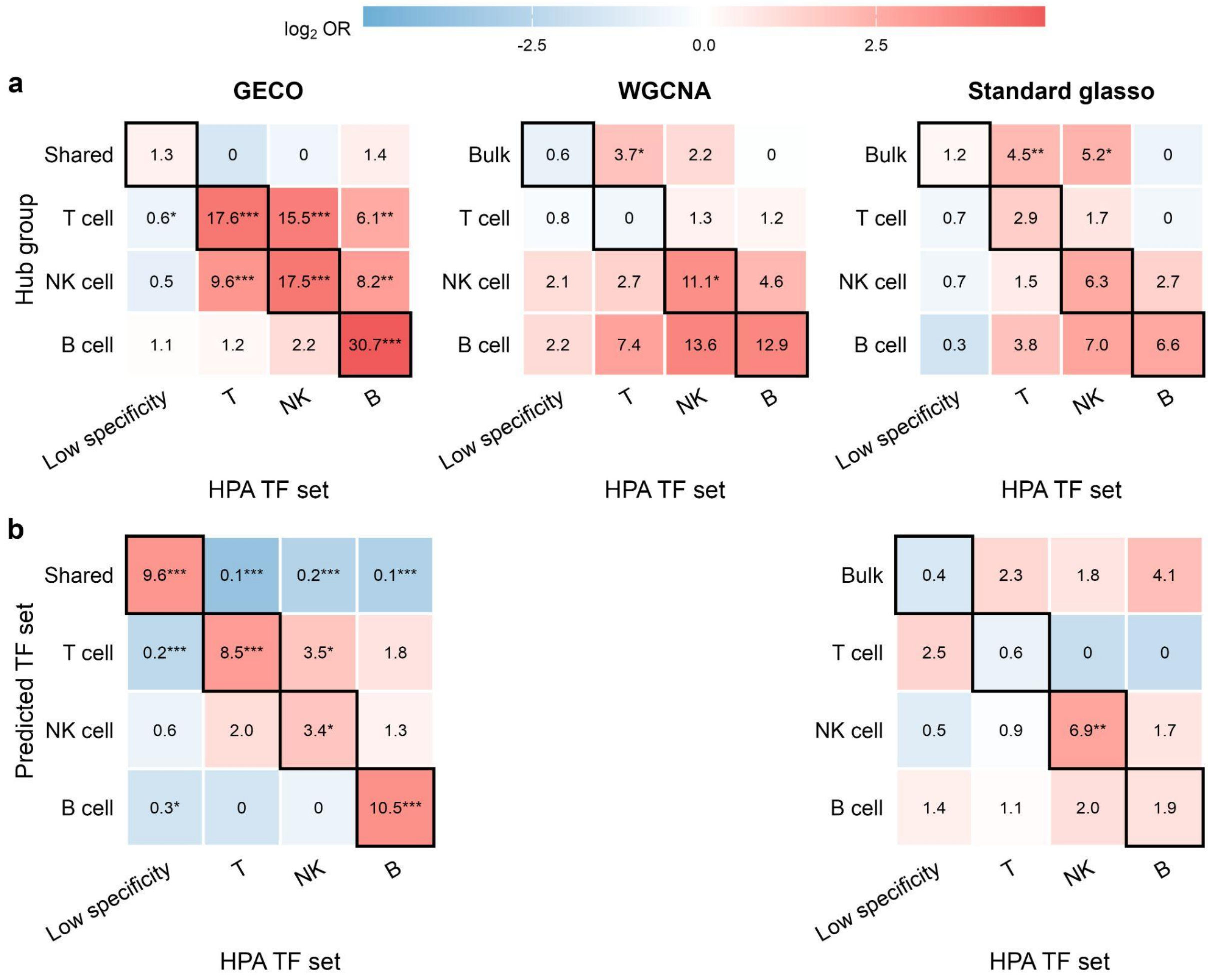


**Supplementary Figure 8. TF enrichment tests for GECO, WGCNA, and standard glasso (a), and TF lineage prediction tests for GECO and standard glasso (b) in OneK1K.** WGCNA was excluded from the lineage prediction test because kME is calculated within individual modules and is not directly comparable across networks. ***: $p < 0.001$; **: $p < 0.01$; *: $p < 0.05$.

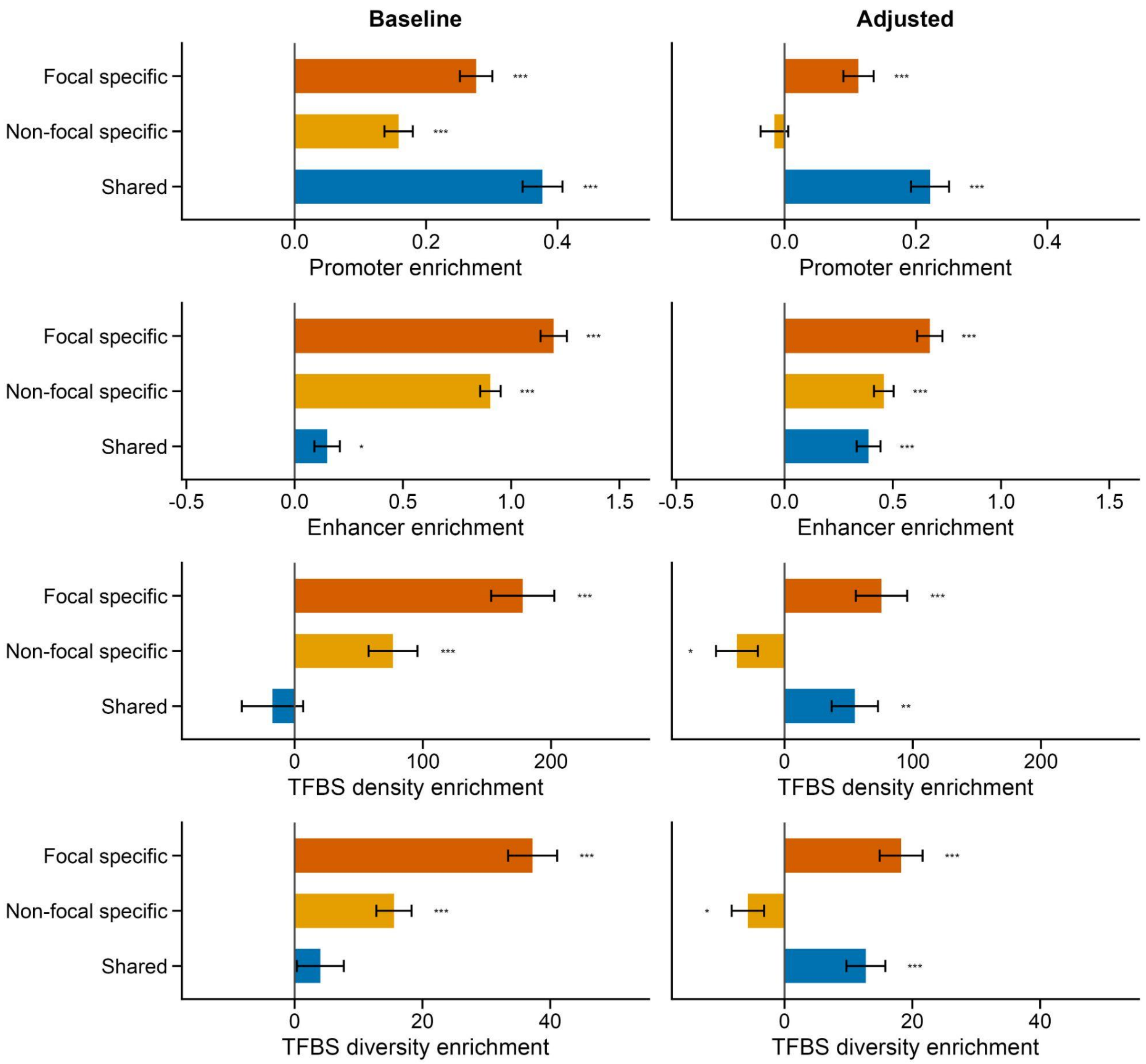


**Supplementary Figure 9. Primary and sensitivity analyses of promoter, enhancer, TFBS density, and TFBS diversity enrichment in OneK1K.** Baseline: primary analysis; Adjusted: sensitivity analysis controlling for mean expression, expression variability, gene length, and GC content. ***: $p < 0.001$; **: $p < 0.01$; *: $p < 0.05$. Error bars indicate ±1 standard error. The x-axis shows regression coefficients (Methods).

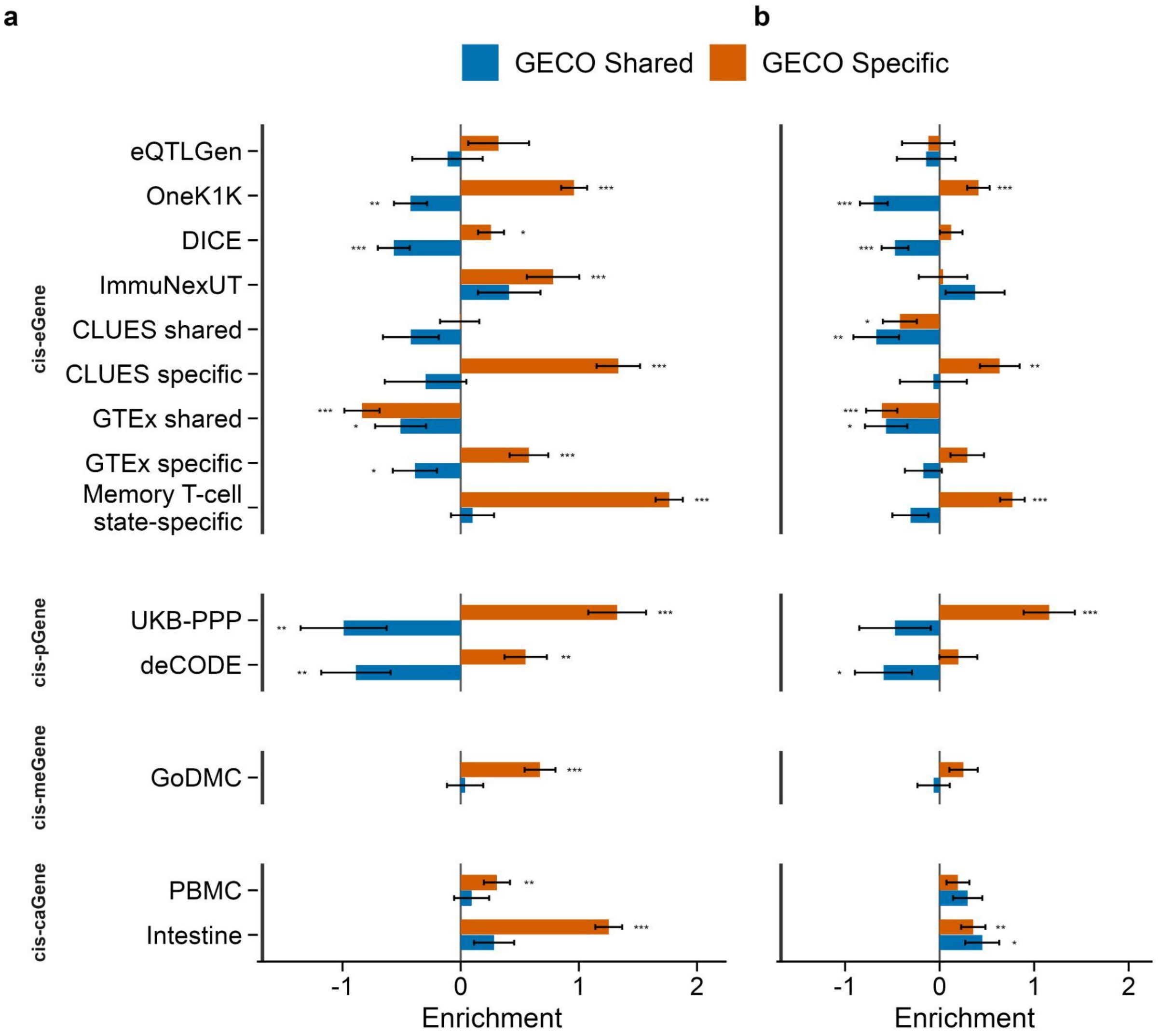


**Supplementary Figure 10. Primary and sensitivity analyses of *cis*-molGene enrichment in OneK1K.** **(a)** Baseline analysis. **(b)** Sensitivity analysis controlling for mean expression, expression variability, gene length, and GC content. ***: $p < 0.001$; **: $p < 0.01$; *: $p < 0.05$. Error bars indicate ±1 standard error. The x-axis shows log2 odds ratios (Methods).

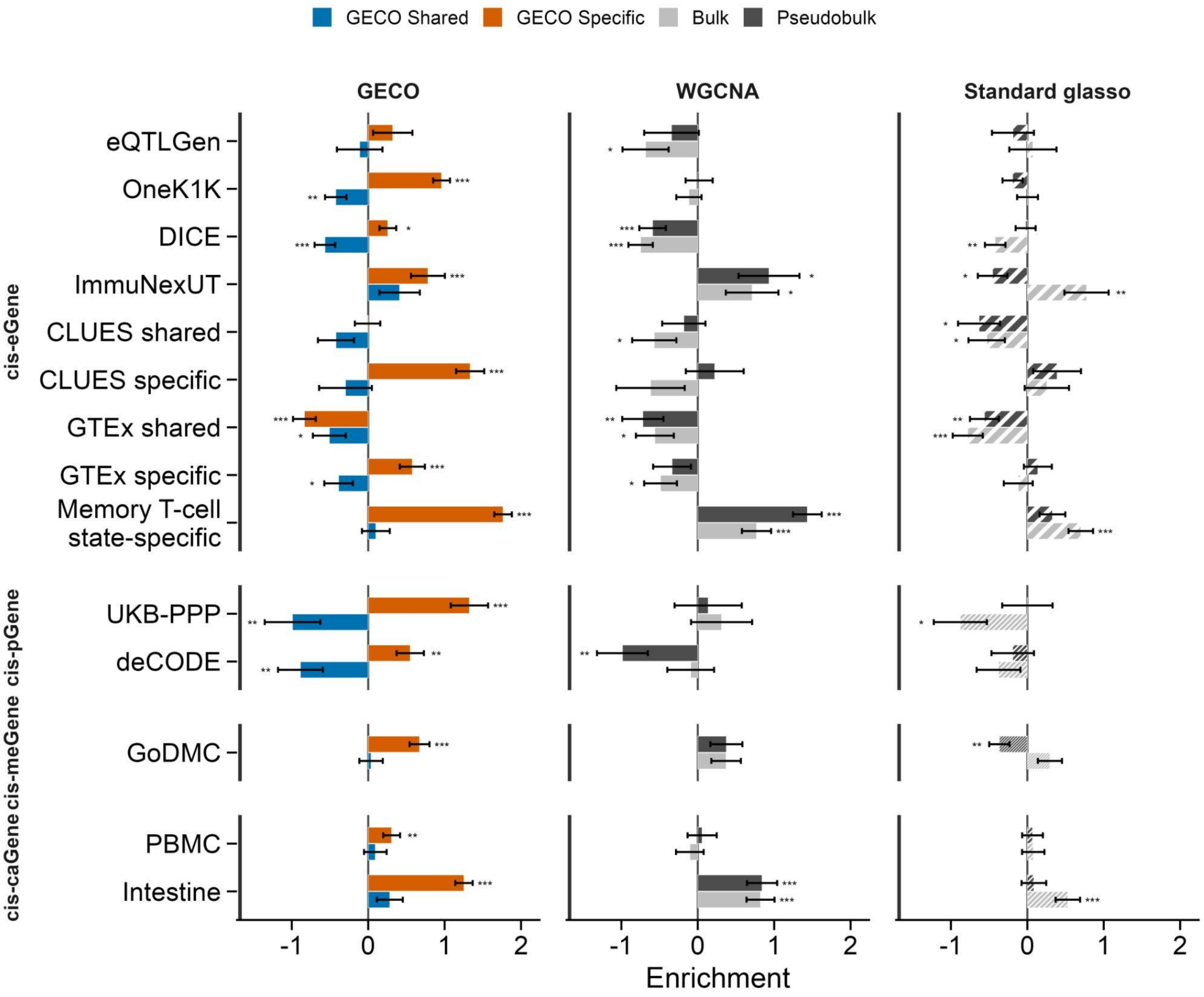


**Supplementary Figure 11. *cis*-molGene enrichment of hubs identified by GECO, WGCNA, and standard glasso in OneK1K.** Baseline: primary analysis; Adjusted: sensitivity analysis controlling for mean expression, expression variability, gene length, and GC content. ***: $p < 0.001$; **: $p < 0.01$; *: $p < 0.05$. Error bars indicate ±1 standard error. The x-axis shows log2 odds ratios (Methods).

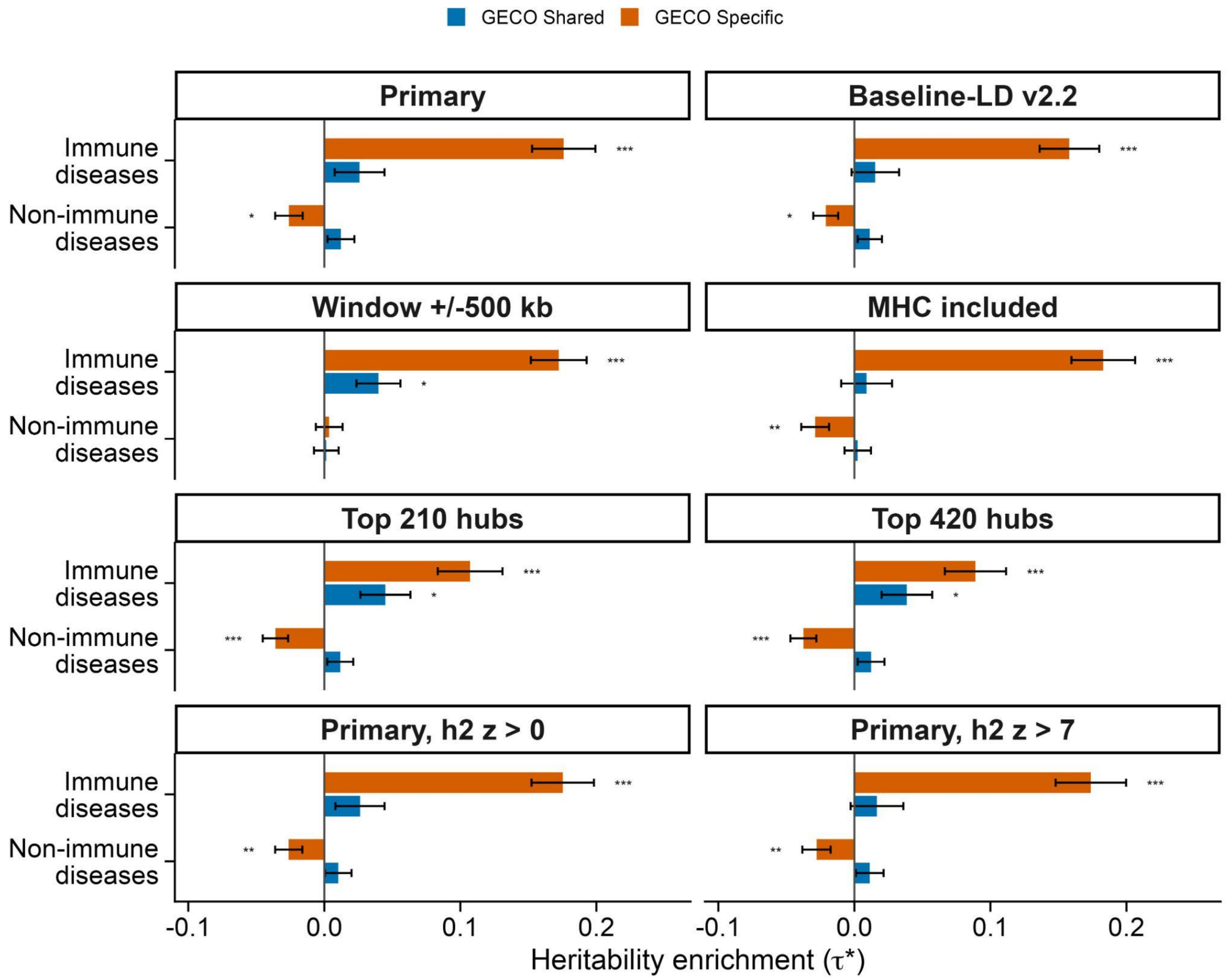


**Supplementary Figure 12. Sensitivity analyses of s-LDSC in OneK1K.** We assessed the robustness of heritability enrichment by using the baseline v2.2 model, including the MHC region, extending the cis window around hubs from ±100 kb to ±500 kb, restricting hubs to the top 210 or 420 genes, and changing the h2 z-score threshold to 0 or 7. ***: $p < 0.001$; **: $p < 0.01$; *: $p < 0.05$. Error bars indicate ±1 standard error.

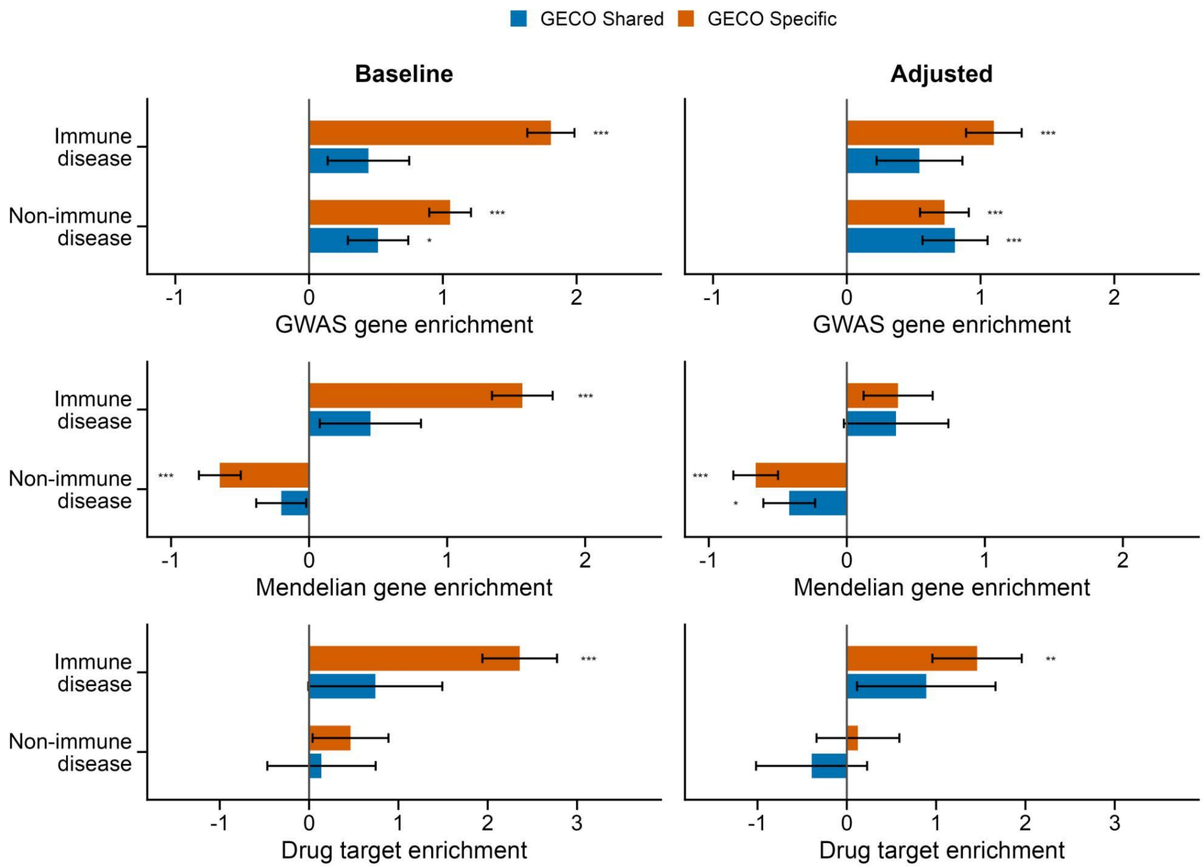


**Supplementary Figure 13. Primary and sensitivity analyses of GWAS gene, Mendelian gene, and approved drug target gene enrichment in OneK1K.** Baseline: primary analysis; Adjusted: sensitivity analysis controlling for mean expression, expression variability, gene length, and GC content. ***: $p < 0.001$; **: $p < 0.01$; *: $p < 0.05$. Error bars indicate ±1 standard error. The x-axis shows log2 odds ratios (Methods).

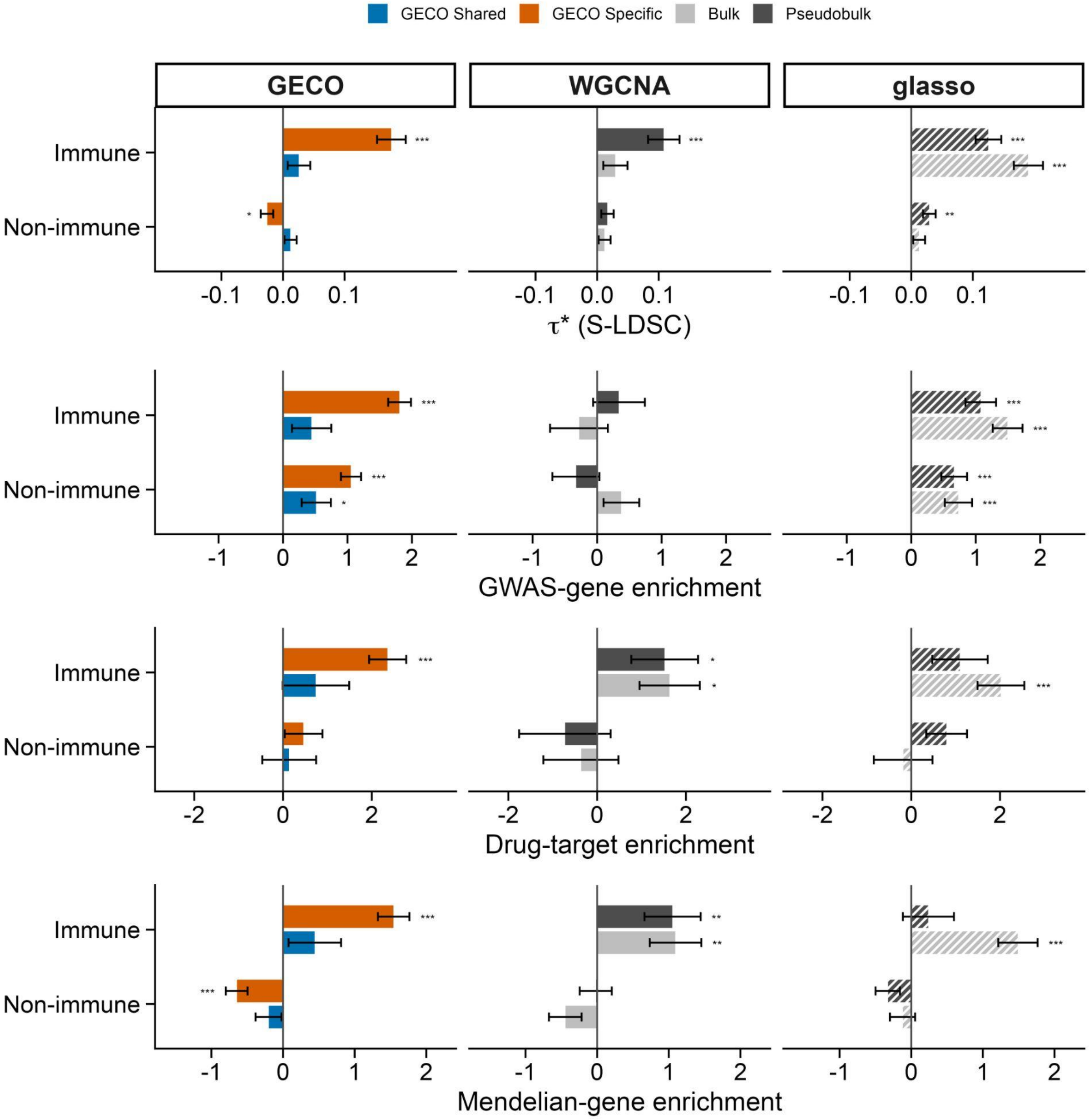


**Supplementary Figure 14. Heritability, GWAS gene, Mendelian gene, and approved drug target gene enrichment of hubs identified by GECO, WGCNA, and standard glasso in OneK1K.** ***: $p < 0.001$; **: $p < 0.01$; *: $p < 0.05$. Error bars indicate ±1 standard error. The x-axis shows log2 odds ratios for all analyses except heritability (Methods).

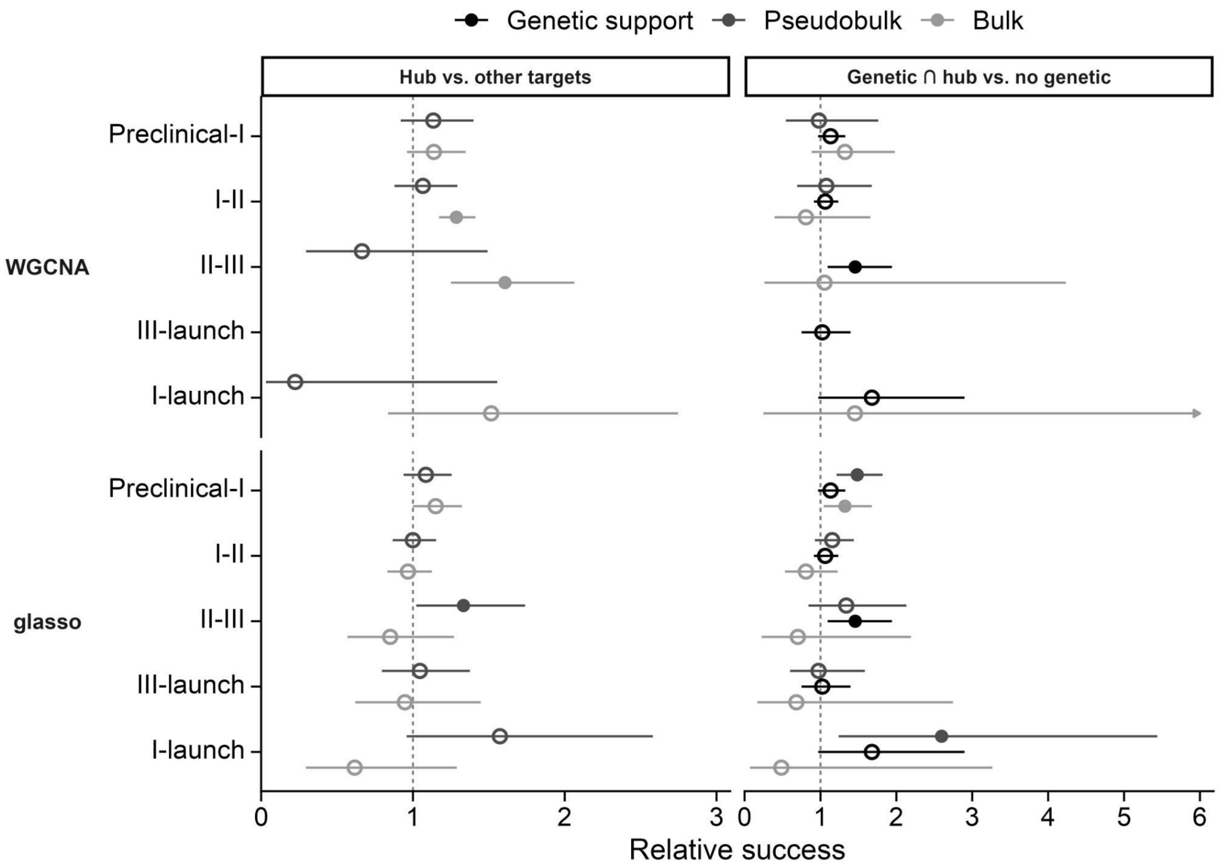


**Supplementary Figure 15. Relative success of target-indication pairs in immune indications from Citeline Pharmaprojects.** Rows show progression of drug targets through phases. Panels show relative success for bulk and pseudobulk hub genes (left) and for the intersection of indication-matched genetically supported genes as in Minikel et al. (OMIM or GWAS, right). These analyses restrict to genes included in GECO. Bars show 95% CI; filled points significantly differ from 1; and stars compare the black and grey/light grey estimates. ***: $p < 0.001$; **: $p < 0.01$; *: $p < 0.05$. WGCNA and standard glasso hubs cannot further improve the relative success of indication-matched genetically supported genes (all $p>0.05$).

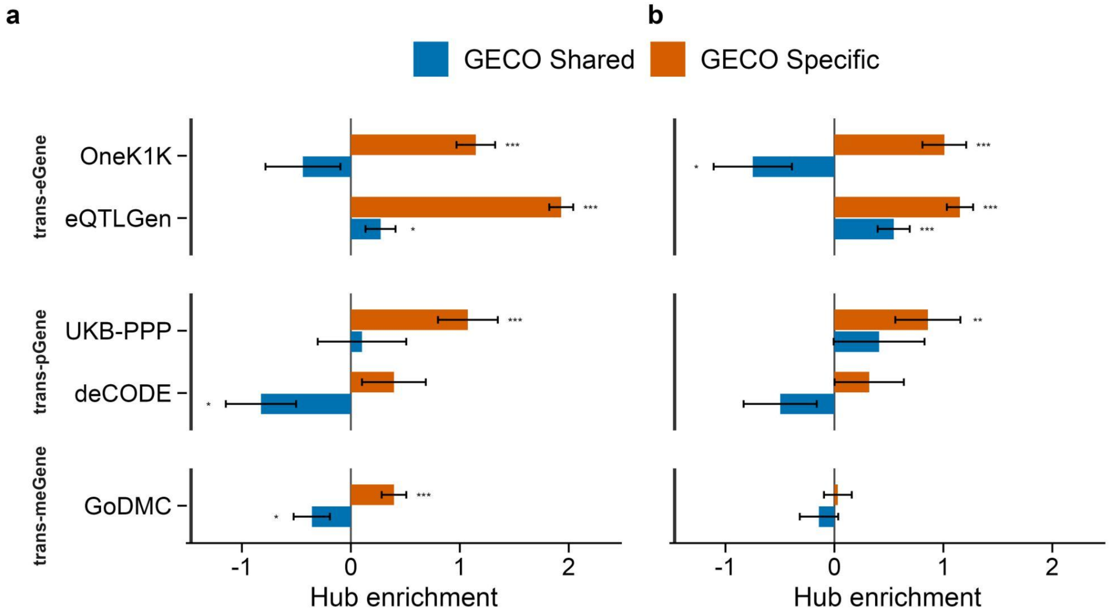


**Supplementary Figure 16. Primary and sensitivity analyses of *trans*-molGene enrichment in OneK1K. (a)** Baseline analysis. **(b)** Sensitivity analysis controlling for mean expression, expression variability, gene length, and GC content. ***: p < 0.001; **: p < 0.01; *: p < 0.05. Error bars indicate ±1 standard error. The x-axis shows log2 odds ratios (Methods).

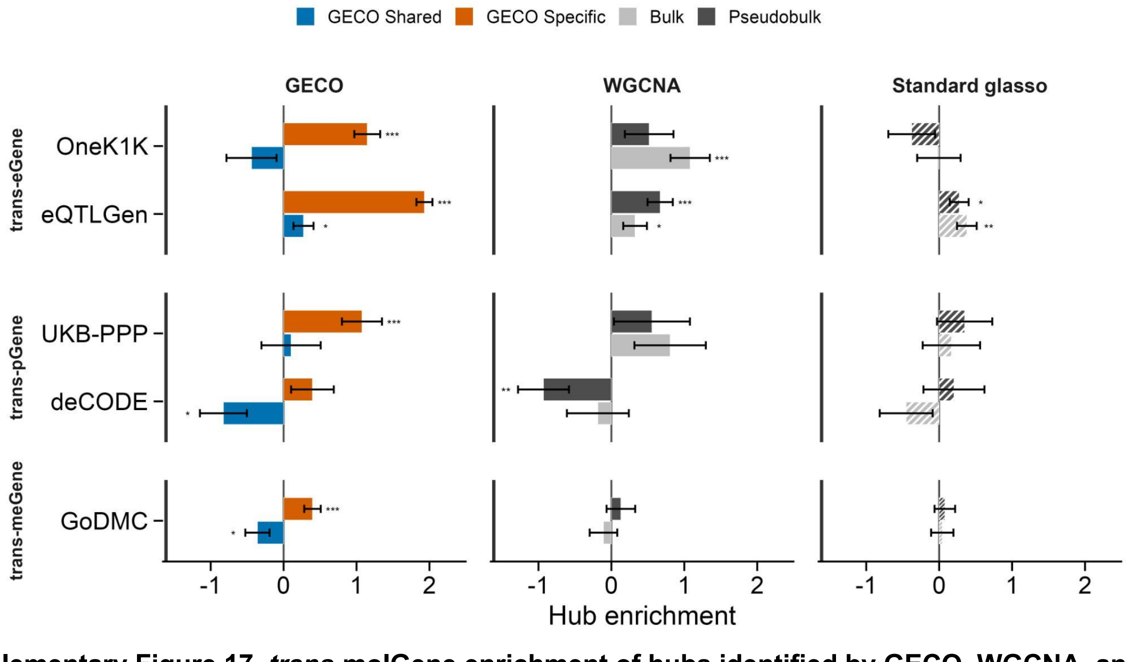


**Supplementary Figure 17. *trans*-molGene enrichment of hubs identified by GECO, WGCNA, and standard glasso in OneK1K.** ***: p < 0.001; **: p < 0.01; *: p < 0.05. Error bars indicate ±1 standard error. The x-axis shows log2 odds ratios (Methods).

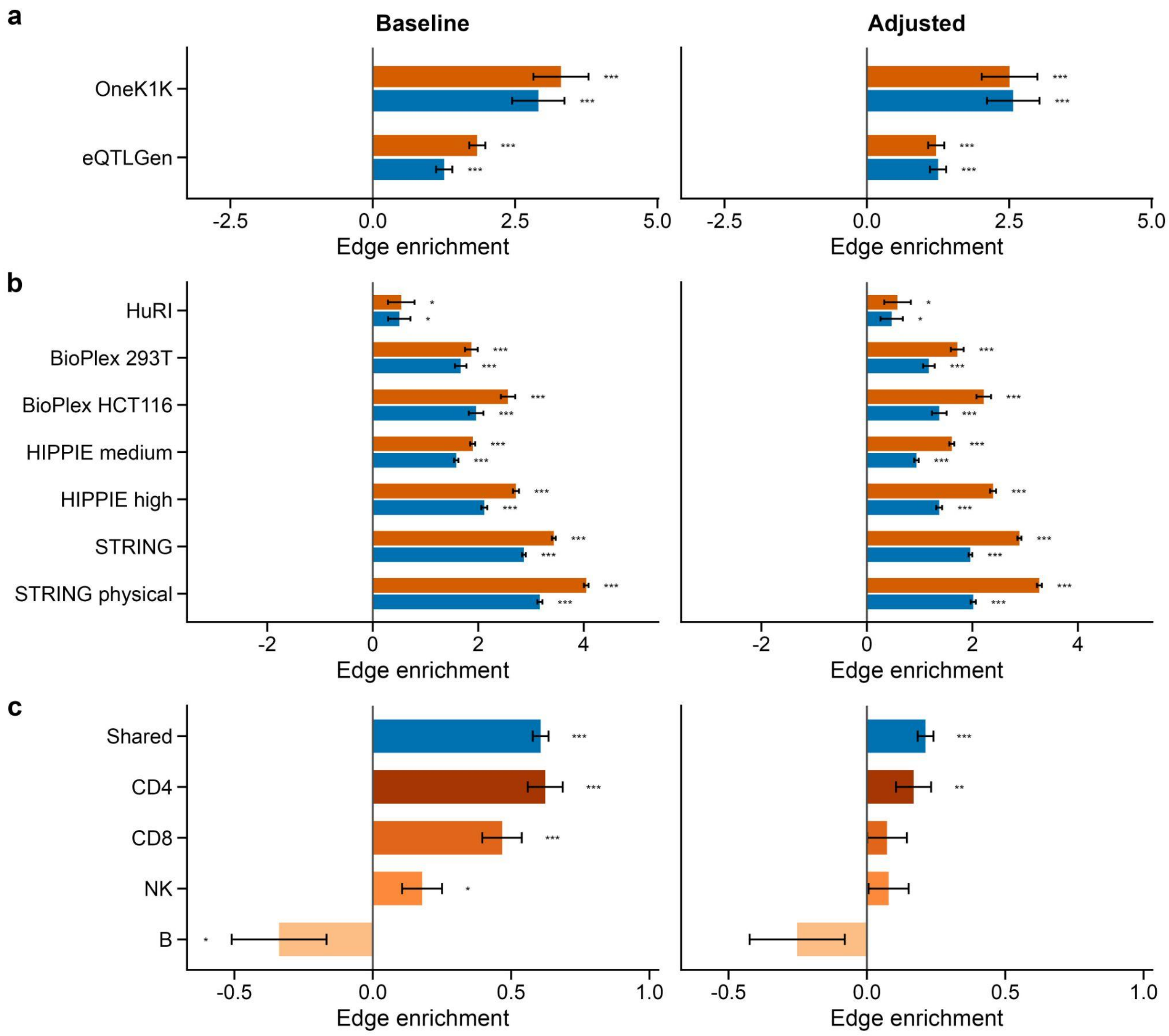


**Supplementary Figure 18. Primary and sensitivity analyses of *cis-trans* eGene pair (a), PPI pair (b), and perturbation-response pair from perturb-seq enrichment in OneK1K.** Baseline: primary analysis; Adjusted: sensitivity analysis controlling for mean expression, expression variability, gene length, and GC content. For each covariate, we adjusted for the mean value across the two genes in each pair. ***: p < 0.001; **: p < 0.01; *: p < 0.05. Error bars indicate ±1 standard error. The x-axis shows log2 odds ratios (Methods).

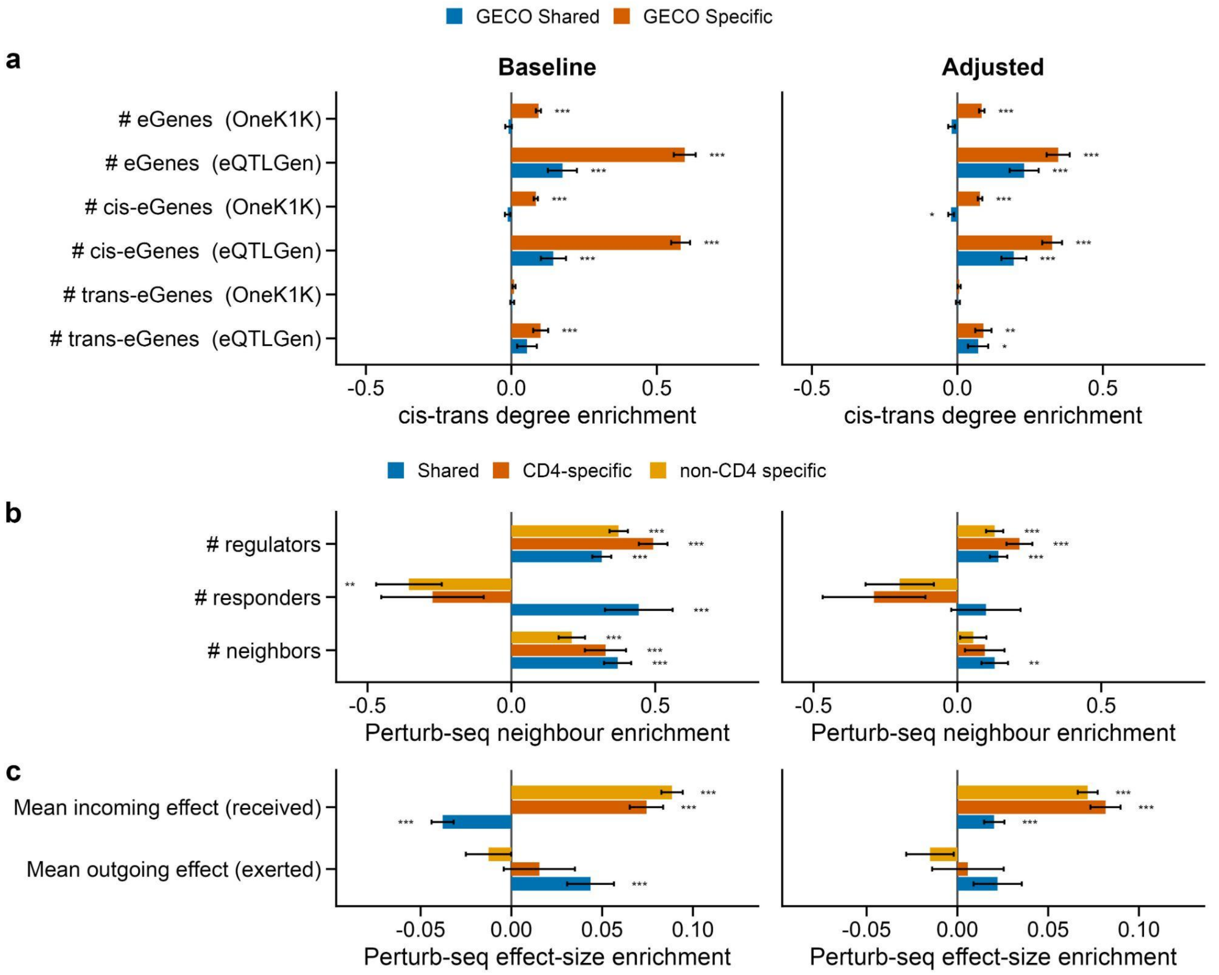


**Supplementary Figure 19. Hub positions in colocalized *cis-trans* eGene pairs (a) and perturbation-response pairs (b,c). (a)** Numbers of colocalized total, *cis*, and *trans* eGenes in the colocalized *cis-trans* eGene datasets. **(b)** Numbers of neighbors, regulators, and responders in the CD4 T cell Perturb-seq dataset. **(c)** Mean incoming and outgoing effects in the CD4 T cell Perturb-seq dataset. For genes whose knockdown produced no significant responders, the normalized effect size was set to 0. The x-axis shows regression coefficients (Methods).

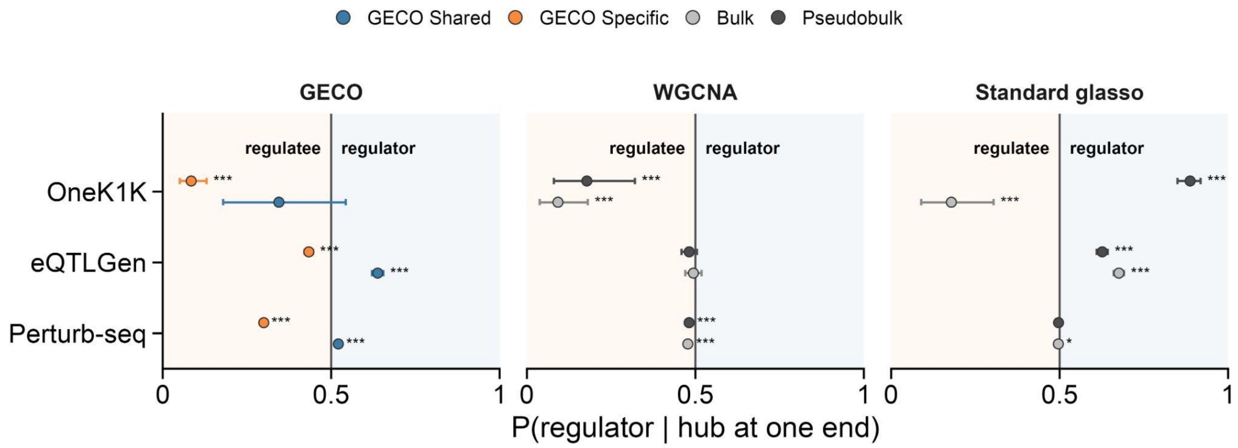


**Supplementary Figure 20. Network positions of hubs identified by GECO, WGCNA, and standard glasso.** Confidence intervals indicate 95% CIs from two-sided binomial tests.

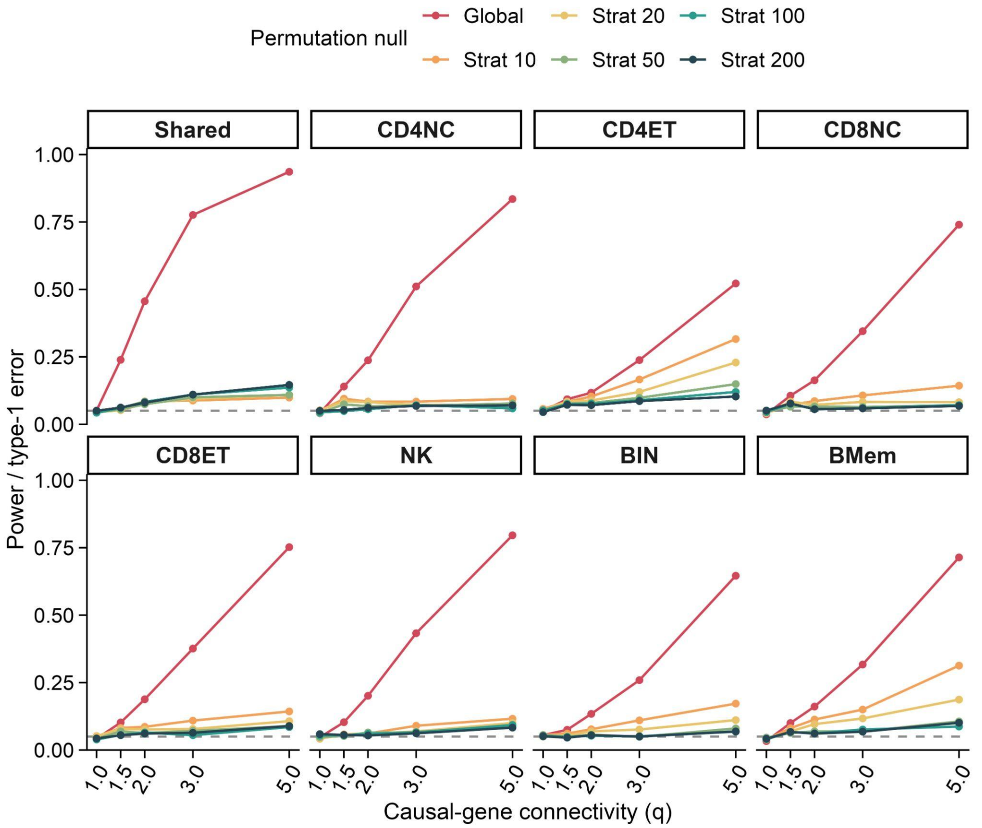


**Supplementary Figure 21. Simulation of networkMAGMA with binary weights.** Using a 0/1 adjacency matrix, networkMAGMA maintained nominal type I error (q = 1) in each permutation null setting across all eight GECO networks. As causal genes formed increasingly connected modules with larger q, power under the global null increased monotonically. Degree-stratified nulls were markedly conservative, remaining close to the 0.05 level even at high causal-gene connectivity. We therefore used the global null in real-data analyses.

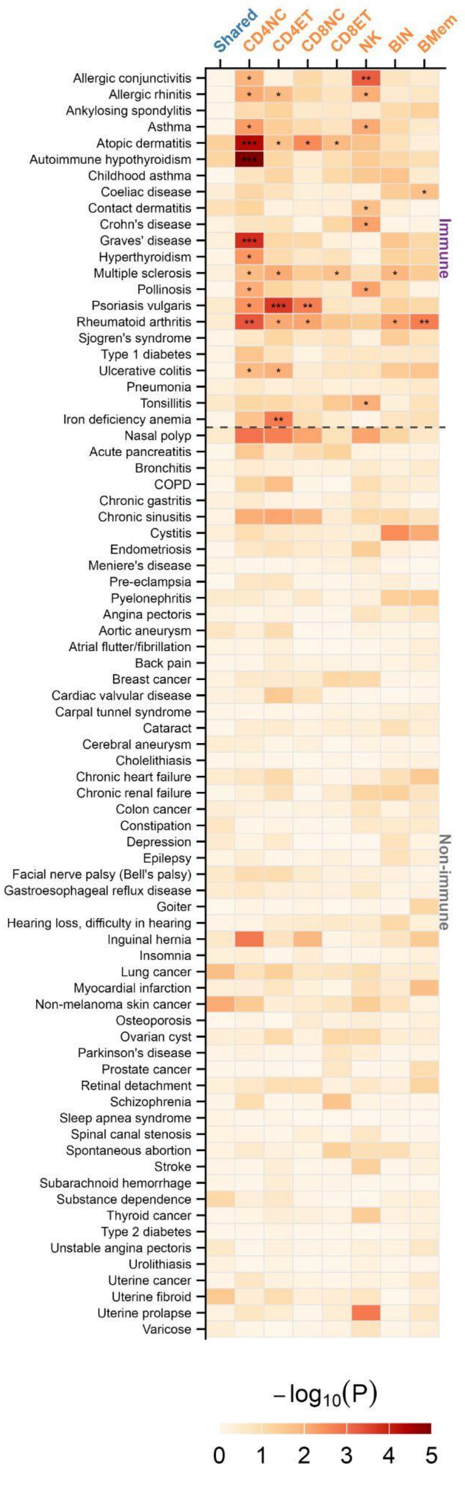


**Supplementary Figure 22. NetworkMAGMA results for each network-disease pair.** ***: q < 0.01; **: q < 0.05; *: q < 0.1. Multiple-testing correction was performed within each disease group.

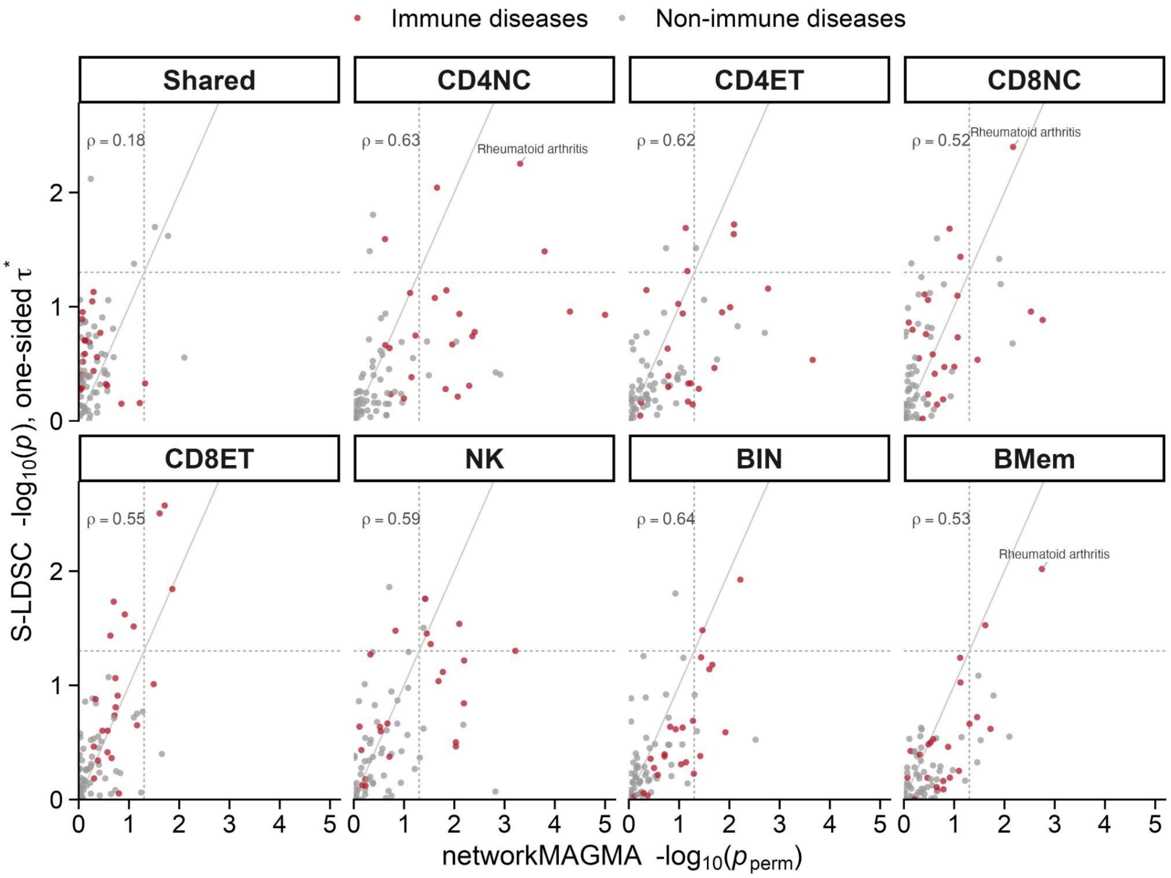


**Supplementary Figure 23. Concordance between networkMAGMA and S-LDSC cell type-disease enrichment signals.** Each panel shows one GECO network; each point is one of 78 FinnGen diseases. The x-axis shows the networkMAGMA GWAS gene pair enrichment strength; the y-axis shows the S-LDSC hub set enrichment strength. Points are colored by disease category. ρ, Spearman correlation between the two methods across the 78 diseases within each panel. The solid grey line is the identity (y = x); dashed lines mark nominal p = 0.05 on each axis. Disease-network pairs reaching p < 0.01 in both methods are labeled (rheumatoid arthritis in the CD4NC, CD8NC, and BMem networks).

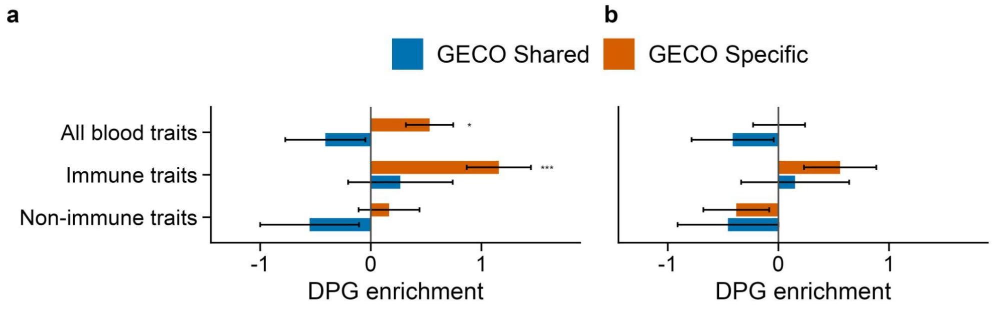


**Supplementary Figure 24. Primary and sensitivity analyses of DPG enrichment in OneK1K.** All blood traits were included in the union of immune and non-immune traits. **(a)** Baseline analysis. **(b)** Sensitivity analysis controlling for mean expression, expression variability, gene length, and GC content. ***: p < 0.001; **: p < 0.01; *: p < 0.05. Error bars show ±1 standard error, x-axes show log2 odds ratios.

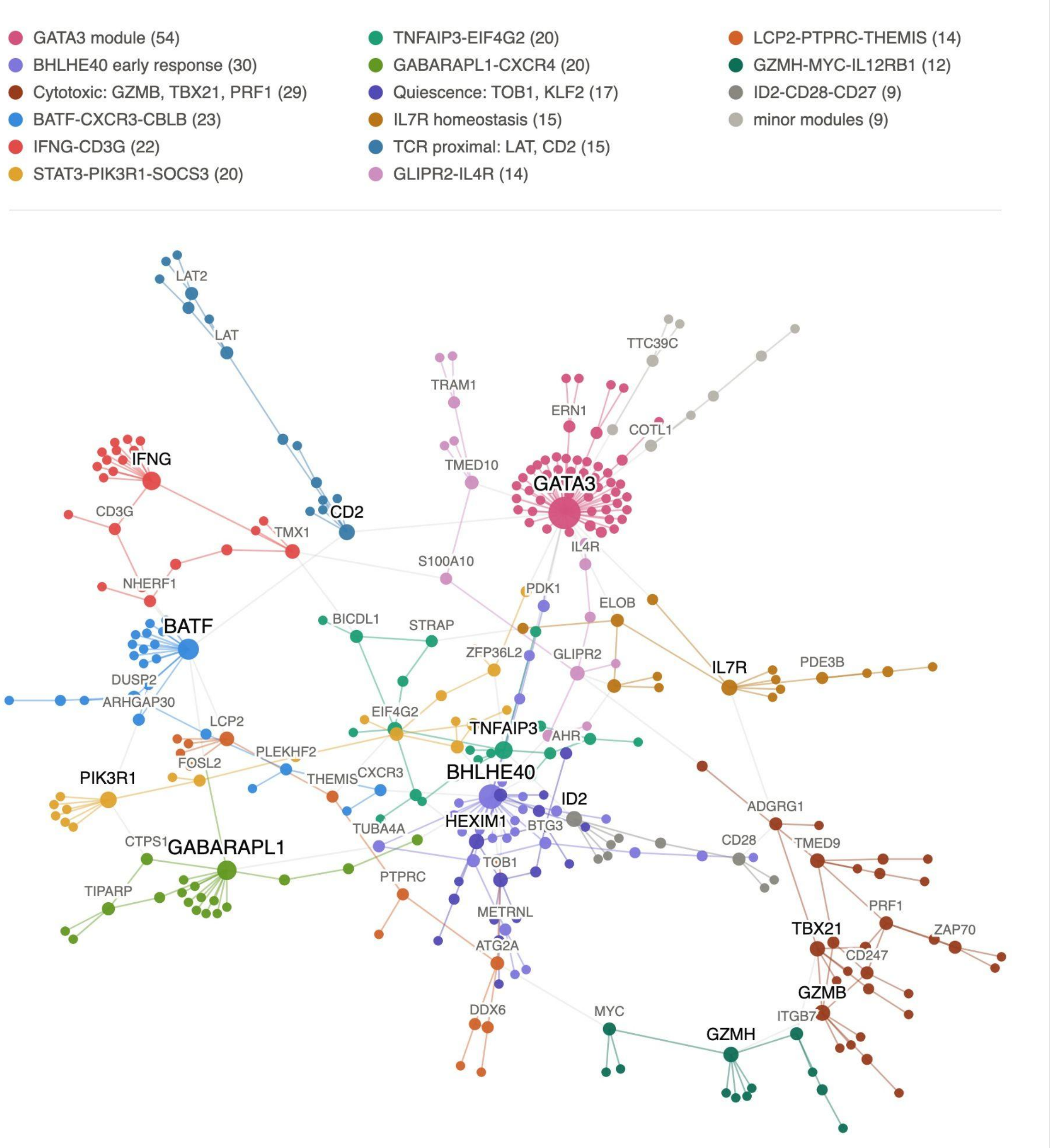


**Supplementary Figure 25. Module map of the GECO CD8 ∩ Perturb-seq overlap network.** Edges in both the GECO CD8-specific and CD4 T cell Perturb-seq networks were intersected, and the largest connected component is shown (323 genes, 370 edges). Nodes are genes positioned by a Fruchterman–Reingold force-directed layout; node color indicates Louvain community membership (17 modules, modularity Q = 0.79), with modules named by their highest-degree members and annotated functions (legend; module size in parentheses). Node and label sizes scale with degree, and only genes above a degree threshold are labeled. Within-module edges are colored by module; between-module edges are grey. The overlap is organized around cytotoxic-lineage programs, including a GZMB–TBX21–PRF1 cytotoxic module, IFNG–CD3G and LAT–CD2 TCR-associated modules, alongside a large GATA3 module and a TOB1–KLF2 quiescence module.

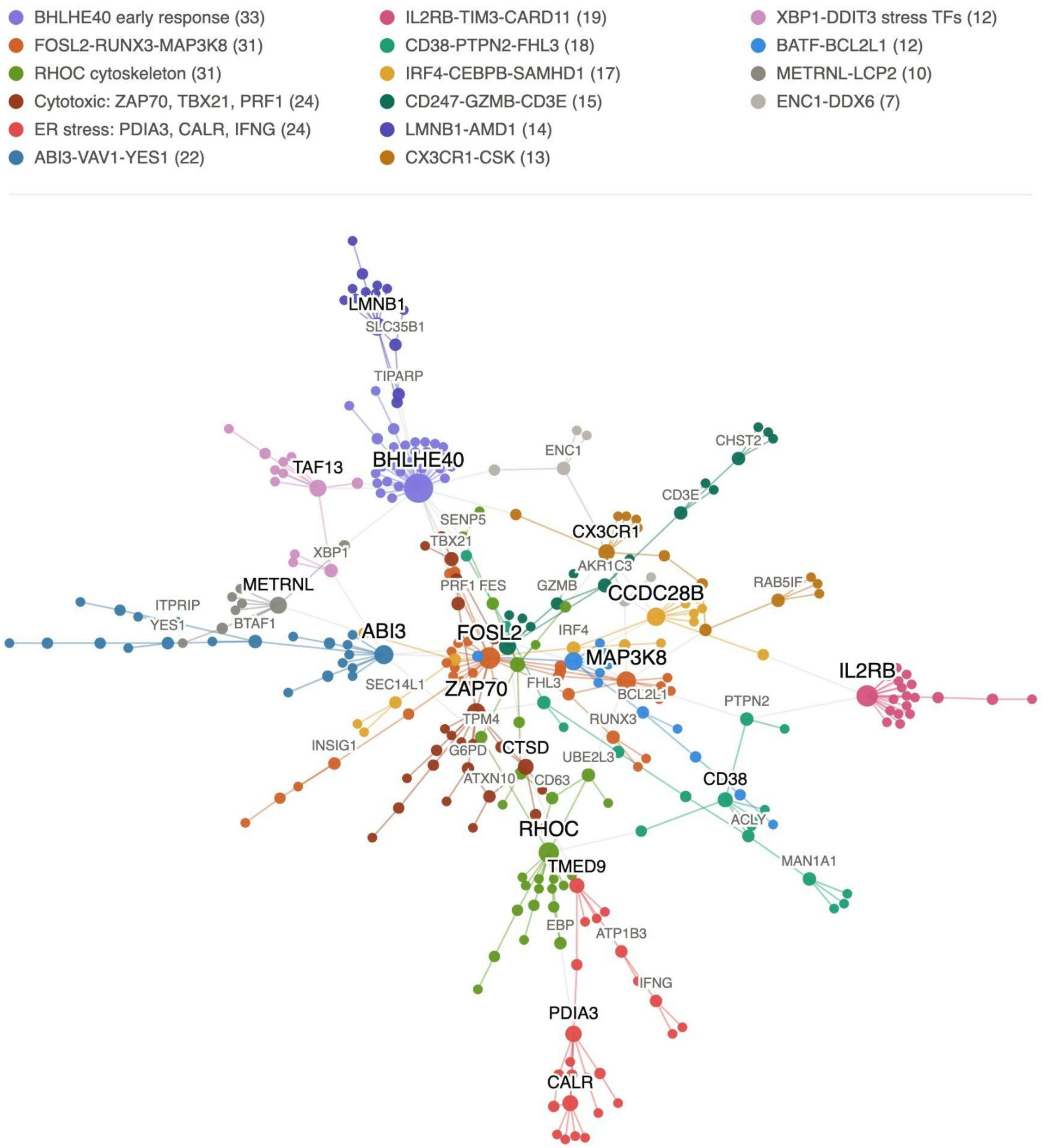


**Supplementary Figure 26. Module map of the GECO NK ∩ Perturb-seq overlap network.** Edges in both the GECO NK-specific and CD4 T cell Perturb-seq networks were intersected, and the largest connected component is shown (302 genes, 339 edges). Nodes are genes positioned by a Fruchterman–Reingold force-directed layout; node color indicates Louvain community membership (16 modules, modularity Q = 0.80), with modules named by their highest-degree members and annotated functions (legend; module size in parentheses). Node and label sizes scale with degree, and only genes above a degree threshold are labeled. Within-module edges are colored by module; between-module edges are grey. The overlap captures innate cytotoxicity and stress-response programs, including a ZAP70–TBX21–PRF1 cytotoxic module, IL2RB–TIM3–CARD11 receptor-signaling, ER-stress/UPR modules (PDIA3–CALR–IFNG; XBP1–DDIT3), and BHLHE40 and FOSL2–RUNX3 early-response modules.

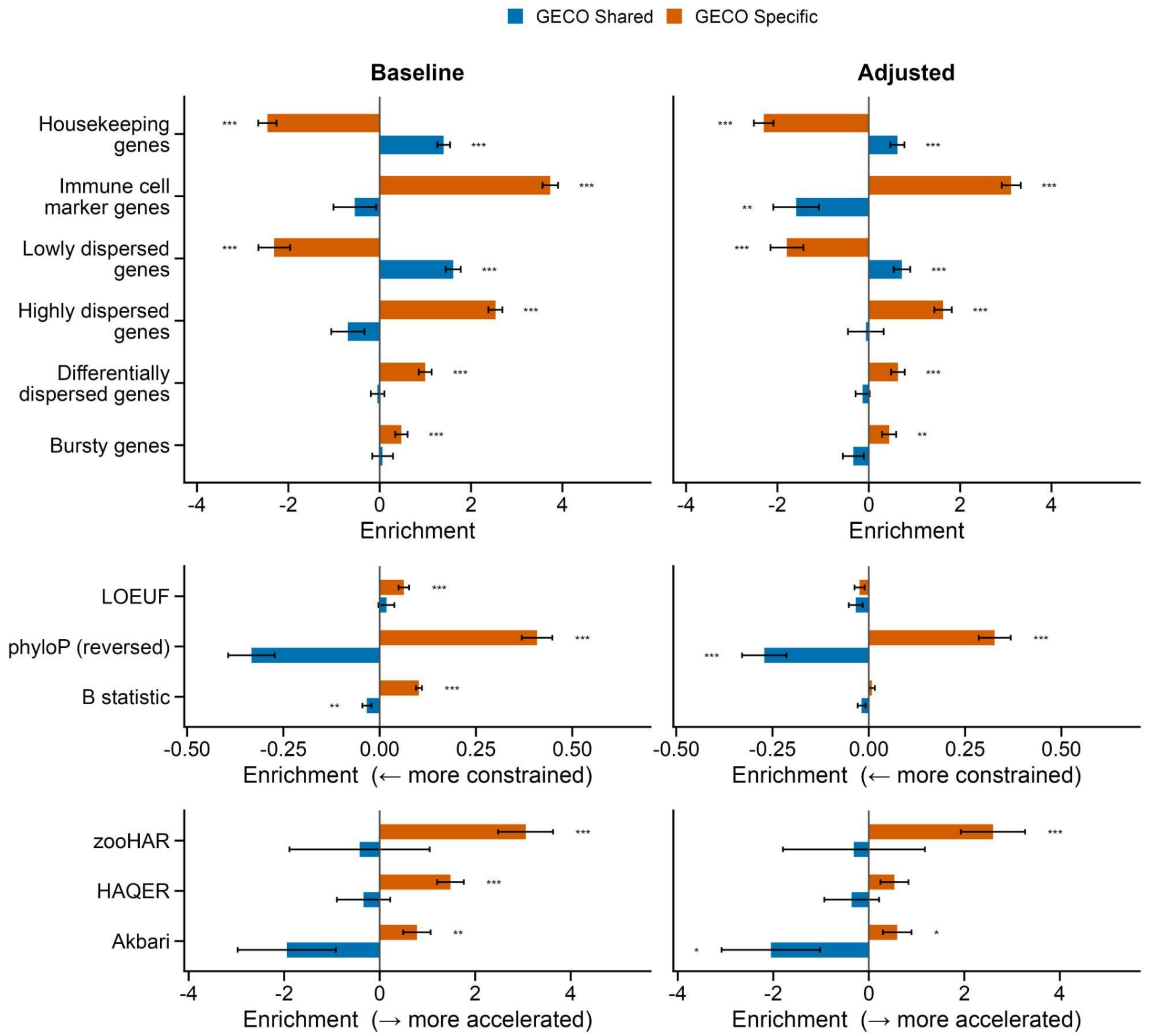


**Supplementary Figure 27. Primary and sensitivity analyses of benchmark gene set and evolutionary feature enrichment in CLUES/ImmVar.** Baseline: primary analysis; Adjusted: sensitivity analysis controlling for mean expression, expression variability, gene length, and GC content. For the six evolutionary features (LOEUF, phyloP, B statistic, zooHAR, HAQER, and Akbari), sensitivity analyses additionally adjusted for recombination rate and number of exons. ***: $p < 0.001$; **: $p < 0.01$; *: $p < 0.05$. Error bars indicate ±1 standard error. For binary annotations, the x-axis shows log2 odds ratios; for continuous annotations, it shows regression coefficients (Methods).

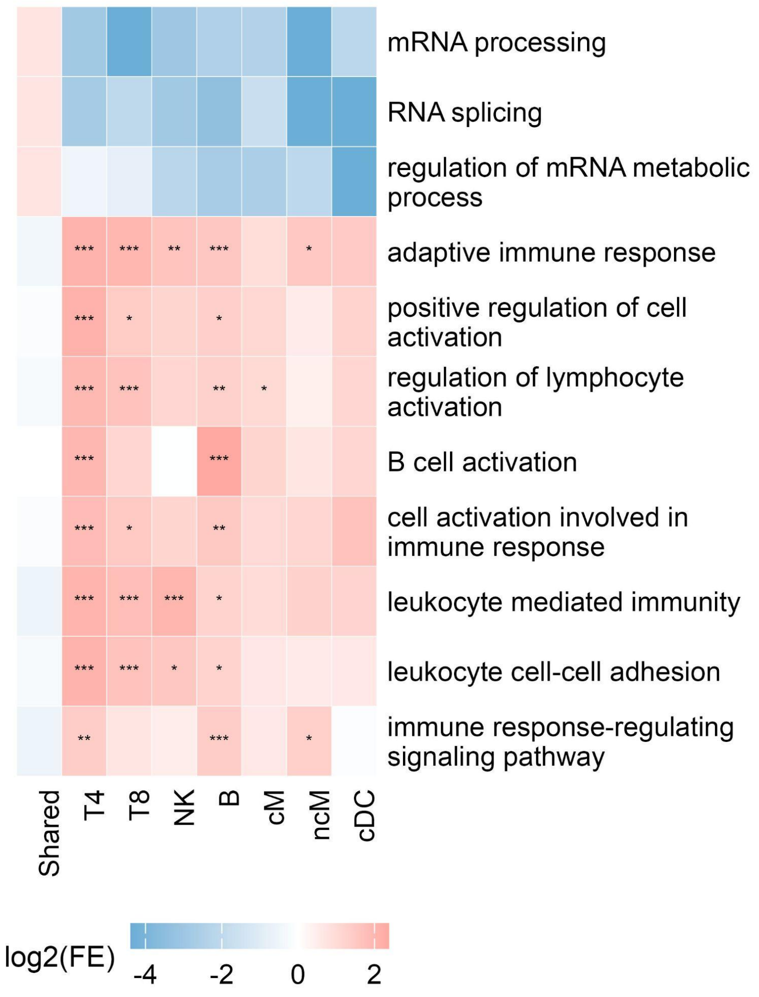


**Supplementary Figure 28. GO terms enrichment of shared and cell type-specific hubs in CLUES/ImmVar.** ***: FDR < 0.001; **: FDR < 0.01; *: FDR < 0.05.

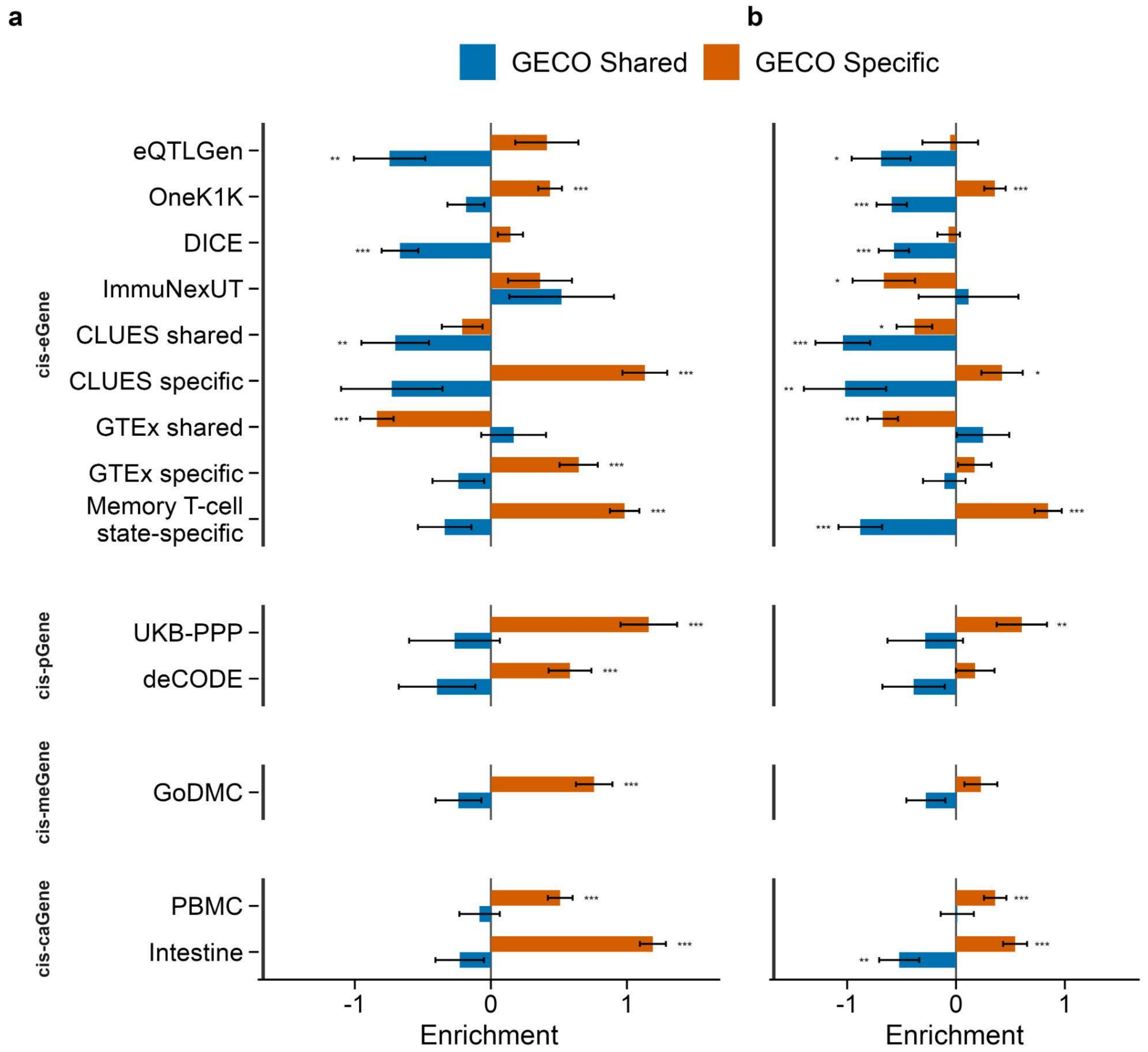


**Supplementary Figure 29. Primary and sensitivity analyses of *cis*-molGene enrichment in CLUES/ImmVar. (a)** Baseline analysis. **(b)** Sensitivity analysis controlling for mean expression, expression variability, gene length, and GC content. ***: $p < 0.001$; **: $p < 0.01$; *: $p < 0.05$. Error bars indicate ±1 standard error. The x-axis shows log2 odds ratios (Methods).

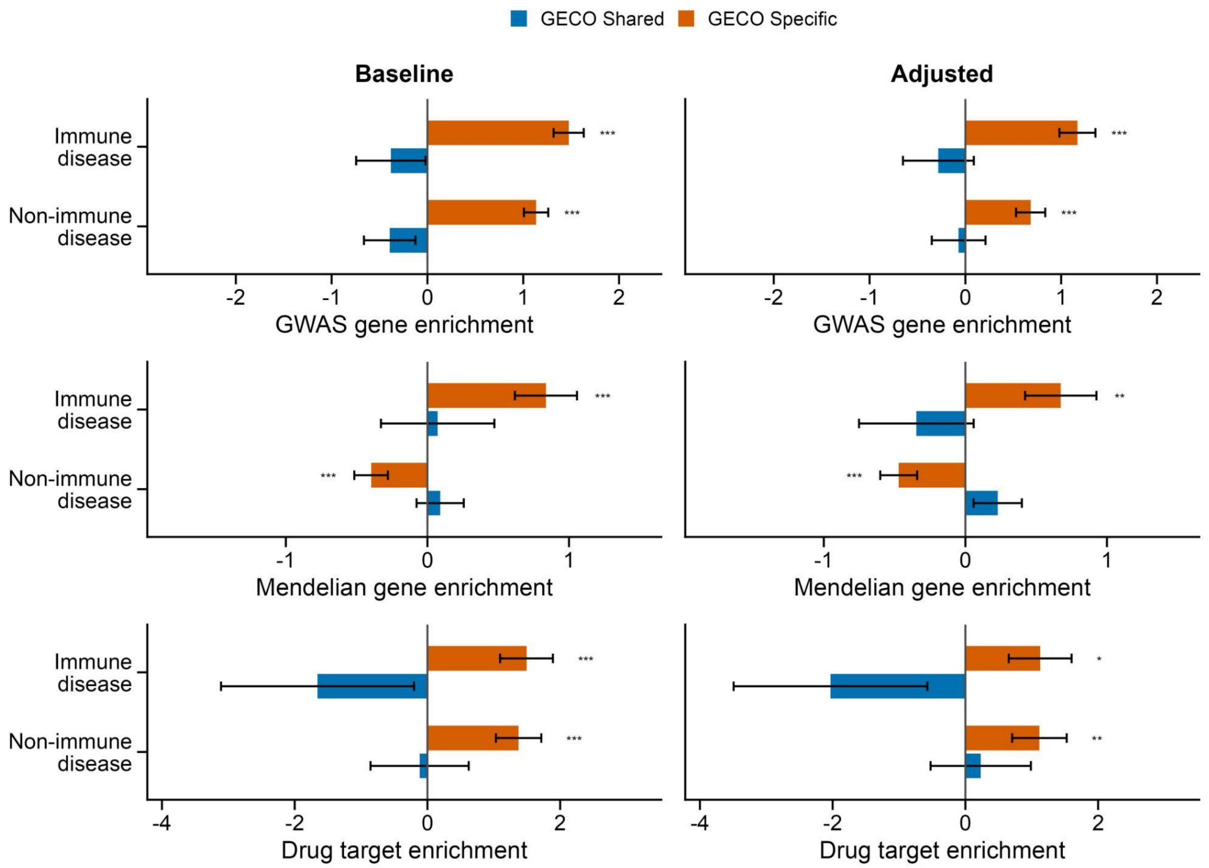


**Supplementary Figure 30. Primary and sensitivity analyses of GWAS gene, Mendelian gene, and approved drug target enrichment in CLUES/ImmVar.** Baseline: primary analysis; Adjusted: sensitivity analysis controlling for mean expression, expression variability, gene length, and GC content. ***: $p < 0.001$; **: $p < 0.01$; *: $p < 0.05$. Error bars indicate ±1 standard error, x-axes show log2 odds ratios.

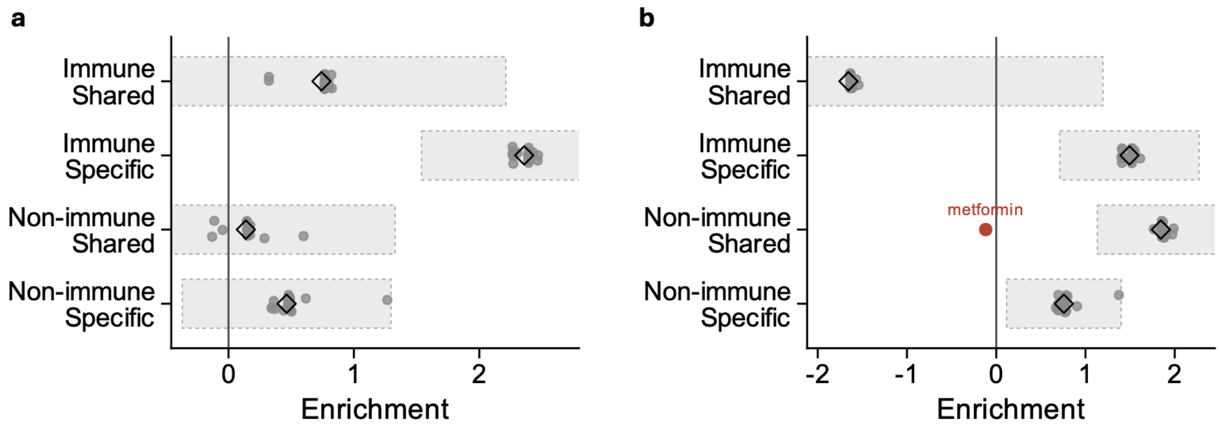


**Supplementary Figure 31. Leave-one-drug-out analysis in OneK1K (a) and CLUES/ImmVar (b).** Triangles indicate estimates based on all drugs, whereas grey dots represent leave-one-drug-out estimates. Shaded regions indicate the 95% CI of the full-data estimates. The x-axis shows log2 odds ratios (Methods). Metformin was excluded from the main CLUES/ImmVar analyses.

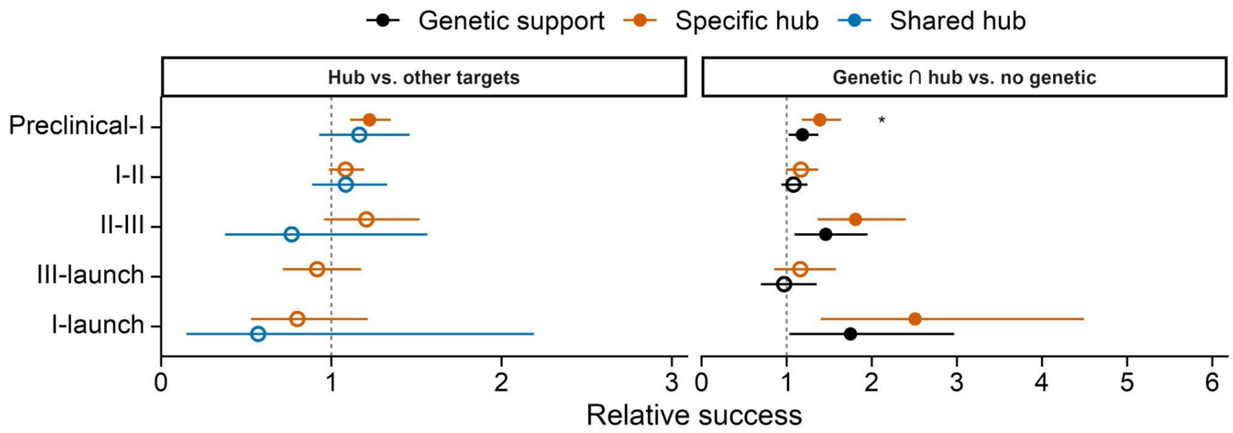


**Supplementary Figure 32. Relative success of target-indication pairs in immune indications from Citeline Pharmaprojects (CLUES/ImmVar).** Rows show progression of drug targets through phases. Panels show relative success for shared and specific hub genes (left) and for the intersection of specific hubs and indication-matched genetically supported genes as in Minikel et al. (OMIM or GWAS, right). These analyses restrict to genes included in GECO. Bars show 95% CI; filled points significantly differ from 1; and stars compare the orange and black estimates. ***: $p < 0.001$; **: $p < 0.01$; *: $p < 0.05$.

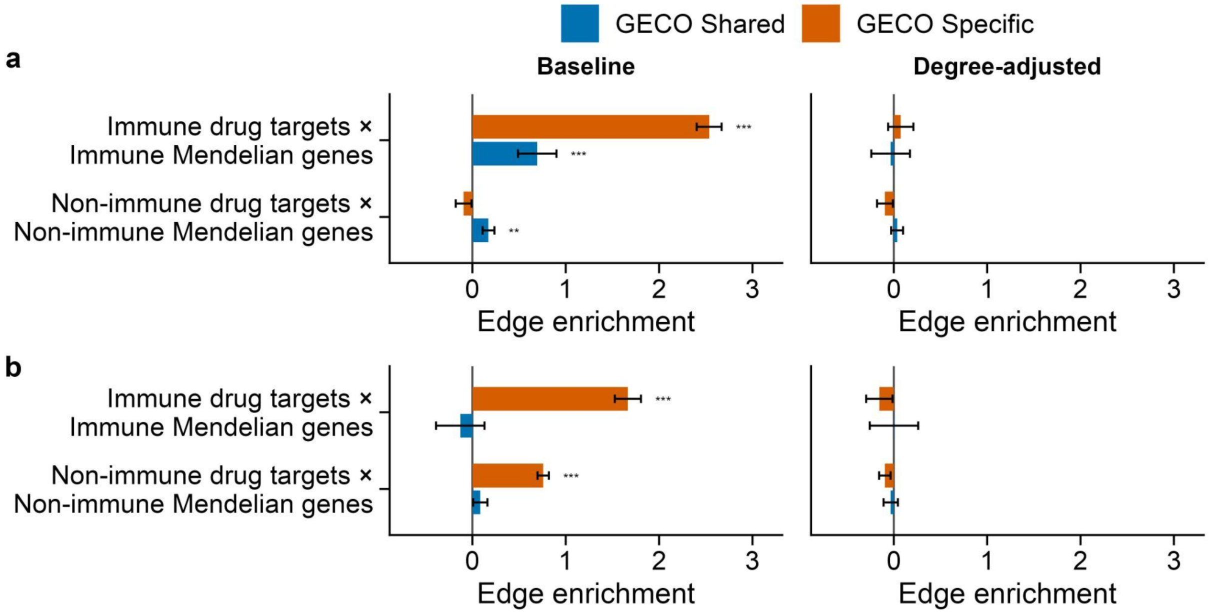


**Supplementary Figure 33. Drug target-Mendelian gene pair enrichment in OneK1K (a) and CLUES/ImmVar (b).** These enrichments become null after adjusting for network degree (degree was binned into 51 categories (zero plus 50 nonzero quantiles), averaged over the two endpoints), suggesting the primary signal is primarily driven by the fact that GWAS, Mendelian, and drug target genes are highly connected in appropriate cell type-specific networks. The x-axis shows log2 odds ratios (Methods).

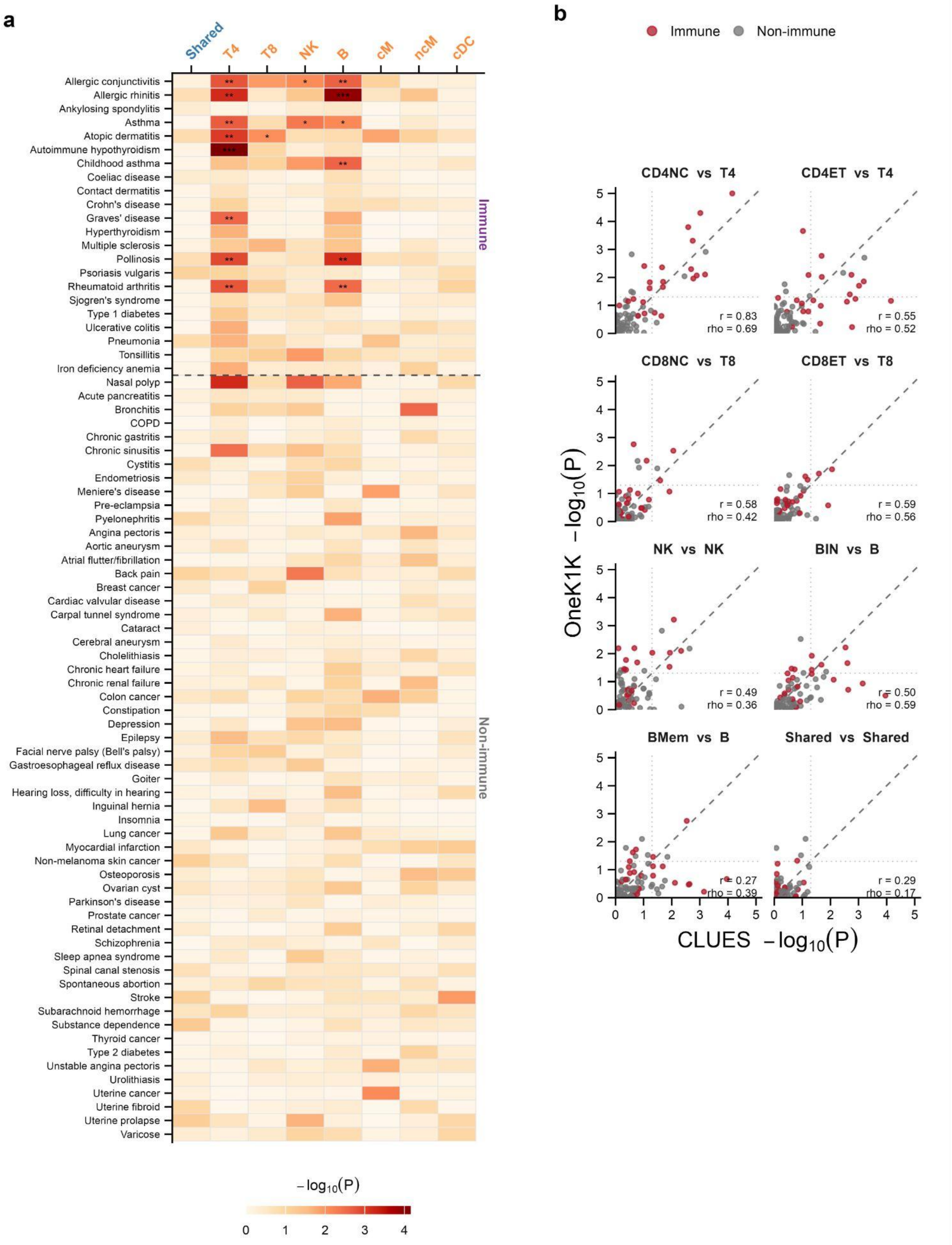


**Supplementary Figure 34. NetworkMAGMA results in CLUES/ImmVar. (a)** NetworkMAGMA results for each network–disease pair in CLUES/ImmVar. ***: q < 0.01; **: q < 0.05; *: q < 0.1. Multiple test correction was performed for immune and non-immune diseases. **(b)** Concordance of networkMAGMA results between OneK1K and CLUES/ImmVar for matched cell types. r and rho are the Pearson and Spearman correlation coefficients between the two cohorts, respectively.
