## supplementary note for "Cell type-specific coexpression networks funnel genetic risk into core genes"

### Model setting

Assume that there are $N$ individuals, $G$ genes and $C$ cell types. Let $y_{ijc}$ denote the pseudobulk expression of gene $j$ in individual $i$ and cell type $c$, and we assume $\sum_{i=1}^{N} y_{ijc} = 0$ for every gene $j$ and cell type $c$.

**Model.** The expression of gene $j$ in individual $i$ and cell type $c$ can be decomposed as

$$y_{ijc} = \alpha_{ij} + \Gamma_{ijc} + \delta_{ijc}\,,$$

where

- $\alpha_{ij}$ is the interindividual deviation shared across cell types, modelled as a random effect with $\alpha_i := (\alpha_{i1}, \ldots, \alpha_{iG})^T \sim N(0, V)$ independently across individuals, where $V$ is the $G \times G$ cell-type-shared covariance matrix;
- $\Gamma_{ijc}$ is the interindividual deviation specific to cell type $c$, modelled as a random effect with $vec(\Gamma_i) \sim N(0, diag(\Omega^1, \ldots, \Omega^C))$ independently across individuals, where $\Omega^c$ is the $G \times G$ cell-type-specific covariance matrix of cell type $c$ and $diag$ concatenates matrices block-wise (i.e., direct sum);
- $\delta_{ijc} := \frac{1}{n_{ic}} \sum_{s=1}^{n_{ic}} \epsilon_{ijcs}$ is the measurement noise arising from averaging a finite number of cells, modelled as a random effect with $\delta_{i \cdot c} \sim N(0, \Sigma_{ic}/n_{ic})$ independently across individuals and cell types, where $\Sigma_{ic}$ is the cell-to-cell covariance matrix of individual $i$ in cell type $c$ and $n_{ic}$ is the number of cells for individual *i* and cell type c.

The random effects $\alpha$, $\Gamma$ and $\delta$ are mutually independent. $\Sigma_{ic}$ is estimated by the within-individual sample covariance $\hat{\Sigma}_{ic} = \frac{1}{n_{ic}-1} \sum_{s=1}^{n_{ic}} (y_{i \cdot cs} - \bar{y}_{i \cdot c})(y_{i \cdot cs} - \bar{y}_{i \cdot c})^T$, where $y_{i \cdot cs}$ represents the expression in cell s of individual i and cell type c, and we write $v_{ijj'c} := (\hat{\Sigma}_{ic})_{jj'}/n_{ic}$.

Let $y \in \mathbb{R}^{NCG}$ be the vectorization of $Y \in \mathbb{R}^{N \times C \times G}$ with indices ordered lexicographically in $(i, c, j)$. Then

$$E(y) = 0\,, \qquad \mathbb{V}(y) = I_N \otimes J_C \otimes V + I_N \otimes \Omega + D\,,$$

where $J_C$ is the $C \times C$ matrix of ones, $\Omega := diag(\Omega^1, \ldots, \Omega^C)$, and $D := diag(\Sigma_{11}/n_{11}, \ldots, \Sigma_{1C}/n_{1C}, \ldots, \Sigma_{NC}/n_{NC})$. The parameters of interest are $V$ and $\Omega^1, \ldots, \Omega^C$; for a gene pair $(j, j')$ we write $\alpha_{jj'} := (V)_{jj'}$ and $g_{jj'}^c := (\Omega^c)_{jj'}$.

### Estimation

**Method of moments.** Let $S^c$ denote the $C \times C$ matrix whose $[c, c]$ entry is 1 and all other entries are 0, and $S^{jj'}$ the symmetric $G \times G$ matrix whose $[j, j']$ and $[j', j]$ entries are 1 and all other entries are 0 (so that $S^{jj}$ has a single nonzero entry). We estimate the variance components by the method of moments, i.e. as the minimizer of $\| yy^T - \mathbb{V}(y) \|_F^2$ over $V$

and $\Omega$. Setting the derivatives with respect to $\alpha_{jj'}$ and $g^c_{jj'}$ to zero gives the moment equations

$$tr\left((yy^T - \mathbb{V}(y))(I_N \otimes J_C \otimes S^{jj'})\right) = 0\,,$$

$$tr\left((yy^T - \mathbb{V}(y))(I_N \otimes S^c \otimes S^{jj'})\right) = 0\,, \quad c = 1, \dots, C\,.$$

The solution can be written as

$$\hat{\alpha}_{jj'} = \frac{tr\big((yy^T - D)\, Q_{1jj'}\big)}{tr(Q_{1jj'} Q^c_{2jj'})}\,, \qquad \hat{g}^c_{jj'} = \frac{tr\left((yy^T - D)\, Q^c_{2jj'}\right)}{tr(Q_{1jj'} Q^c_{2jj'})}\,,$$

with $Q_{1jj'} := I_N \otimes (I_C - J_C) \otimes S^{jj'}$ and $Q^c_{2jj'} := I_N \otimes [J_C - I_C - (C^2 - C)S^c] \otimes S^{jj'}$.

**Computation.** Expanding the traces, the estimators reduce to sums over individuals:

$$\hat{\alpha}_{jj'} = \frac{1}{N}\sum_{i=1}^{N} z_i\,, \qquad z_i := \frac{\big(\sum_{c=1}^{C} y_{ijc}\big)\big(\sum_{c=1}^{C} y_{ij'c}\big) - \sum_{c=1}^{C} y_{ijc}\ y_{ij'c}}{C(C-1)} = \frac{\sum_{c\neq c'} y_{ijc}\ y_{ij'c'}}{C(C-1)}\,,$$

$$\hat{g}^c_{jj'} = \frac{1}{N}\sum_{i=1}^{N} w^c_i\,, \qquad w^c_i := y_{ijc}\, y_{ij'c} - v_{ijj'c} - z_i\,.$$

Each gene pair requires $O(NC)$ operations, and the per-individual terms $z_i$ and $w^c_i$ are reused for the standard errors below.

### Theoretical properties of the estimator

**Unbiasedness.** For any matrix $Q$, $E(tr(yy^T Q)) = E(y^T Q y) = tr(Q\, \mathbb{V}(y))$. Using

$$tr\big(Q_{1jj'}(I_N \otimes J_C \otimes V)\big) = N\, tr((I_C - J_C)J_C) tr(S^{jj'} V) = q_{jj'}\, \alpha_{jj'}\,,$$

$$tr\big(Q_{1jj'}(I_N \otimes \Omega)\big) = N \sum_{c=1}^{C} (I_C - J_C)_{cc}\, tr(S^{jj'} \Omega^c) = 0\,,$$

where $q_{jj'} := tr(Q_{1jj'} Q^c_{2jj'}) = -NC(C-1)$ for $j = j'$ and $-2NC(C-1)$ for $j \neq j'$, we obtain

$$E\big(\hat{\alpha}_{jj'}\big) = \frac{tr\big(Q_{1jj'} \mathbb{V}(y)\big) - tr(DQ_{1jj'})}{q_{jj'}} = \frac{q_{jj'}\, \alpha_{jj'} + tr(Q_{1jj'} D) - tr(DQ_{1jj'})}{q_{jj'}} = \alpha_{jj'}\,.$$

Similarly, $tr\left(Q^c_{2jj'}(I_N \otimes J_C \otimes V)\right) = N\, tr((J_C - I_C - (C^2 - C)S^c)J_C) tr(S^{jj'} V) = 0$ and $tr\left(Q^c_{2jj'}(I_N \otimes \Omega)\right) = -N(C^2 - C)\, tr(S^{jj'} \Omega^c) = q_{jj'}\, g^c_{jj'}$, so that $E\left(\hat{g}^c_{jj'}\right) = g^c_{jj'}$. Both estimators are therefore unbiased.

**Asymptotic normality and standard errors.** $z_i$ and $w_i^c$ are independent across $i$ with finite variances, and by the central limit theorem

$$\sqrt{N}\left(\hat{\alpha}_{jj'} - \alpha_{jj'}\right) \xrightarrow{d} N(0, \sigma_z^2), \qquad \sqrt{N}\left(\hat{g}_{jj'}^c - g_{jj'}^c\right) \xrightarrow{d} N(0, \sigma_{w^c}^2),$$

where $\sigma_z^2$ and $\sigma_{w^c}^2$ are consistently estimated by the sample variances $\hat{\sigma}_z^2$ and $\hat{\sigma}_{w^c}^2$ of $\{z_i\}$ and $\{w_i^c\}$. The standard errors of $\hat{\alpha}_{jj'}$ and $\hat{g}_{jj'}^c$ are therefore $\hat{\sigma}_z/\sqrt{N}$ and $\hat{\sigma}_{w^c}/\sqrt{N}$, which are asymptotically equivalent to a delete-one jackknife over individuals as in our prior work (Chen 2024 Nat Comm) but without the N-fold additional computation from jackknife.

**Effect of centering.** In the above section, we assume $\sum_{i=1}^{N} y_{ijc} = 0$ for every gene $j$ and cell type $c$. This is a common practice for moment-based estimators [Wu & Sankararaman 2018 Bioinformatics, Yang 2017 Nat Genet] and here we show the effect can be computed exactly and is negligible for the sample sizes considered. Let $A \in \mathbb{R}^{N\times K}$ be the covariate matrix, including an intercept column, $P := A(A^TA)^{-1}A^T$ the corresponding projection matrix, and $p_i := P_{ii}$ the leverage of individual $i$ (with $\sum_i p_i = K$), $M := I_N \otimes I_C \otimes I_G - \frac{1}{N}J_N \otimes \left(I_C - \frac{1}{C}J_C\right) \otimes I_G - P \otimes \frac{1}{C}J_C \otimes I_G$. Let $v_{ijj'c}$ denote the true noise covariance $(\Sigma_{ic})_{jj'}/n_{ic}$ and $d_i := \sum_{c=1}^{C} v_{ijj'c}$. Let $\tilde{y}$ denote the pseudobulk expression before centering, so that $y = M\tilde{y}$. Since $E(y) = 0$, for any matrix $Q$ we have $E(\mathrm{tr}(yy^TQ)) = \mathrm{tr}(Q\,M\,\mathbb{V}(\tilde{y})\,M)$, and a direct calculation gives

$$E\left(\hat{\alpha}_{jj'}\right) = \frac{N-K}{N}\,\alpha_{jj'} - \frac{K-1}{NC^2}\sum_{c=1}^{C} g_{jj'}^c - \frac{1}{NC^2}\sum_{i=1}^{N}\left(p_i - \frac{1}{N}\right)d_i\,,$$

$$E\left(\hat{g}_{jj'}^c\right) = \frac{N-1-2(K-1)/C}{N}\,g_{jj'}^c + \frac{2(K-1)}{NC^2}\sum_{c'=1}^{C} g_{jj'}^{c'} - \frac{1}{N^2}\sum_{i=1}^{N} v_{ijj'c} + \frac{2}{N}\sum_{i=1}^{N}\left(p_i - \frac{1}{N}\right)\left(\frac{1}{C^2}d_i - \frac{1}{C}v_{ijj'c}\right).$$

Apart from the leading scaling factors, all terms are of order $K/N$. The effect therefore vanishes with large N.
